# Identification of key genes and associated pathways in neuroendocrine tumors through bioinformatics analysis and predictions of small drug molecules

**DOI:** 10.1101/2020.12.23.424165

**Authors:** Praveenkumar Devarbhavi, Basavaraj Vastrad, Anandkumar Tengli, Chanabasayya Vastrad, Iranna Kotturshetti

## Abstract

Neuroendocrine tumor (NET) is one of malignant cancer and is identified with high morbidity and mortality rates around the world. With indigent clinical outcomes, potential biomarkers for diagnosis, prognosis and drug target are crucial to explore. The aim of this study is to examine the gene expression module of NET and to identify potential diagnostic and prognostic biomarkers as well as to find out new drug target. The differentially expressed genes (DEGs) identified from GSE65286 dataset was used for pathway enrichment analyses and gene ontology (GO) enrichment analyses and protein - protein interaction (PPI) analysis and module analysis. Moreover, miRNAs and transcription factors (TFs) that regulated the up and down regulated genes were predicted. Furthermore, validation of hub genes was performed. Finally, molecular docking studies were performed. DEGs were identified, including 453 down regulated and 459 up regulated genes. Pathway and GO enrichment analysis revealed that DEGs were enriched in sucrose degradation, creatine biosynthesis, anion transport and modulation of chemical synaptic transmission. Important hub genes and target genes were identified through PPI network, modules, target gene - miRNA network and target gene - TF network. Finally, survival analyses, receiver operating characteristic (ROC) curve and RT-PCR validated the significant difference of ATP1A1, LGALS3, LDHA, SYK, VDR, OBSL1, KRT40, WWOX, NINL and PPP2R2B between metastatic NET and normal controls. In conclusion, the DEGs and hub genes with their regulatory elements identified in this study will help us understand the molecular mechanisms underlying NET and provide candidate targets for future research.

## Introduction

A small intestine neuroendocrine tumors (NET) is common cancer of small bowel [1]. The absence of early diagnostic and biomarkers are the main reasons for death caused by NET. The 5 year survival rate across all stages of NET is only 70-100% [2]. In the past several decades, numerous attempts have been taken to disclose the molecular pathogenesis of NET, and to advance the patient prognosis through many therapeutic strategies. However, narrow progress has been made to lengthen the survival and to diminish the mortality. For patients, surgery, trans arterial embolisation and chemotherapy offer only limited options for NET treatment [3–4]. However, due to surgical therapy limitations, progressive tumors can reappear even after complete surgical resection; maximum patients with NET are not suitable for surgical resection and fewer than 23.8% of patients respond to conventional chemotherapy [5]. In addition, the clinical success of NET always compromised due to initial metastasis and chemo resistance. Consequently, extensive investigations of potential diagnosis biomarkers and identification of new therapeutic targets for NET patients are urgently needed.

Recently, the gene expression profile chip, a high throughput and efficient technique, has been extensively used in an array of disease research fields to explain the association between disease and genes and pathways, and provide the important clues for the progression of the NET [6]. For example, alteration in CDKN1B was linked with development of NET [7]. Genes such as NAP1L1, MAGE-D2, MTA1, MIB-1, p53, and bcl-2 were responsible for pathogenesis of NET [8–9]. High expression of CDX-2 was liable for development of NET [10]. Loss of tumor suppressor gene TCEB3C was linked with development of NET [11]. Pathways such as mTOR signaling pathway [12], notch signaling [13], phosphatidylinositol 3-kinase/Akt signaling [14], sonic hedgehog-gli1 signaling pathway [15] and GPCR pathways [16] were responsible for pathogenesis of NET. Thus, more work is needed to discover the underlying molecular mechanisms in NET.

In this study, we diagnosed important genes in NET by combining bioinformatics analyses. In this study, we diagnosed important genes in NET by combining bioinformatics analyses. Differentially expressed genes (DEGs), pathways, and gene ontology (GO) terms associated with NET were investigated. Subsequently, the identification of hub genes from protein–protein interactions (PPIs) network, modules analysis, construction of target gene - miRNA network and target gene - TF network. Furthermore, survival analyses, receiver operating characteristic (ROC) curve and RT-PCR were carried out for validation of hub genes. Finally, molecular docking studies was performed. We investigated the potential candidate biomarkers for their utility in diagnosis, prognosis, and drug targeting in NET.

## Materials and methods

### Microarray data

Gene expression profile GSE65286 was downloaded from the National Center for Biotechnology Information Gene Expression Omnibus (GEO, http://www.ncbi.nlm.nih.gov/geo) [17] database. GSE65286 included 33 patients with metastatic NET and on 10 normal controls. The expression profile was provided on GPL4133 Agilent-014850 Whole Human Genome Microarray 4x44K G4112F (Feature Number version).

### DEG screening

The original txt files were downloaded and classified as patients with metastatic NET and normal controls. The limma package of Bioconductor (http://www.bioconductor.org/) [18] was used for data standardization and transforming raw data into expression values. The significance analysis of the empirical Bayes methods within limma package was applied to diagnose DEGs between patients with metastatic NET and on normal controls. The genes with the following cutoff criteria were considered as the significant DEGs: p < 0.05, and |logFC|> 3.01 for up regulated genes and |logFC|> −2.75 for down regulated genes.

### Pathway enrichment analysis

Pathway enrichment analysis was used to diagnose the potential functional and metabolic pathways associated with DEGs. BIOCYC (https://biocyc.org/) [19], Kyoto Encyclopedia of Genes and Genomes (KEGG) (http://www.genome.jp/kegg/pathway.html) [20], Pathway Interaction Database (PID) (https://wiki.nci.nih.gov/pages/viewpage.action?pageId=315491760) [21], REACTOME (https://reactome.org/) [22], GenMAPP (http://www.genmapp.org/) [23], MSigDB C2 BIOCARTA (v6.0) (http://software.broadinstitute.org/gsea/msigdb/collections.jsp) [24], PantherDB (http://www.pantherdb.org/) [25], Pathway Ontology (http://www.obofoundry.org/ontology/pw.html) [26] and Small Molecule Pathway Database (SMPDB) (http://smpdb.ca/) [27] were a collection of databases that store a large number of information about genomes, biological pathways, diseases, chemical substances, and drugs. We performed pathway enrichment analysis by ToppCluster (https://toppcluster.cchmc.org/) [28]. ToppCluster is a commonly used online biological information database that provides comprehensive pathway interpretations. p < 0.05 was considered statistically significant.

### Gene ontology (GO) enrichment analysis

The GO (http://www.geneontology.org) [29] database associated three categories such as biological process (BP), cellular component (CC), and molecular function (MF). ToppCluster (https://toppcluster.cchmc.org/) [28] provides a set of functional annotation tools to analyze the biological roles of genes. In this study, GO terms were analyzed using the DAVID online tool with the enrichment threshold of P<0.05.

### Protein–protein interaction (PPI) network construction and module analysis

Human Integrated Protein-Protein Interaction rEference (HIPPIE) (http://cbdm.uni-mainz.de/hippie/) [30] was used to construct the PPI network which integrates different PPI databases such as IntAct (https://www.ebi.ac.uk/intact/) [31], BioGRID (https://thebiogrid.org/) [32], HPRD (http://www.hprd.org/) [33], MINT (https://mint.bio.uniroma2.it/) [34], BIND (http://download.baderlab.org/BINDTranslation/) [35], MIPS (http://mips.helmholtz-muenchen.de/proj/ppi/) [36] and DIP (http://dip.doe-mbi.ucla.edu/dip/Main.cgi) [37]. The interactions with a combined score >0.4 were considered significant. The PPI network was visualized through the Cytoscape software (http://www.cytoscape.org/) [38]. Five topological properties such as node degree (number of interactions with author genes) [39], betweenness (fraction of shortest paths between node pairs in a network) [40], stress (node is a node traversed by a high number of shortest paths) [41], closeness (inverse of the average length of the shortest paths to/from all the other nodes) [42] and clustering coefficient (measure of degree to which nodes in a graph tend to cluster together) [43] were implemented for PPI network analysis. PEWCC1 plug in of the Cytoscape software was applied to screen significant modules from the PPI network [44]. The degree cutoff = 10, node score cutoff = 0.2, k-core = 2, and max. depth = 100 were used as selection criterion.

### Construction of the target gene - miRNA network

NetworkAnalyst (https://www.networkanalyst.ca/) [45] was identify microRNAs (miRNAs) around which significant changes occur at the transcriptional level, we employed the comprehensive human post-transcriptional regulatory network consisting of the experimentally verified target gene miRNA network from TarBase (http://diana.imis.athena-innovation.gr/DianaTools/index.php?r=tarbase/index) [46] and miRTarBase (http://mirtarbase.mbc.nctu.edu.tw/php/download.php) [47] databases. The target gene miRNA network was visualized using Cytoscape software (http://www.cytoscape.org/) [38].

### Construction of the target gene - TF network

NetworkAnalyst (https://www.networkanalyst.ca/) [45] was identify transcription factors (TFs) around which significant changes occur at the transcriptional level, we employed the comprehensive human transcriptional regulatory network consisting of the experimentally verified target gene TF network from ChEA (http://amp.pharm.mssm.edu/lib/chea.jsp) [48] database. The target gene TF network was visualized using Cytoscape software (http://www.cytoscape.org/) [38].

### Validations of hub genes

PROGgeneV2 (http://genomics.jefferson.edu/proggene/) [49] was used to perform a survival analysis of the core genes. The hazard ratio (HR) with 95% confidence intervals and log-rank P value were calculated and displayed on the plot. The data of PROGgeneV2 are based on GEO and the Cancer Genome Atlas database. The receiver operating characteristic (ROC) curve was plotted and the area under the curve (AUC) was calculated by using the generalized linear model (GLM) in machine learning algorithms with R package“pROC” to evaluate the capability of genetic markers in distinguishing metastatic NET and normal controls using GEO database [50]. Finally, reverse transcription polymerase chain reaction (RT-PCR) was carried out for validation of hub genes. Total RNA from in vitro cultured cells of NET (ATCC® CRL-3254) and normal (ATCC® CRL-7869) was isolated with TRI Reagent® (Sigma, USA) according to the manufacturer’s protocol. According protocol for cDNA synthesis was reverse transcription at 42°C for 75 min and at 98°C for 5 min by using Reverse transcription cDNA kit (Thermo Fisher Scientific, Waltham, MA, USA). PCR amplification was performed using the QuantStudio 7 Flex real-time PCR system (Thermo Fisher Scientific, Waltham, MA, USA). The RT -PCR thermocycling conditions were as follows: initial denaturation at 95°C for 30 sec and 40 cycles of 95°C for 5 sec and 60°C for 34 sec. The relative expression level was determined as targeting genes divided by β in. Relative miRNA expression was generated with 2^−ΔΔCt^ method [51]. Primers used in this study are shown in Table 1. All experiments were performed independently in triplicate.

**Table 1.**
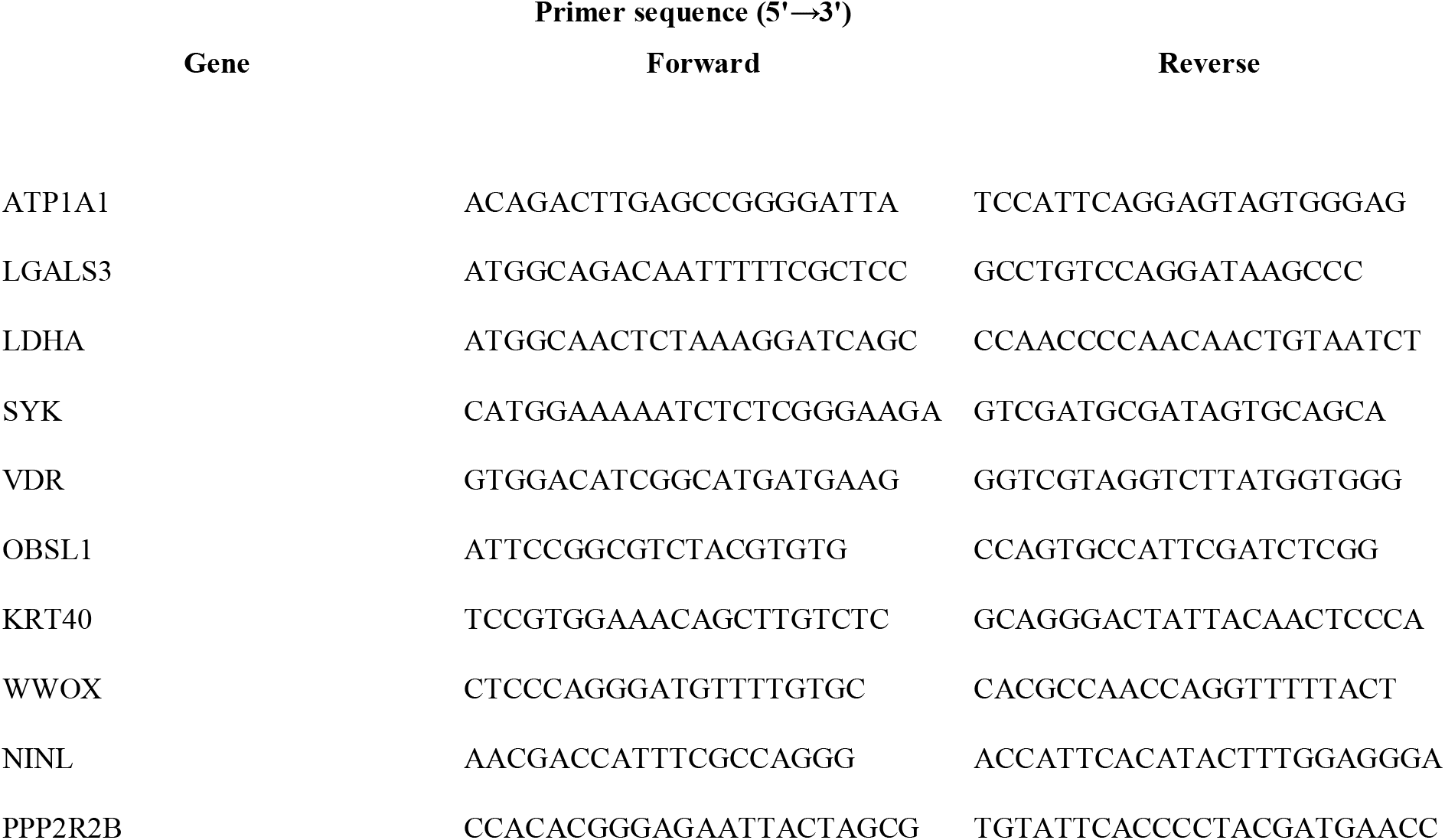
Primers used for quantitative PCR

### Molecular docking studies

The Surflex-Docking docking studies were conducted using the perpetual software module SYBYL-X 2.0 for the built molecules. The molecules were sketched using ChemDraw Tools, imported and saved into sdf. format using free Open Babel software. The co-crystallised protein structures of CFTR, EZR, ATP1A1, TUBA1C and LGALS3 and their co-crystallized PDB code of 4WZ6, 4RM8, 3KDP, 3E22 and 6FOF respectively were selected for docking and were extracted from Protein Data Bank [52–55]. Optimization of the designed molecules were performed by converting 3D concord structure, applying TRIPOS force field and adding Gasteiger Huckel (GH) charges, In addition, MMFF94s and MMFF94 algorithm processes have been used or energy minimization. After the introduction of protein, protein processing was accomplished. The co-crystallized ligand and all water molecules were expelled from the crystal structure; more hydrogen was added and the side chain was refined. The TRIPOS force field was used to reduce structure complexity and the interaction efficiency of the compounds with the receptor was represented in kcal/mol units by the Surflex-Dock score. The best spot between the protein and the ligand was inserted into the molecular region. The visualisation of ligand interaction with receptor is done by using discovery studio visualizer.

## Results

### DEG screening

In the current study, DEGs and their significant biological characteristics were diagnosed based on GEO data set by integrated bioinformatics analysis of NET. The NET expression microarray data set GSE65286 was first selected and down loaded. There were a total of 43 samples including 33 patients with metastatic NET and 10 normal controls. After gene expression data processing and normalizing (Fig 1A and Fig 1B), we screened DEGs among GEO data set using the limma package with corrected p < 0.05, and |logFC|> 3.01 for up regulated genes and |logFC|> −2.75 for down regulated genes (Table 2). There were 912 DEGs including 459 up regulated and 453 down regulated genes. The volcano plots of DEGs is shown in Fig. 2. A heat map demonstrated the significant differential distribution of the DEGs (up and down regulated genes) using data profile GSE65286 as a reference (Fig. 3A and Fig. 3B).

**Figure 1.**
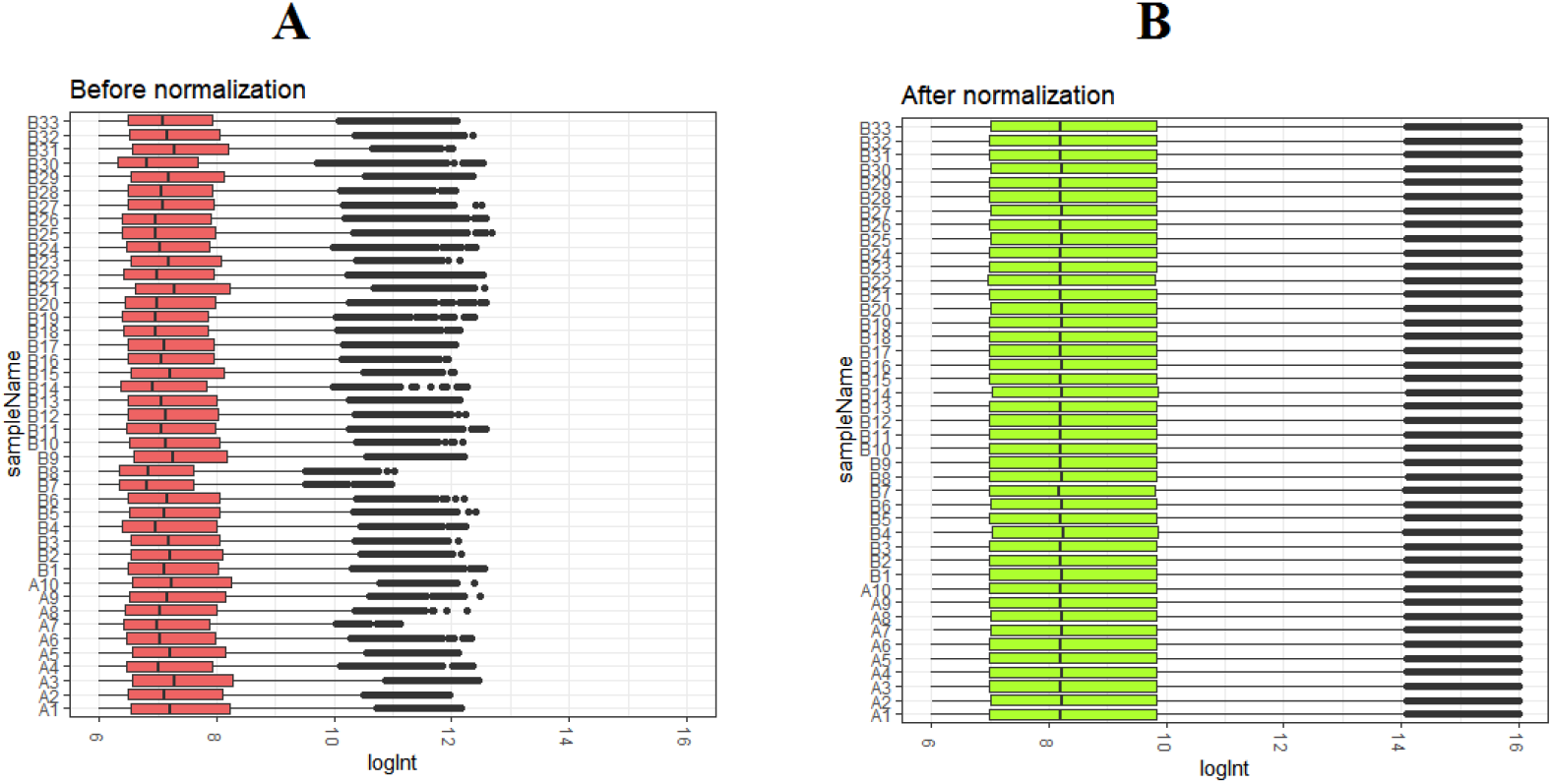
Box plots of the gene expression data before and after normalization. Horizontal axis represents the sample symbol and the vertical axis represents the gene expression values. The black line in the box plot represents the median value of gene expression. (A1-A10= normal control; B1-, B33 = metastatic NETs)

**Figure 2.**
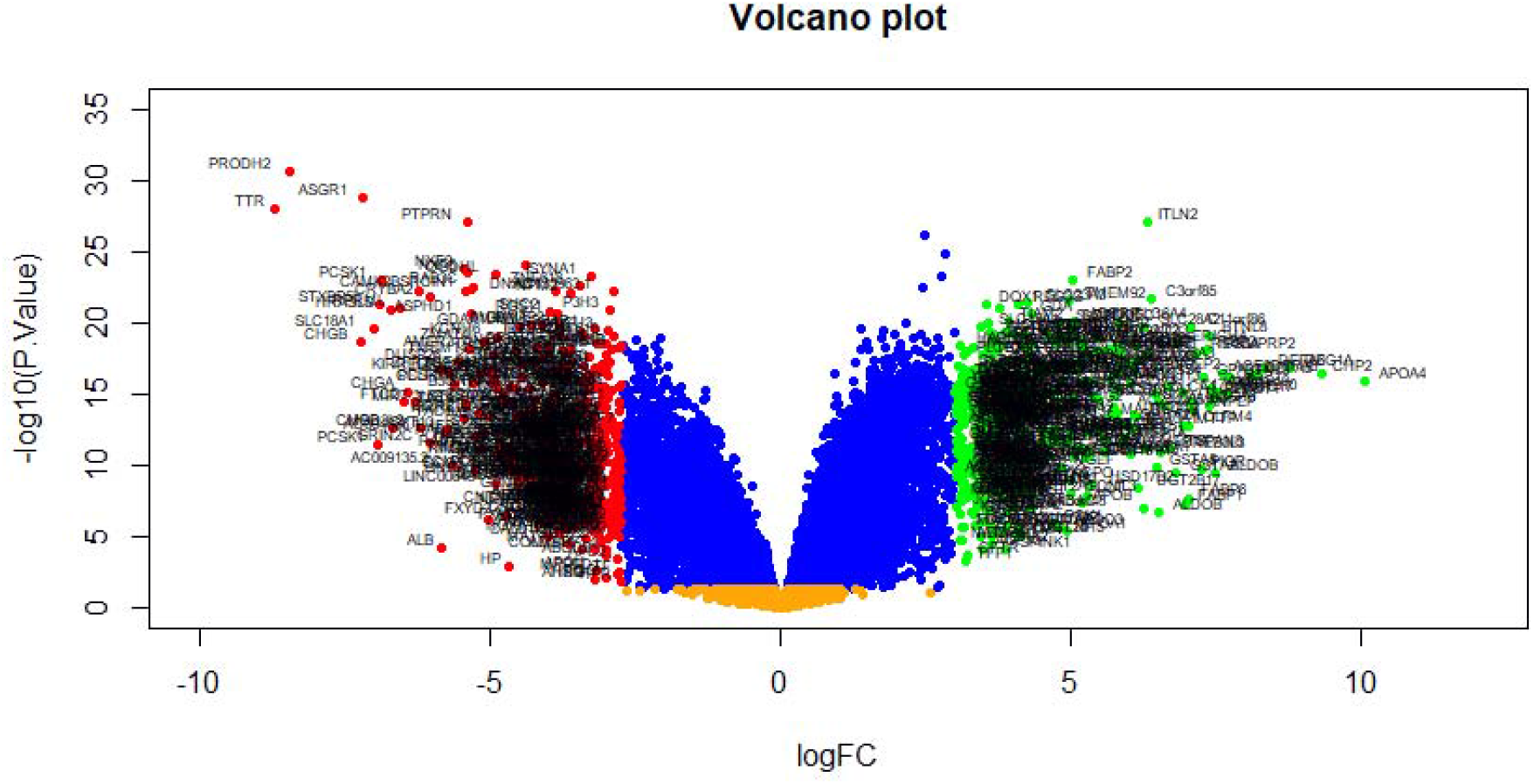
Volcano plot of differentially expressed genes. Genes with a significant change of more than two-fold were selected.

**Figure 3.**
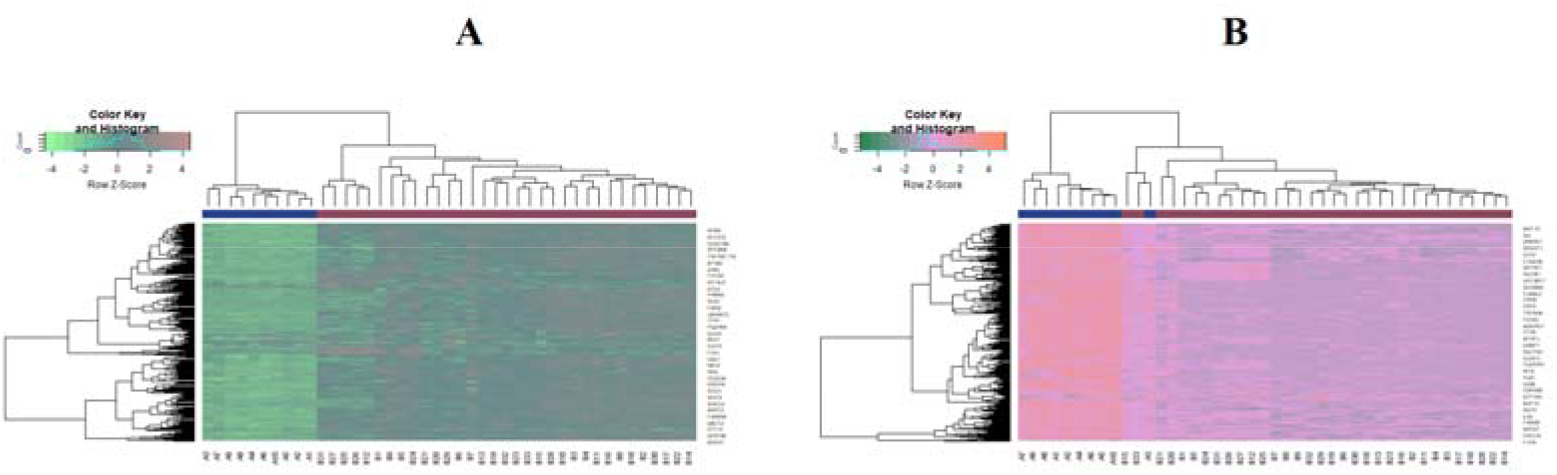
Heat map of (A) up regulated differentially expressed genes (B) down regulated differentially expressed genes. Legend on the top left indicate log fold change of genes. (A1-A10= normal control; B1-, B33 = metastatic NETs)

**Table 2.**
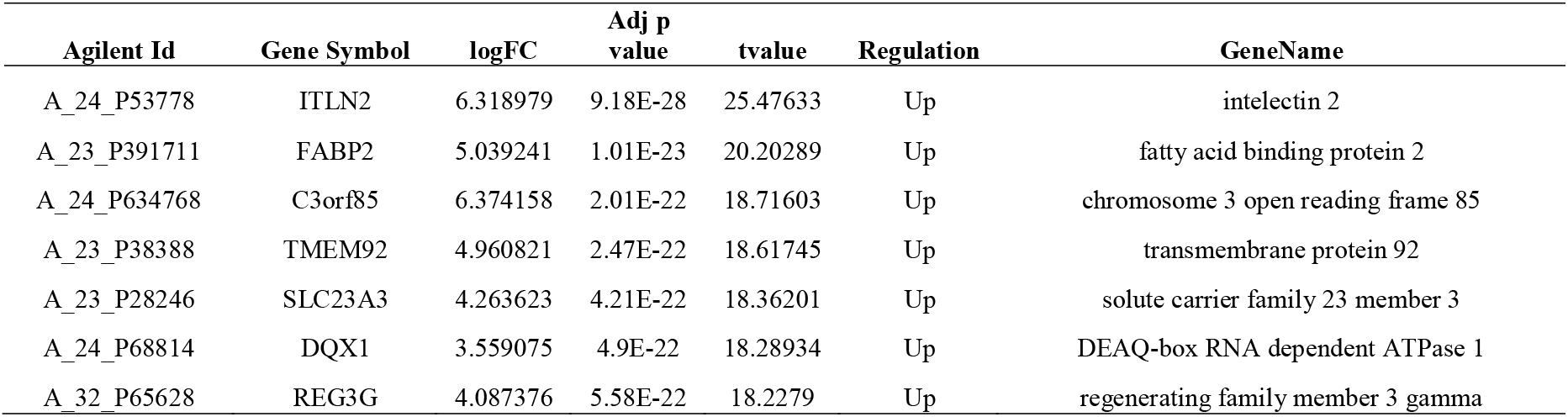
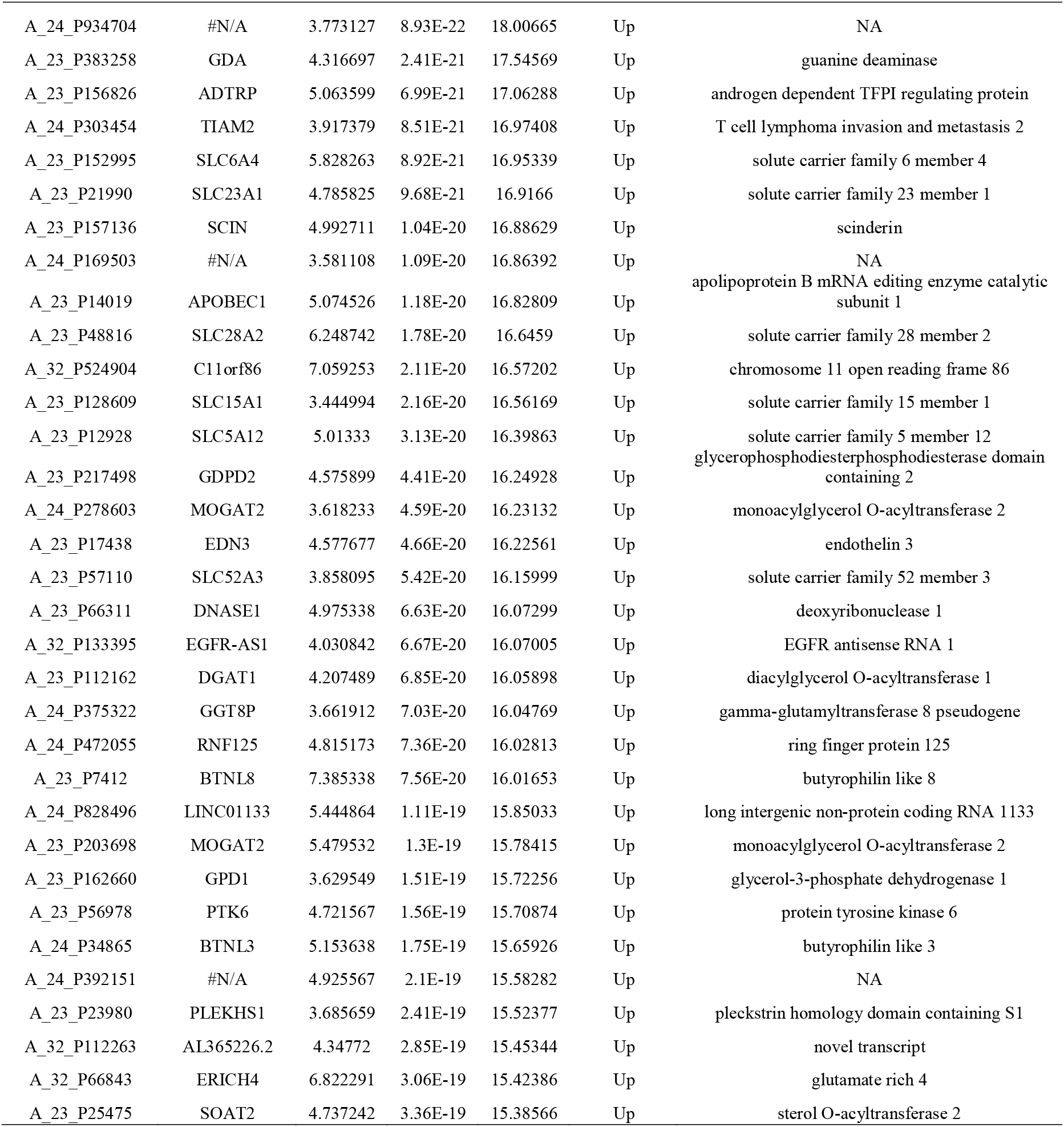
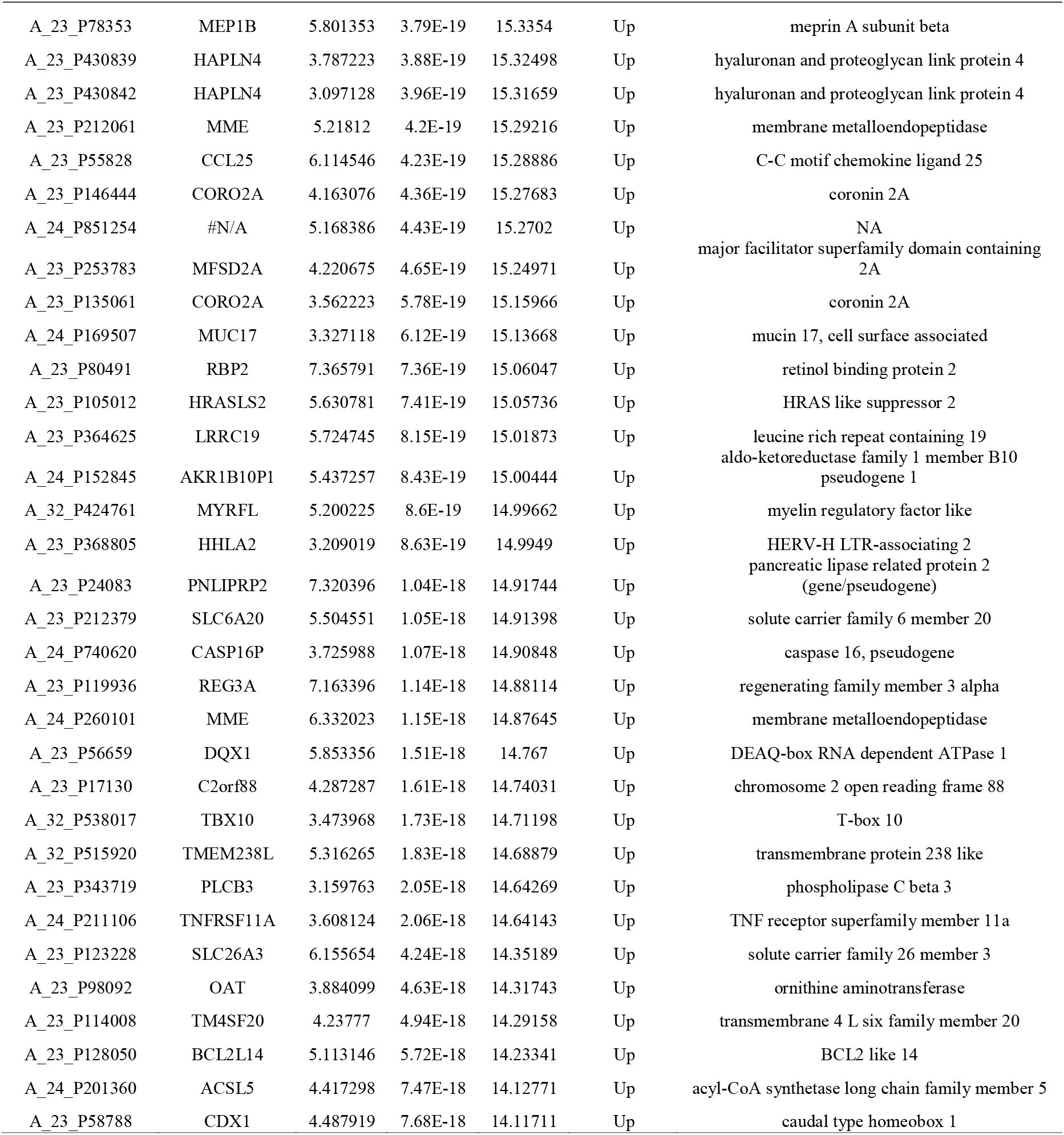
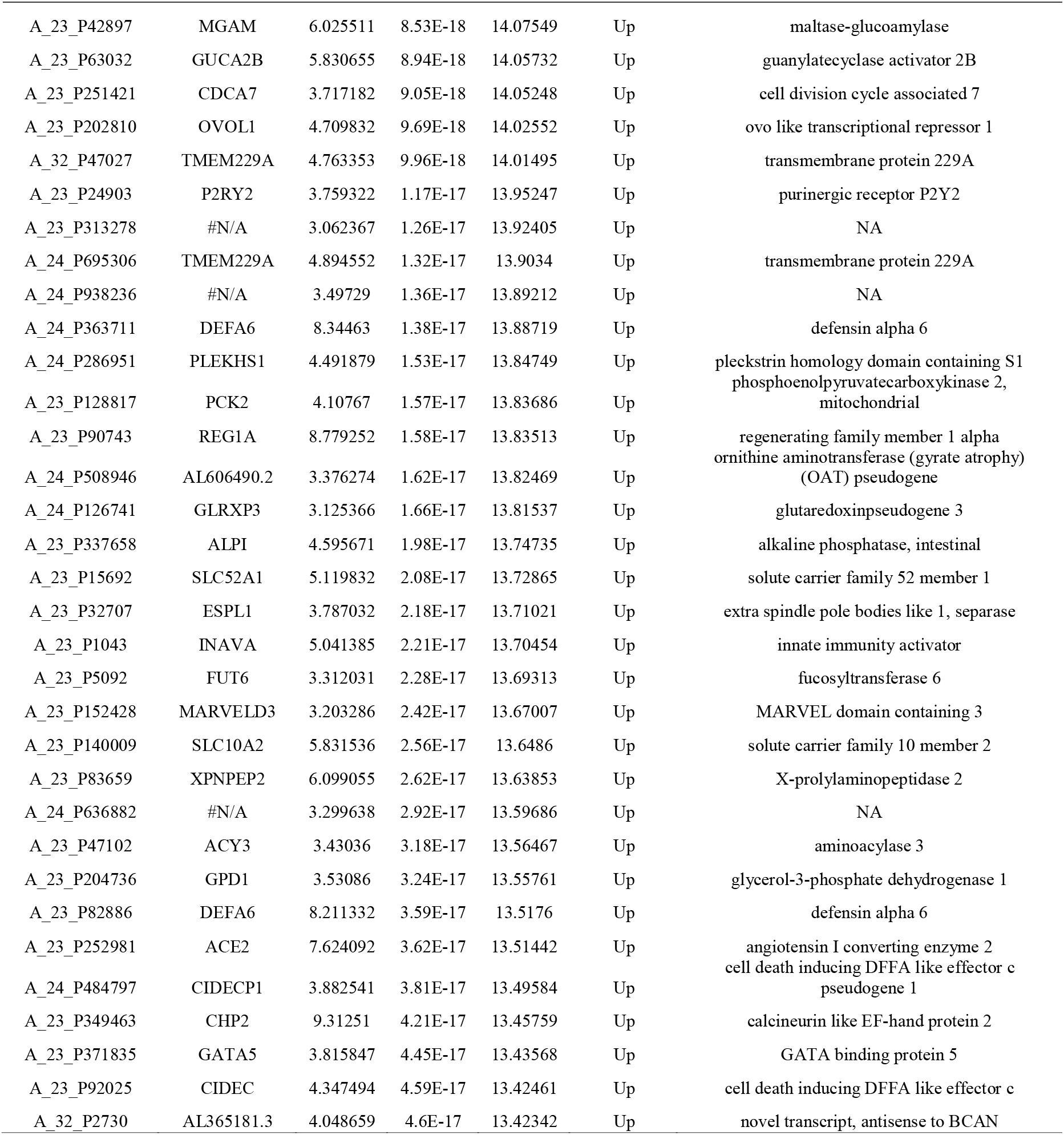
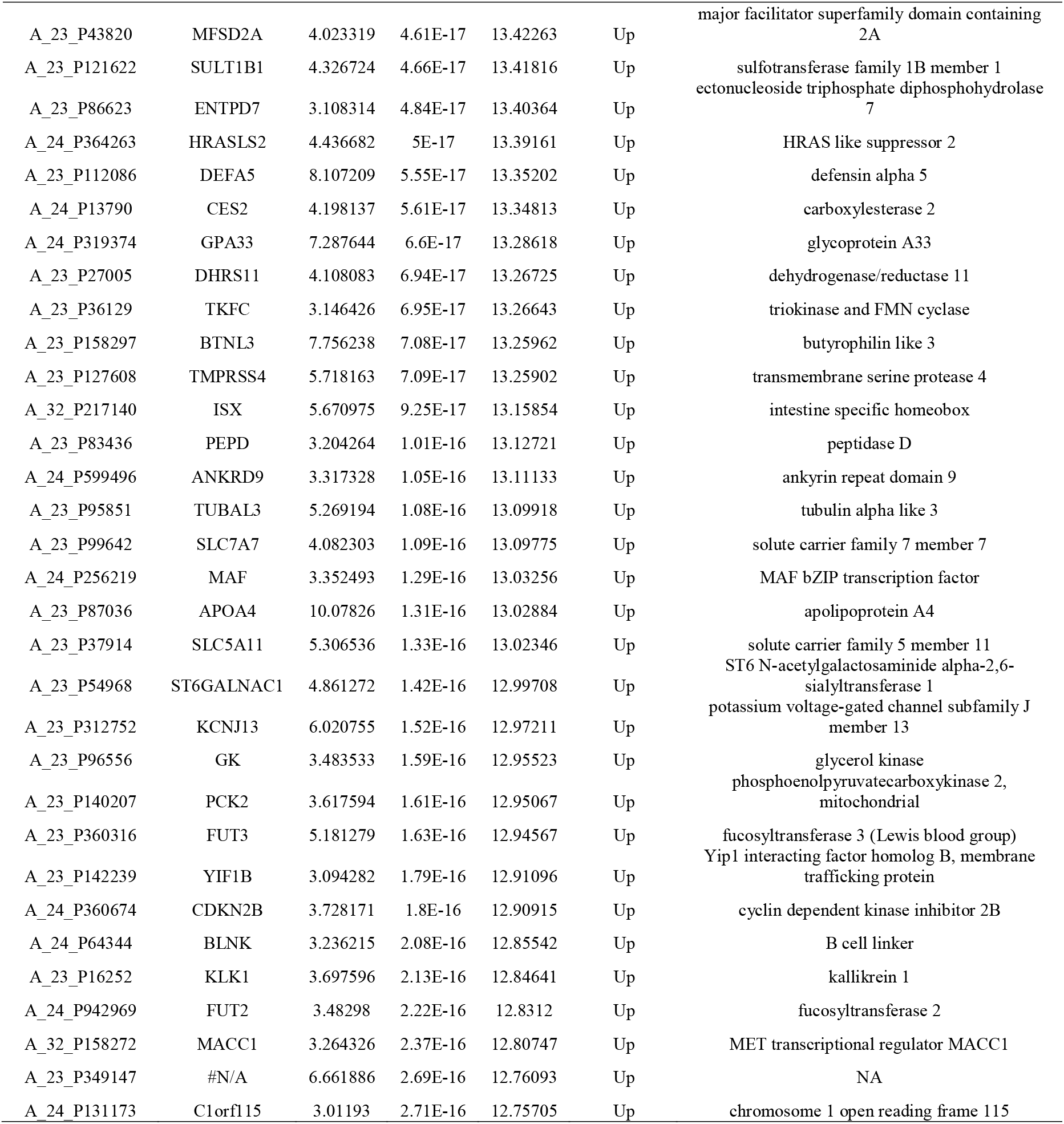
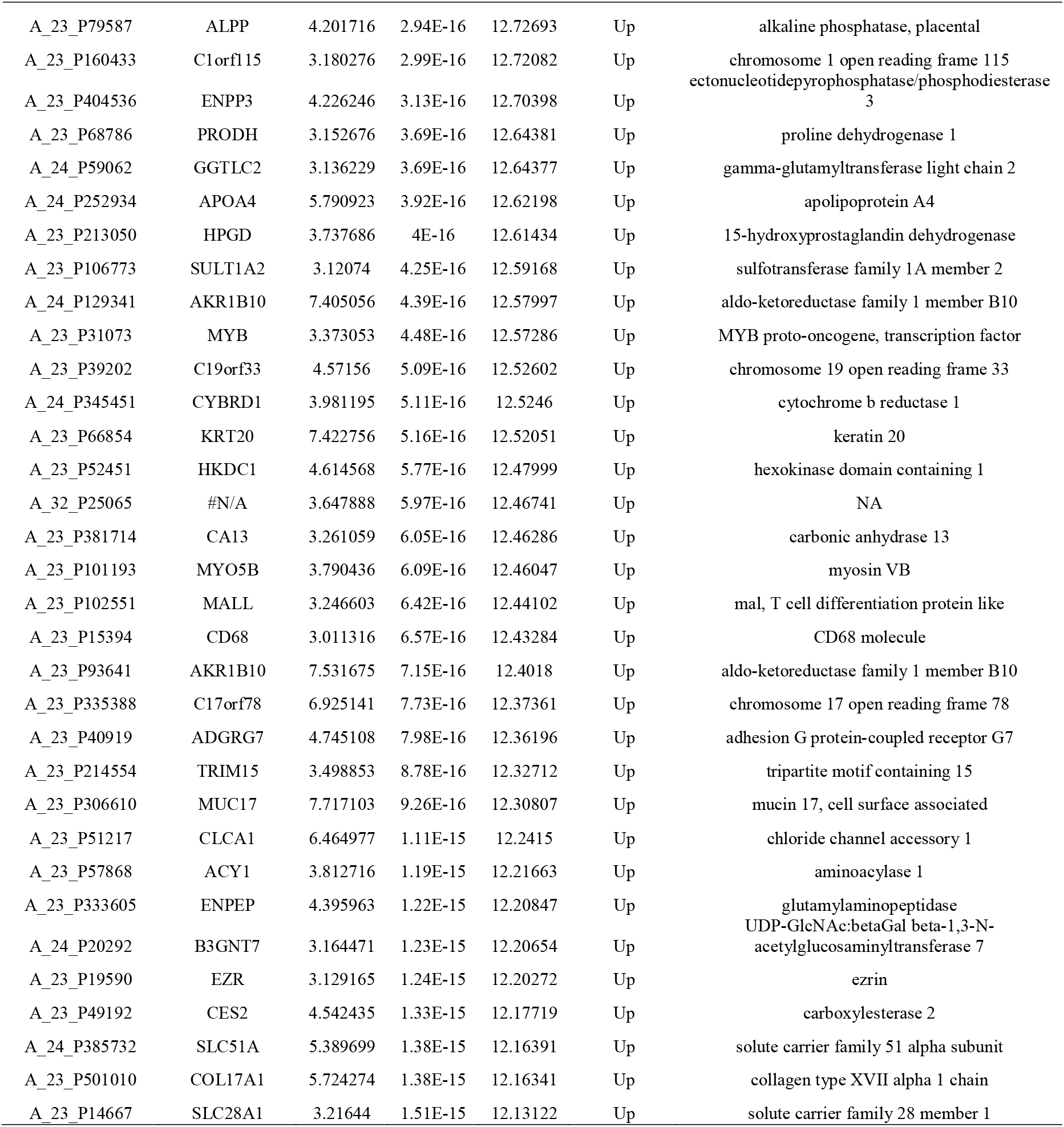
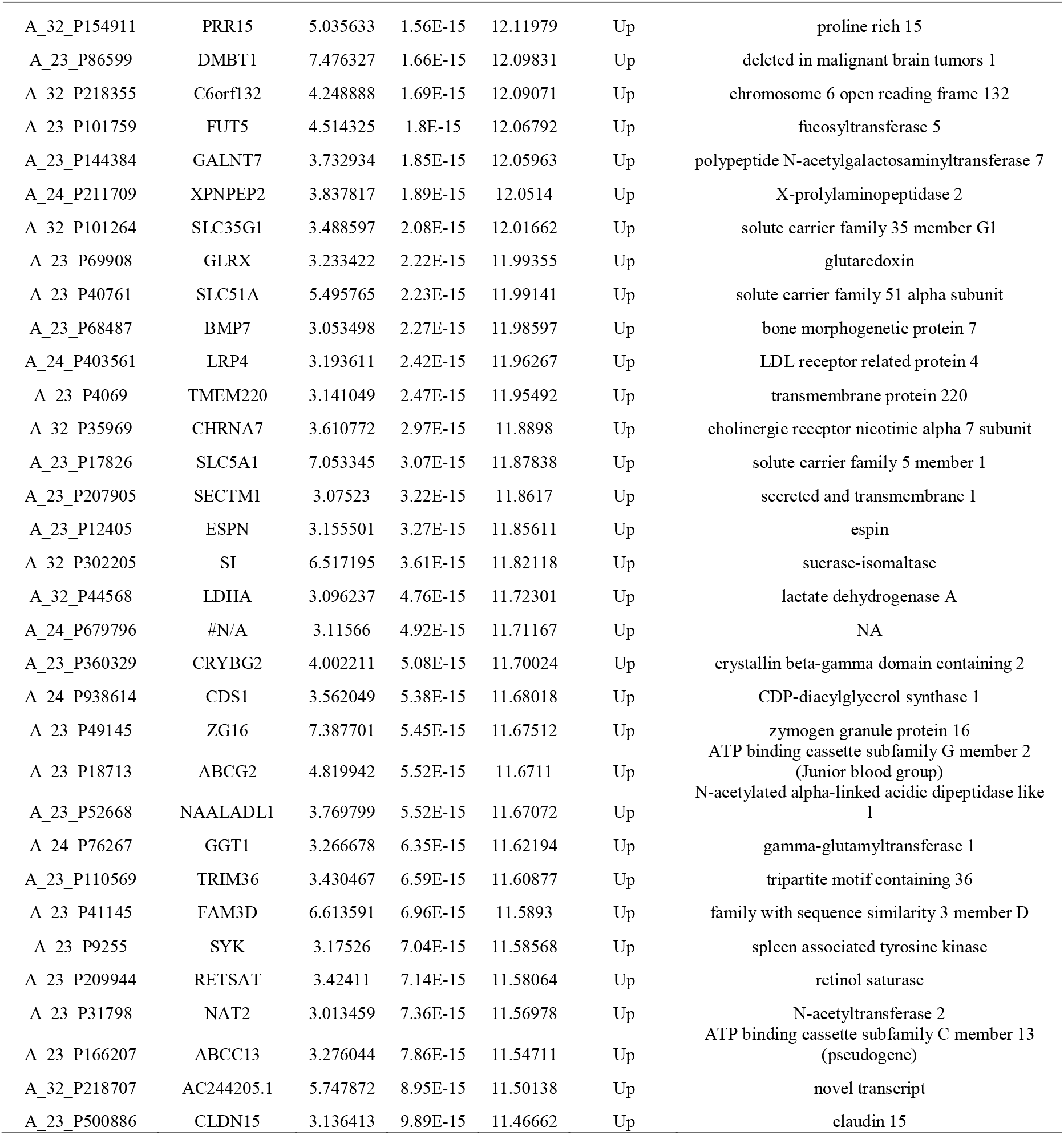
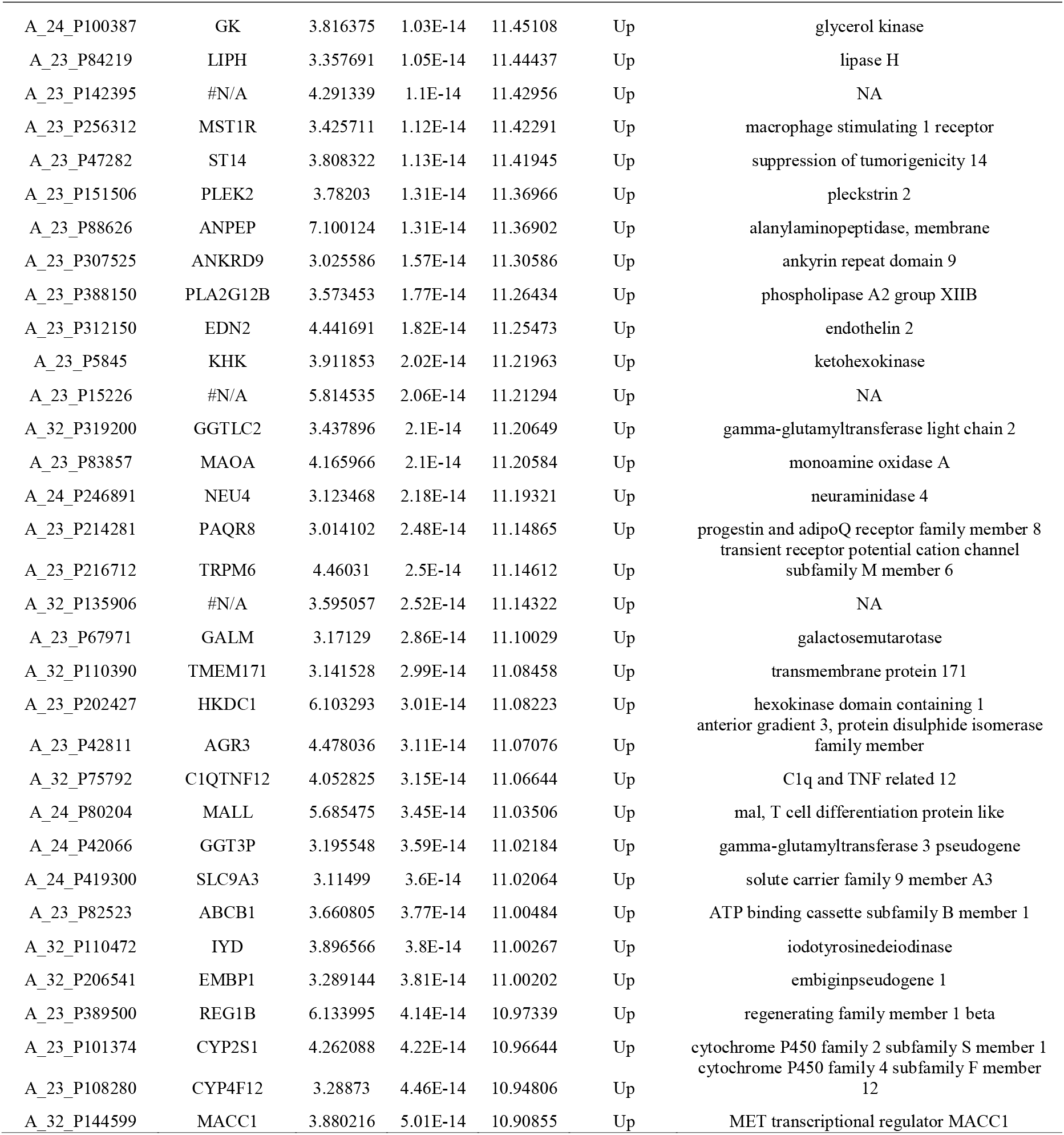
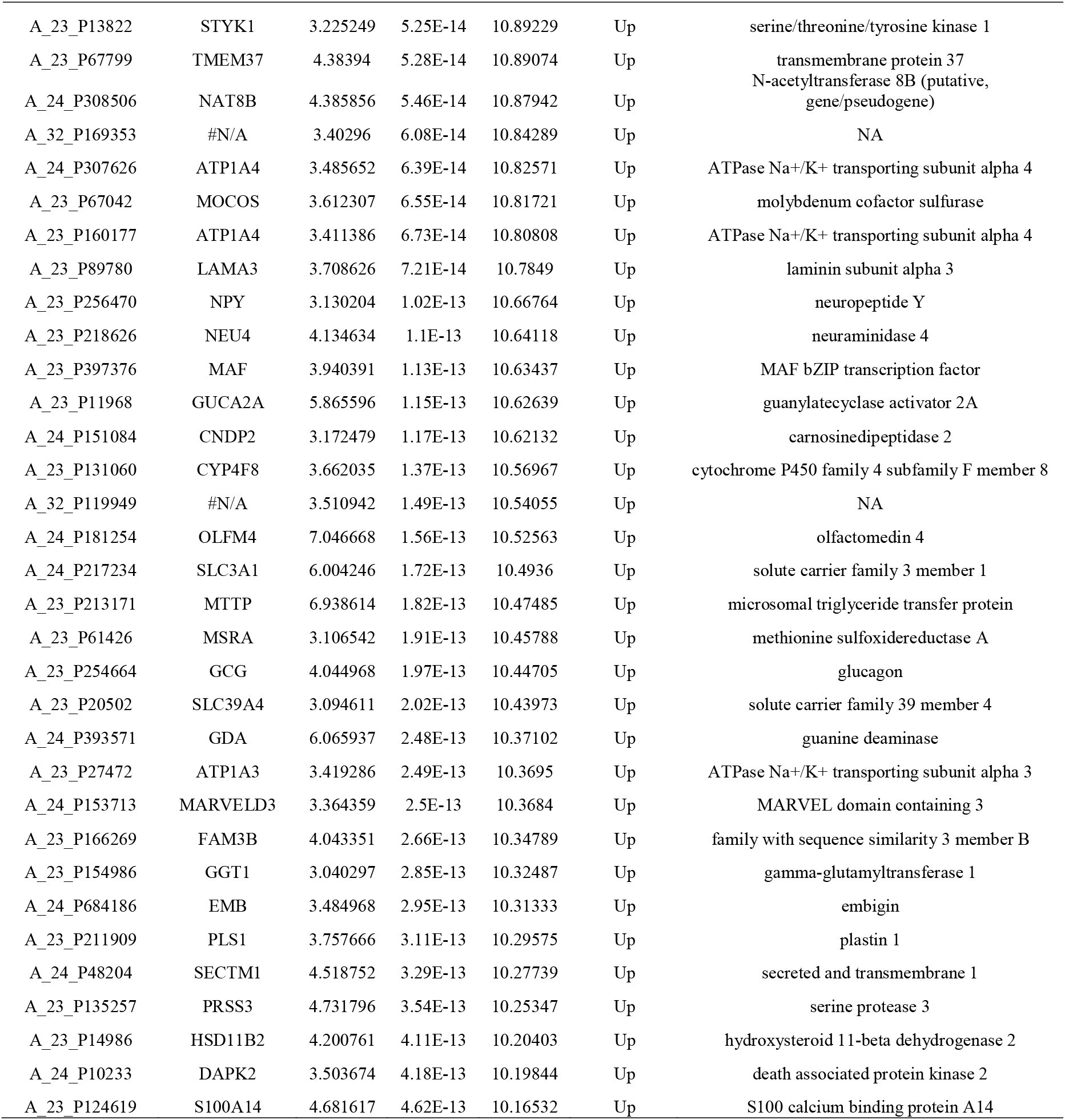
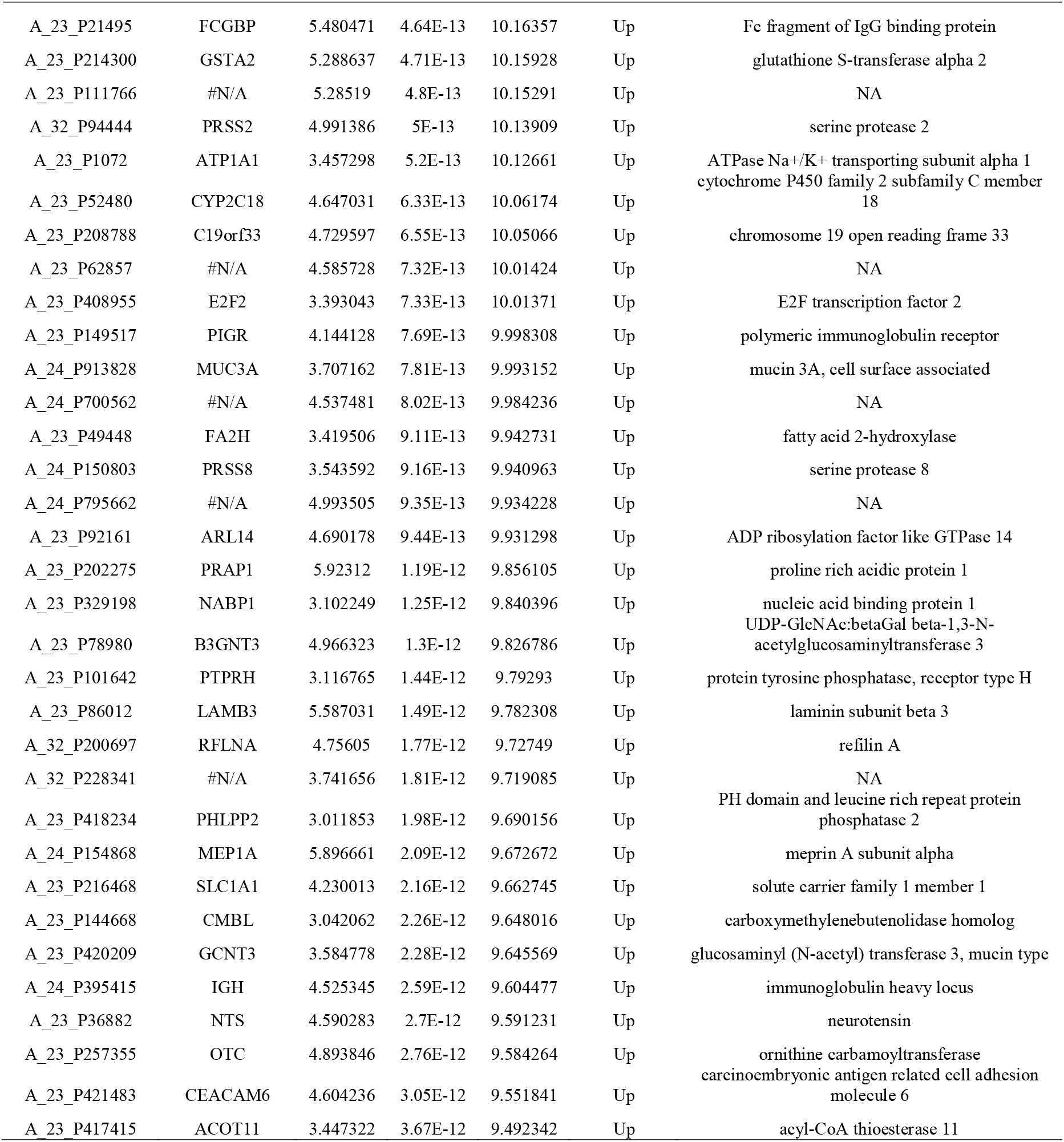
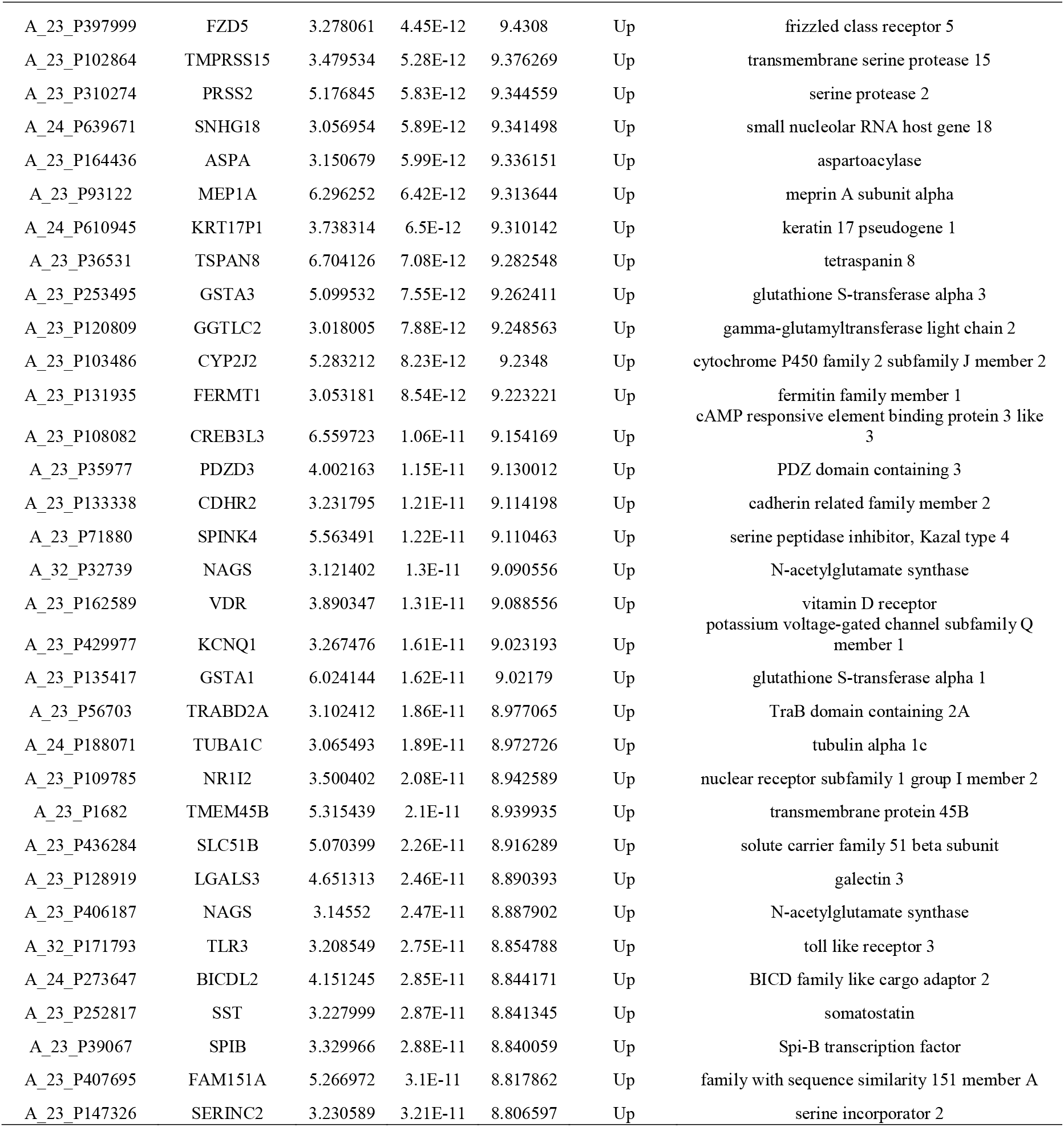
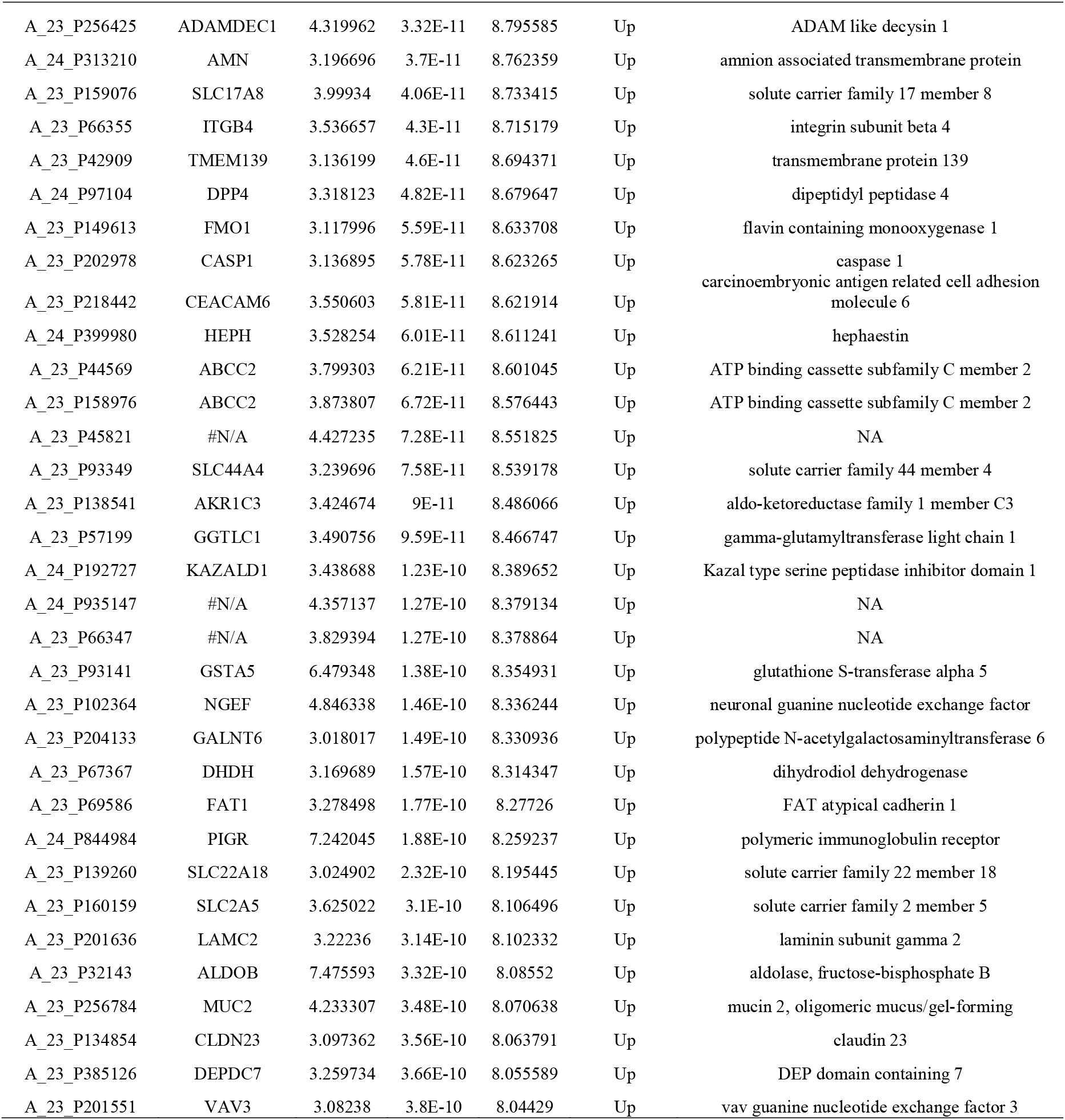
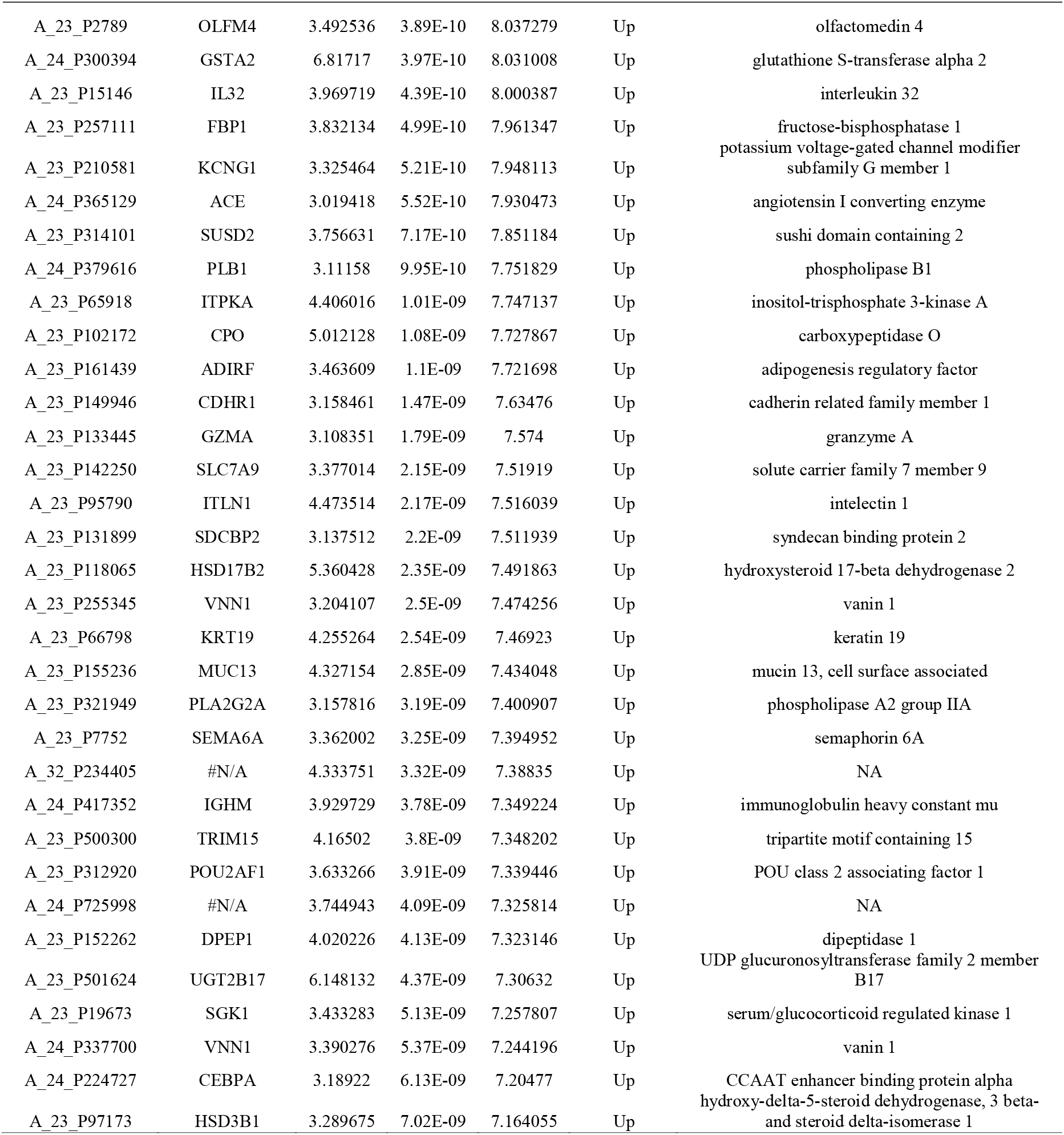
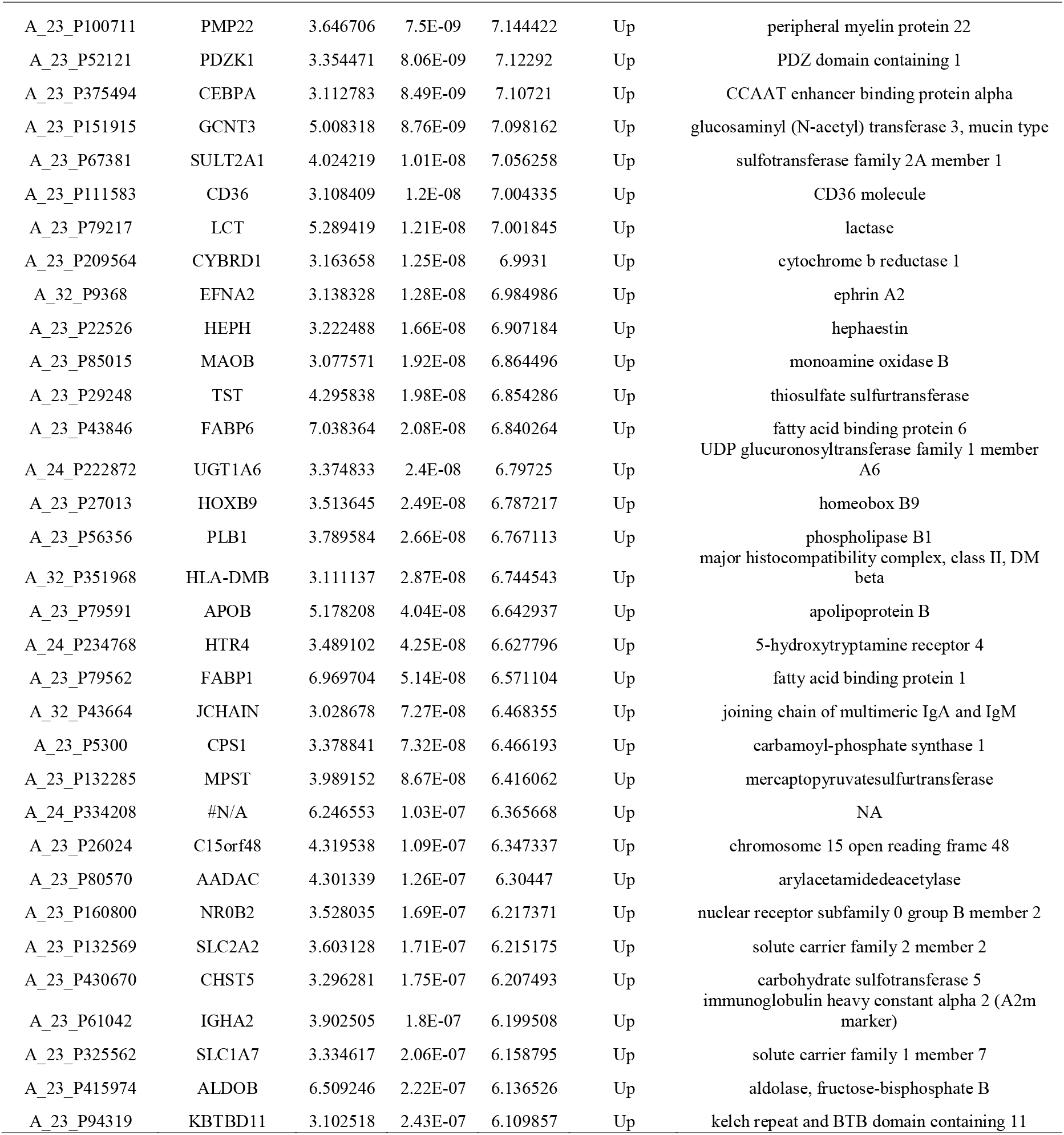
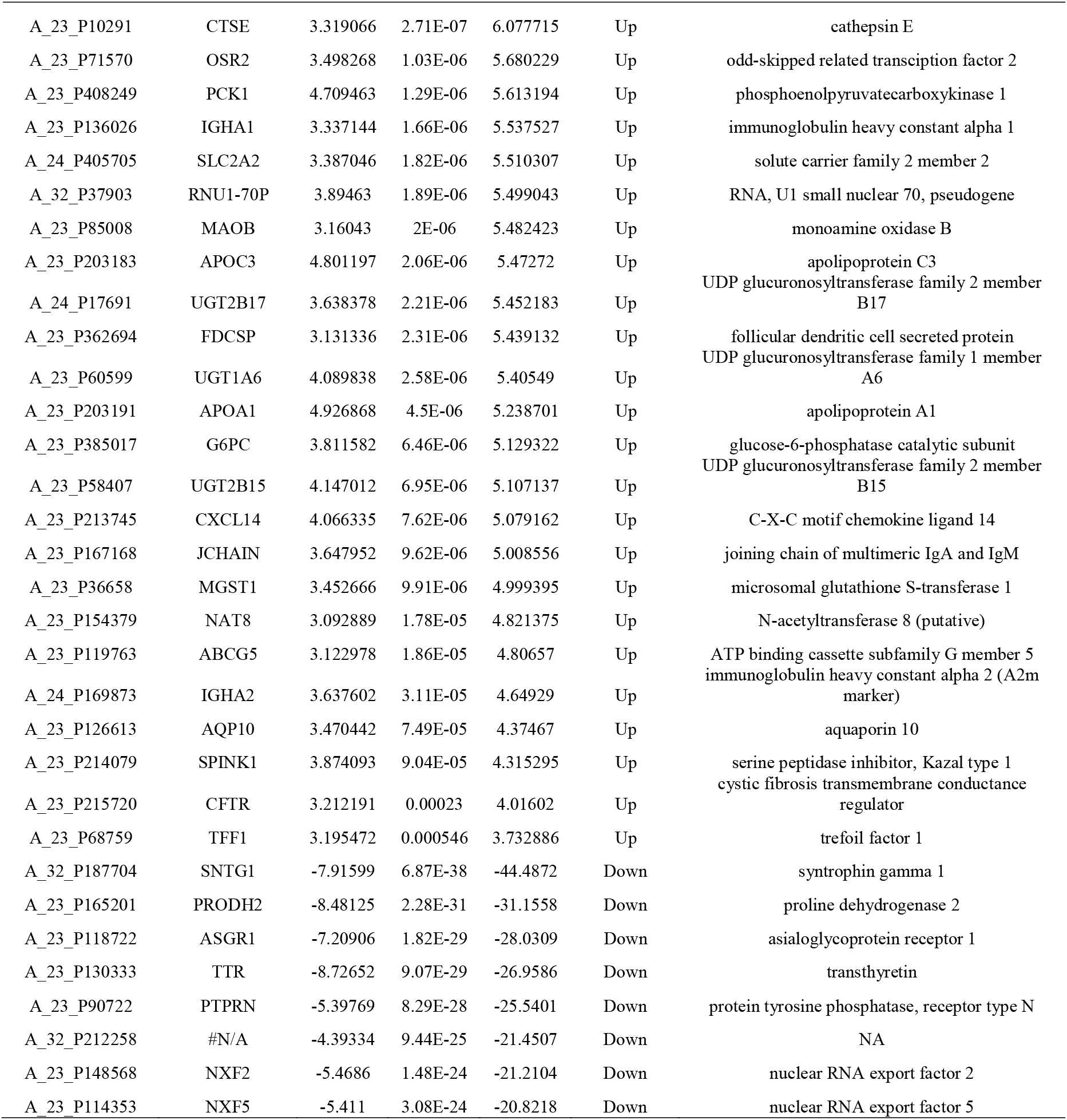
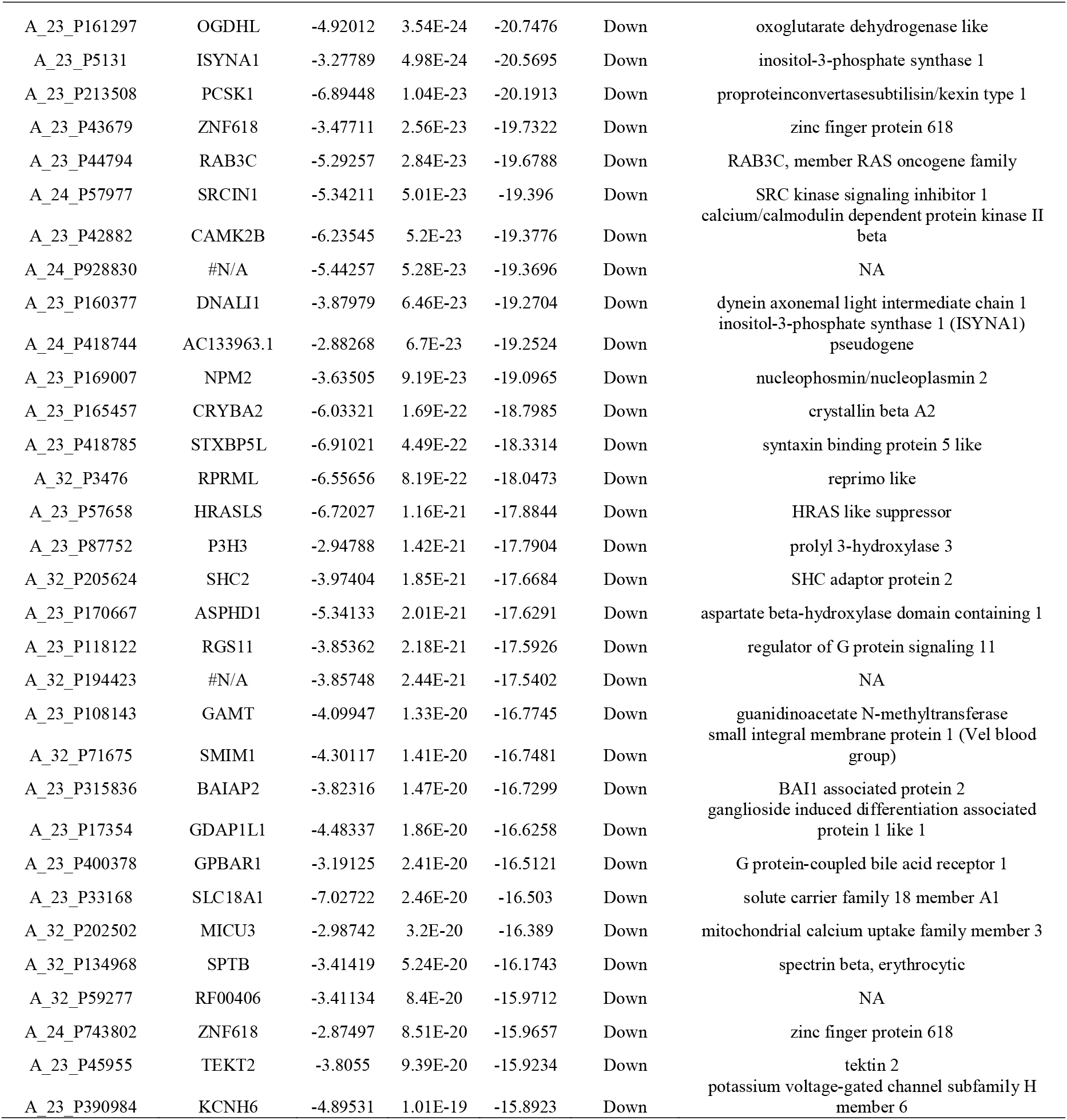
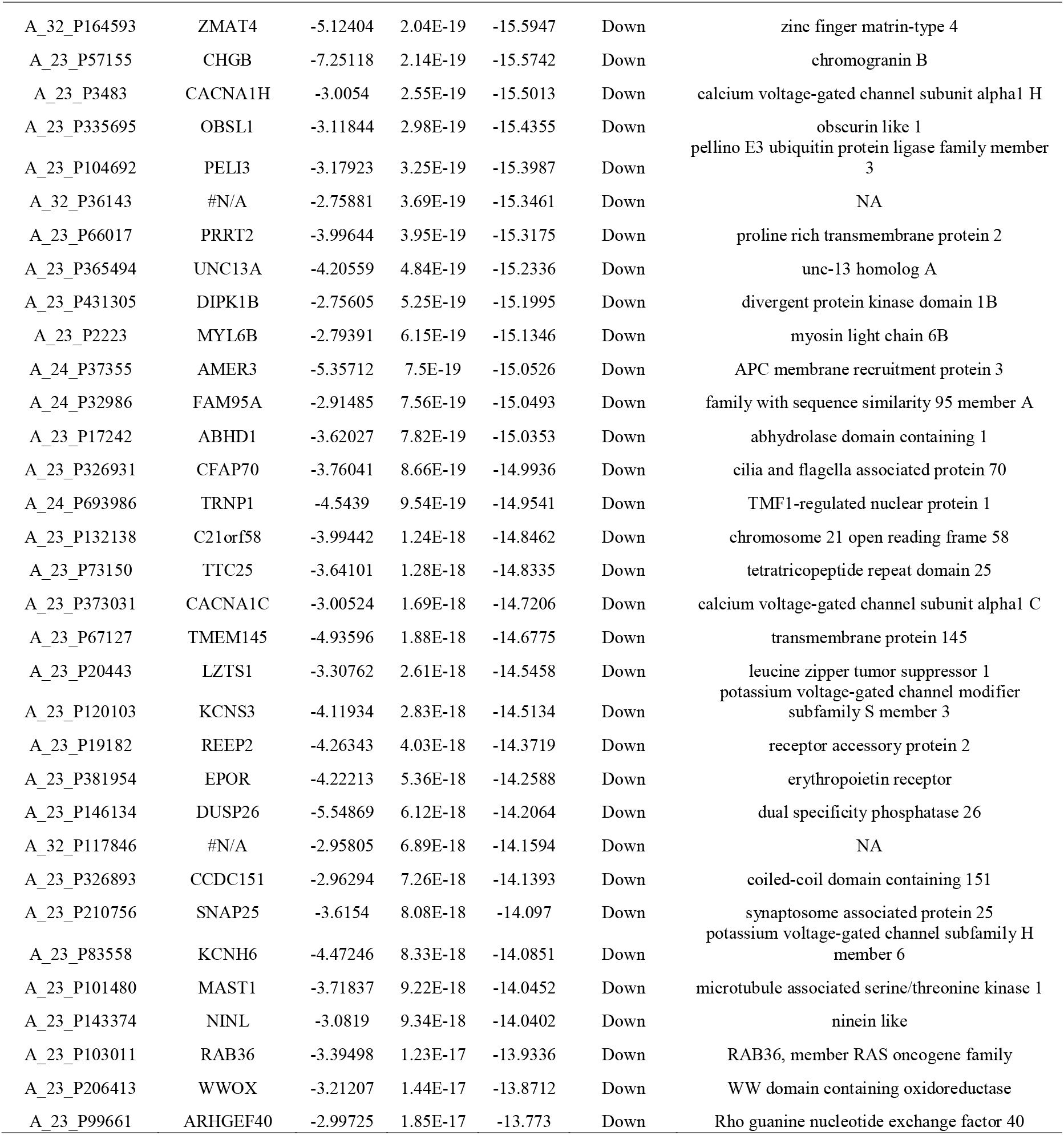
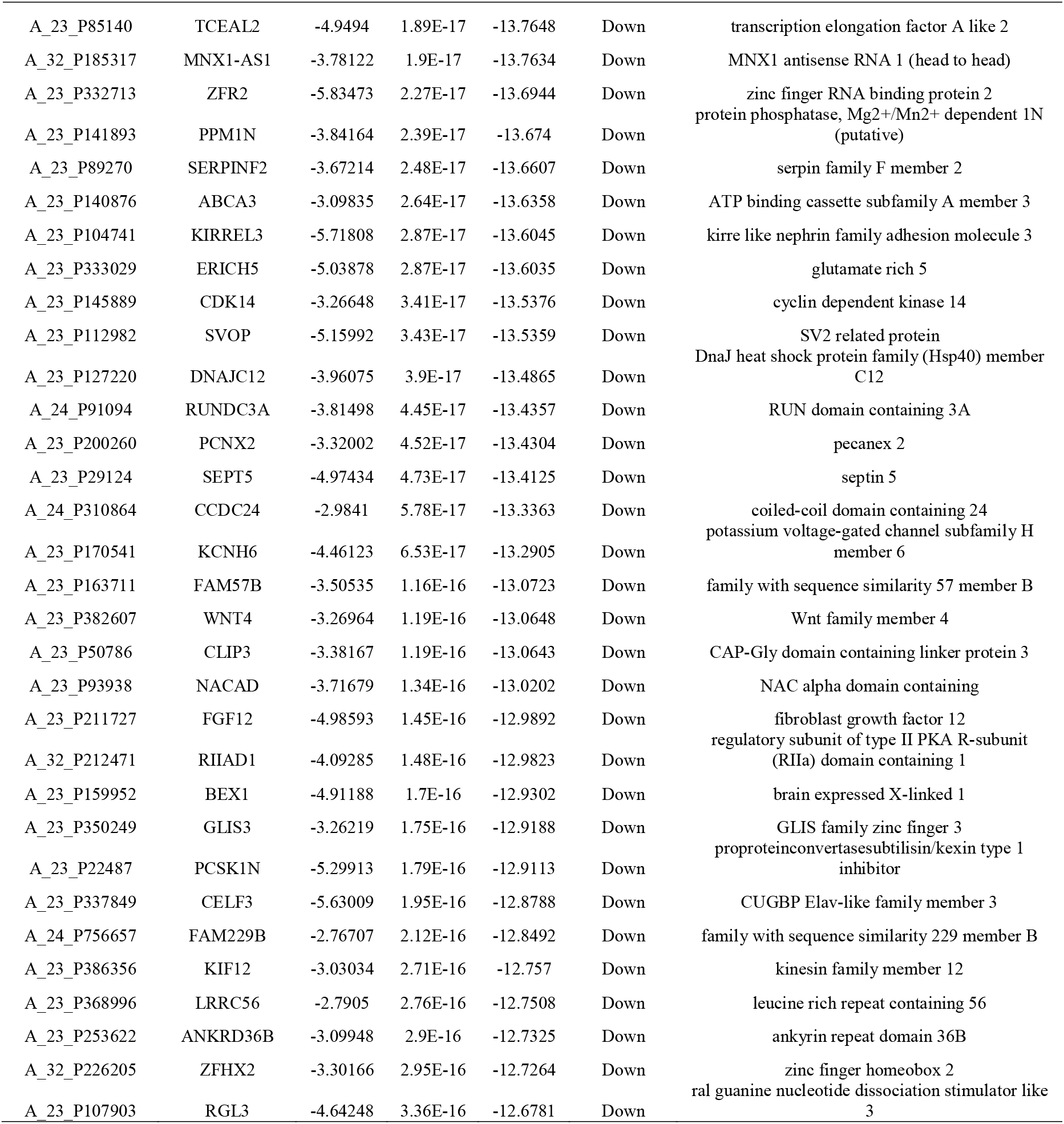
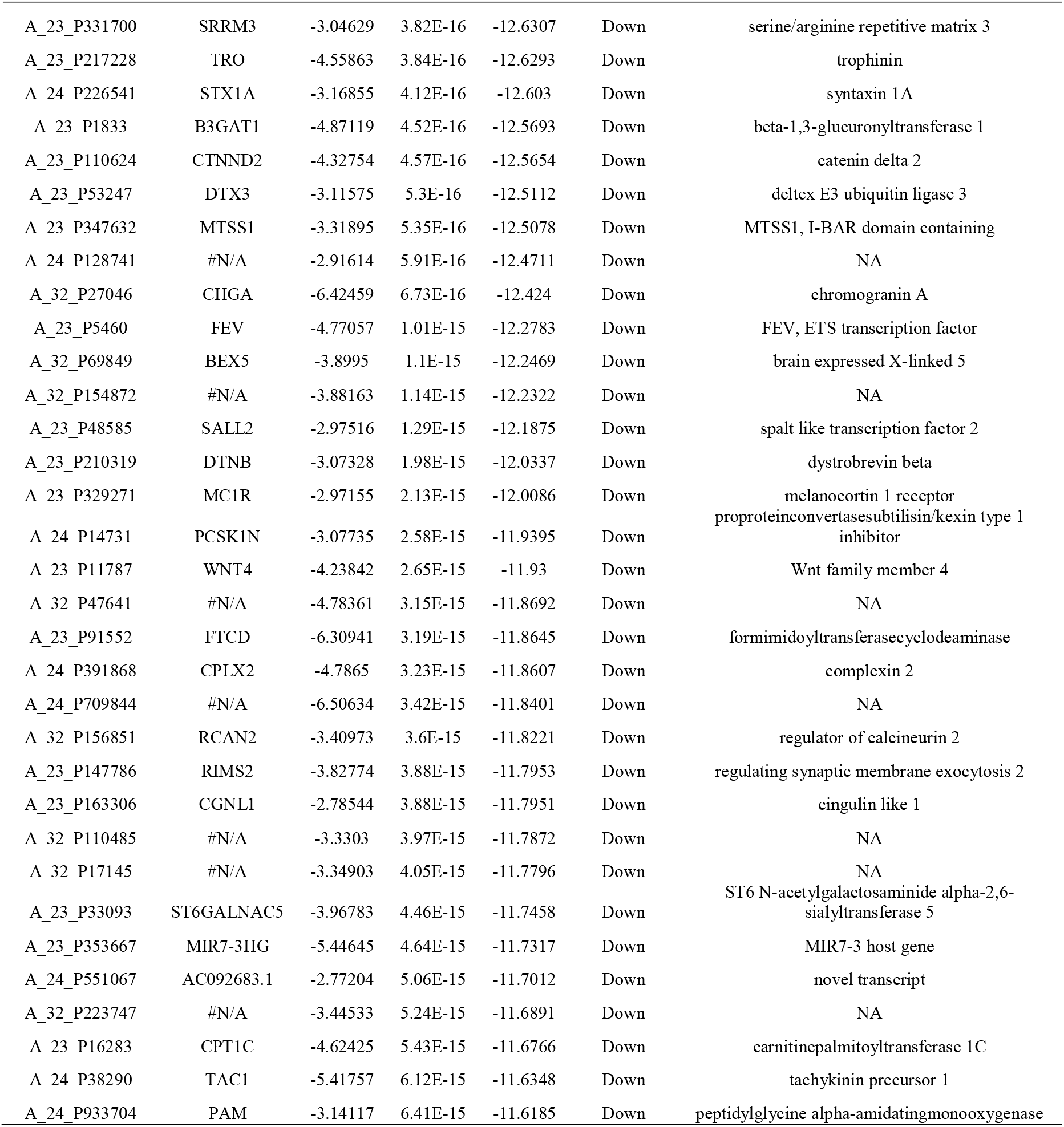
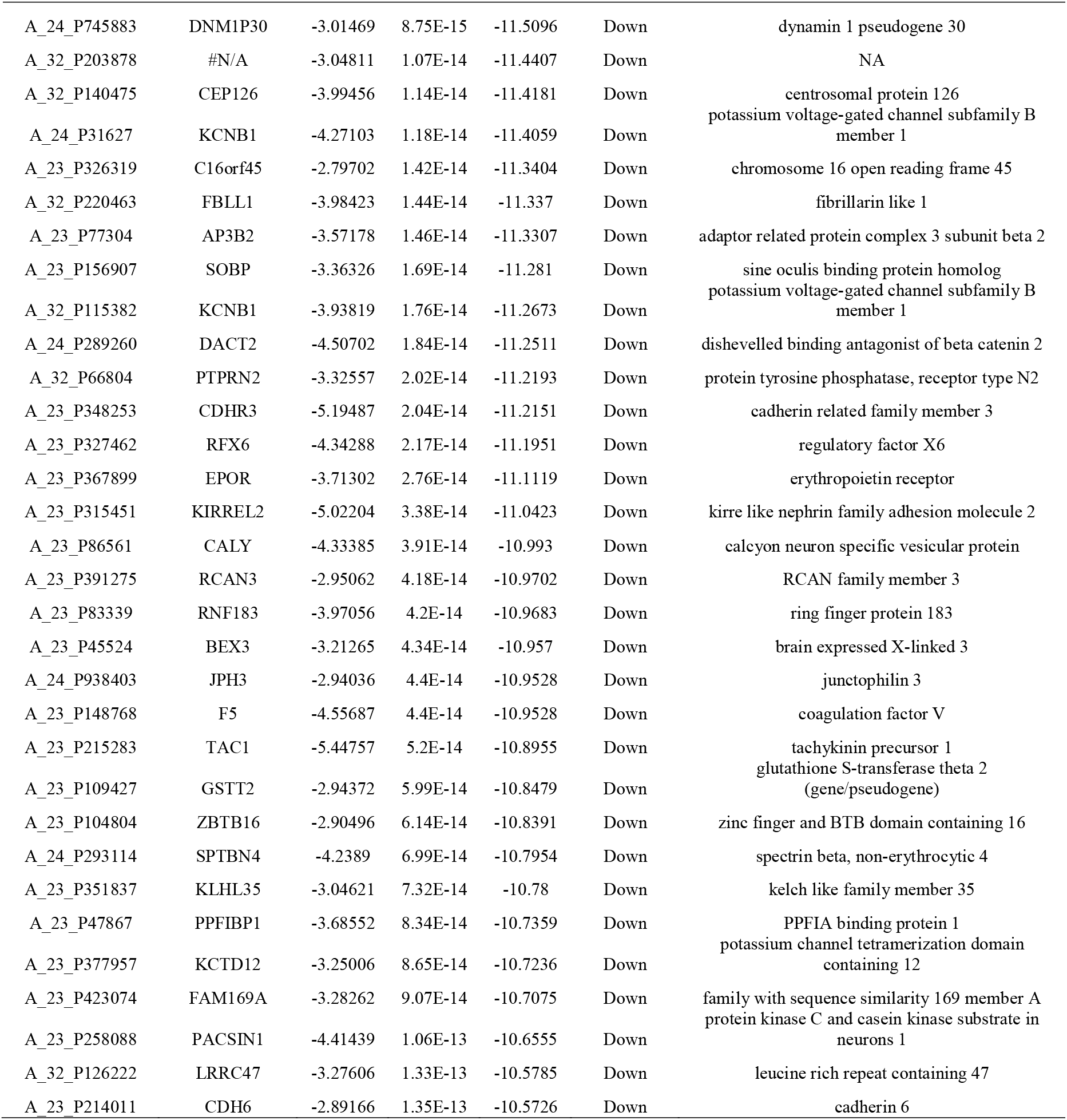
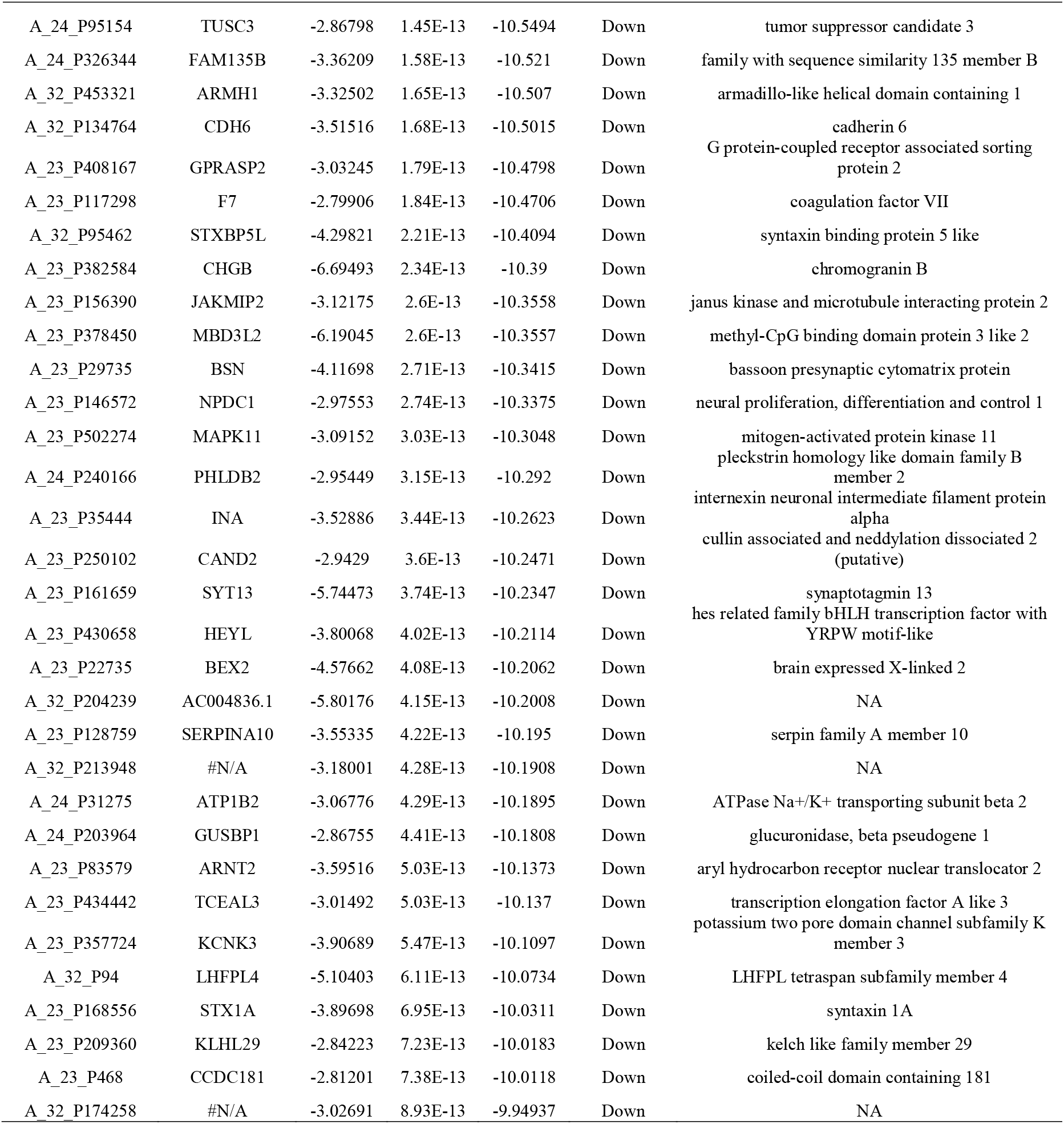
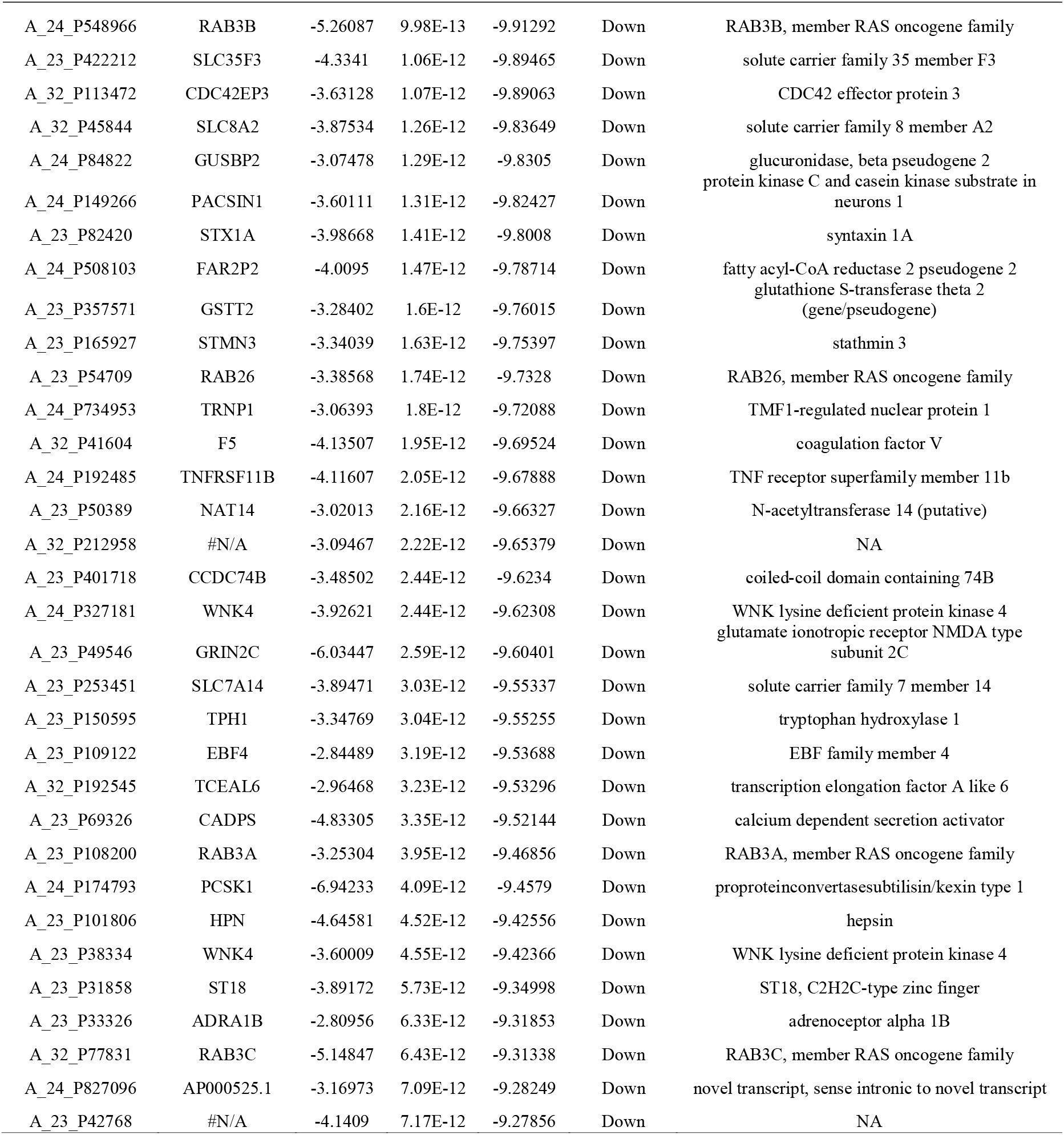
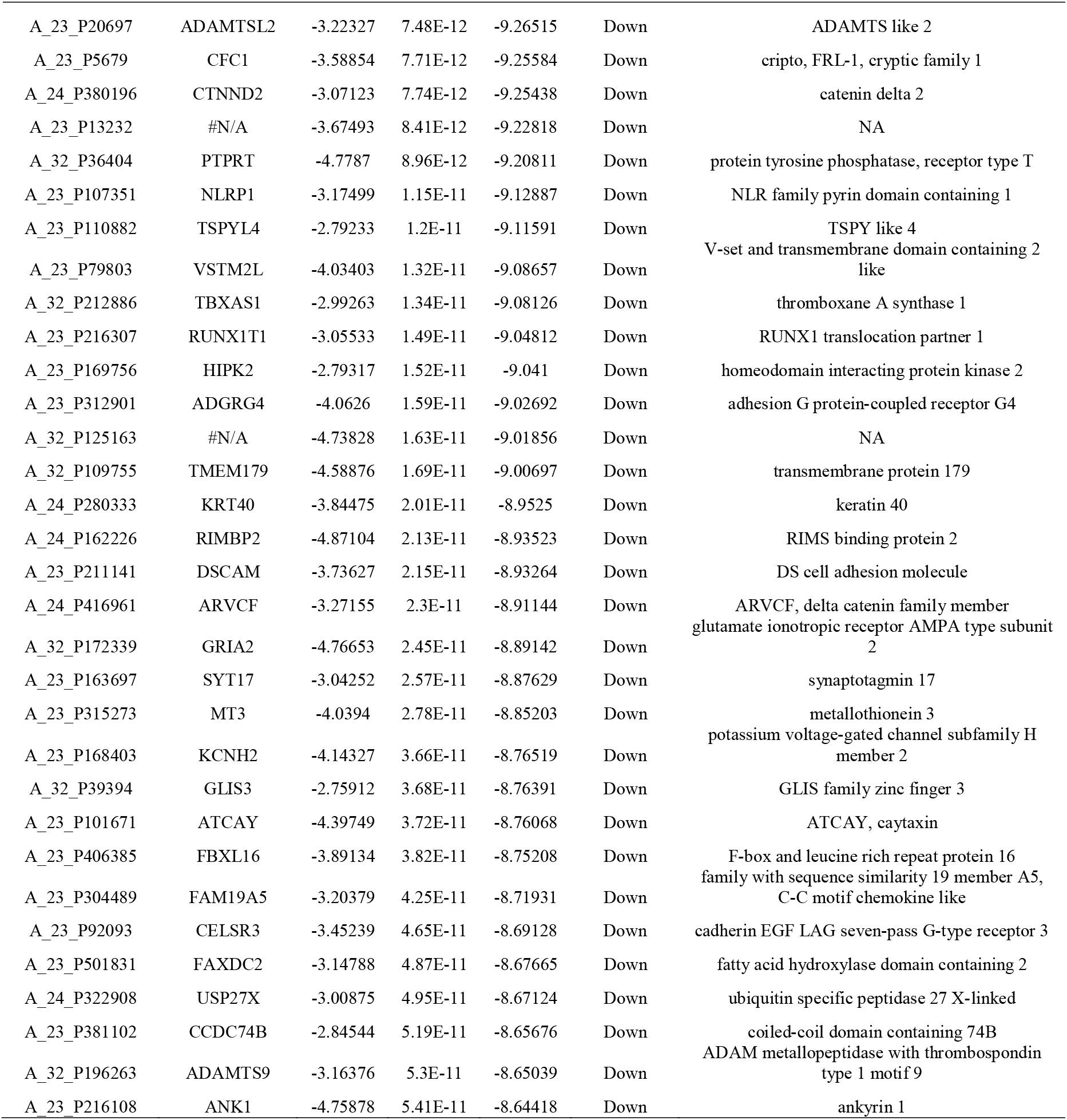
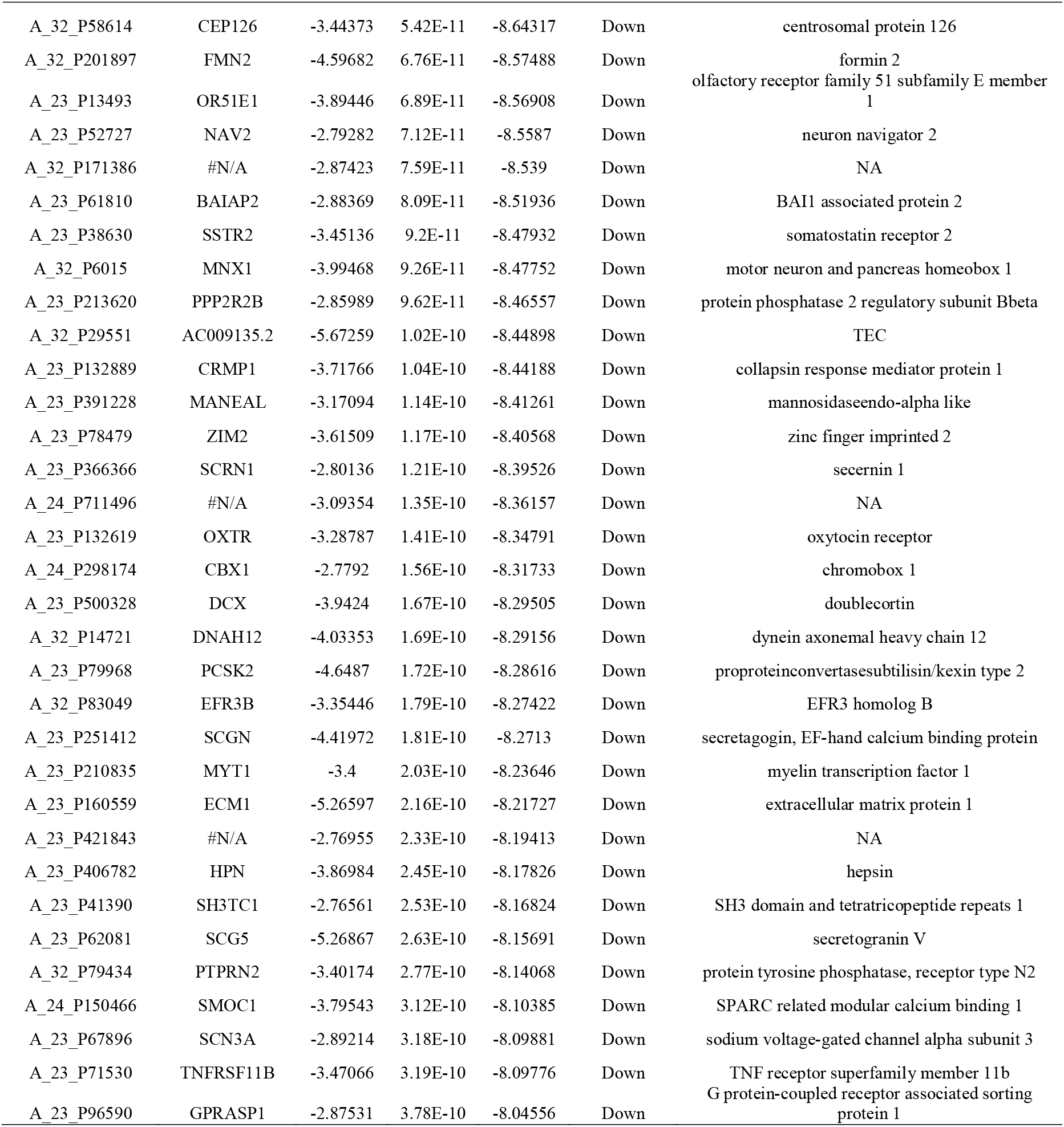
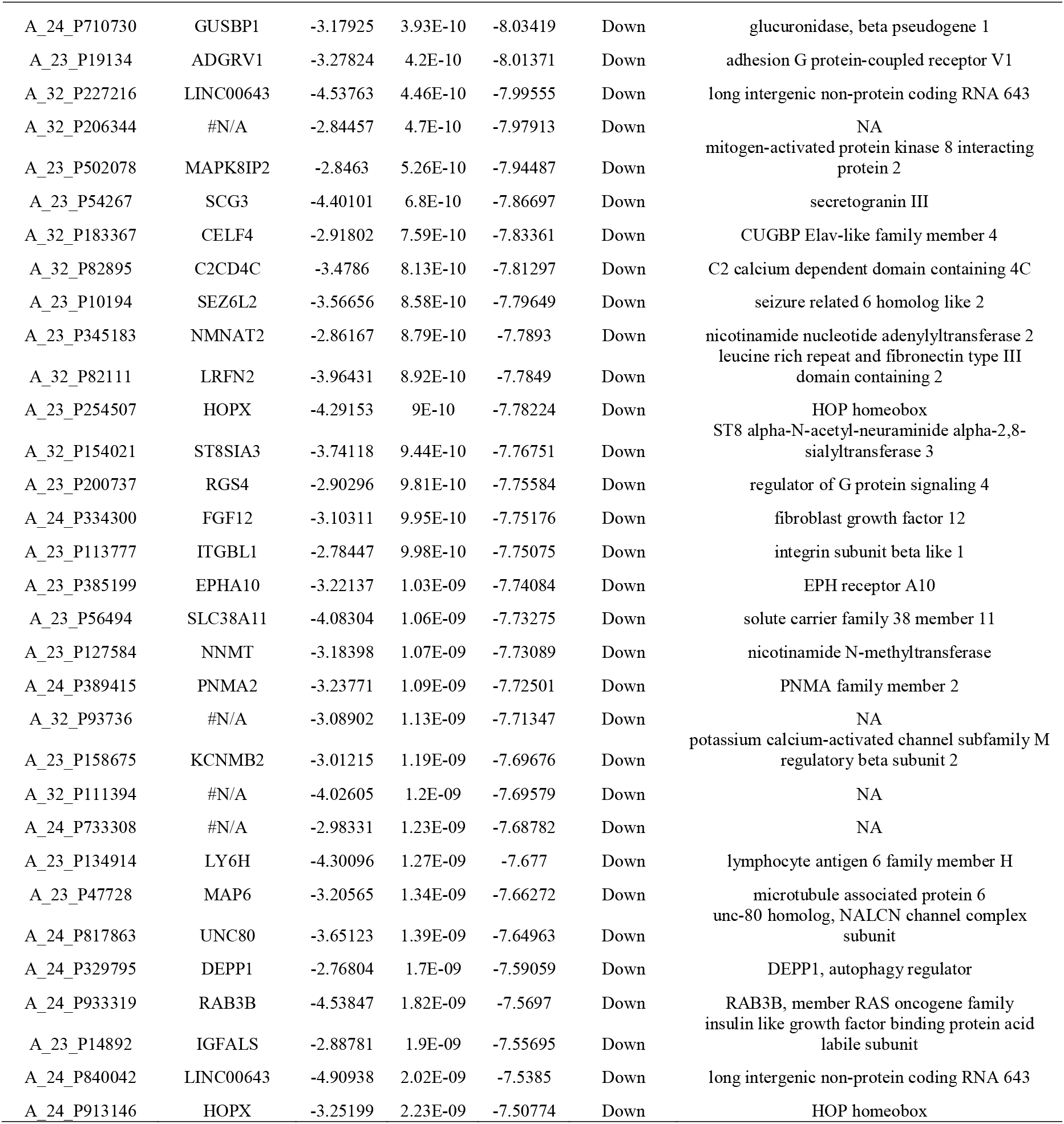
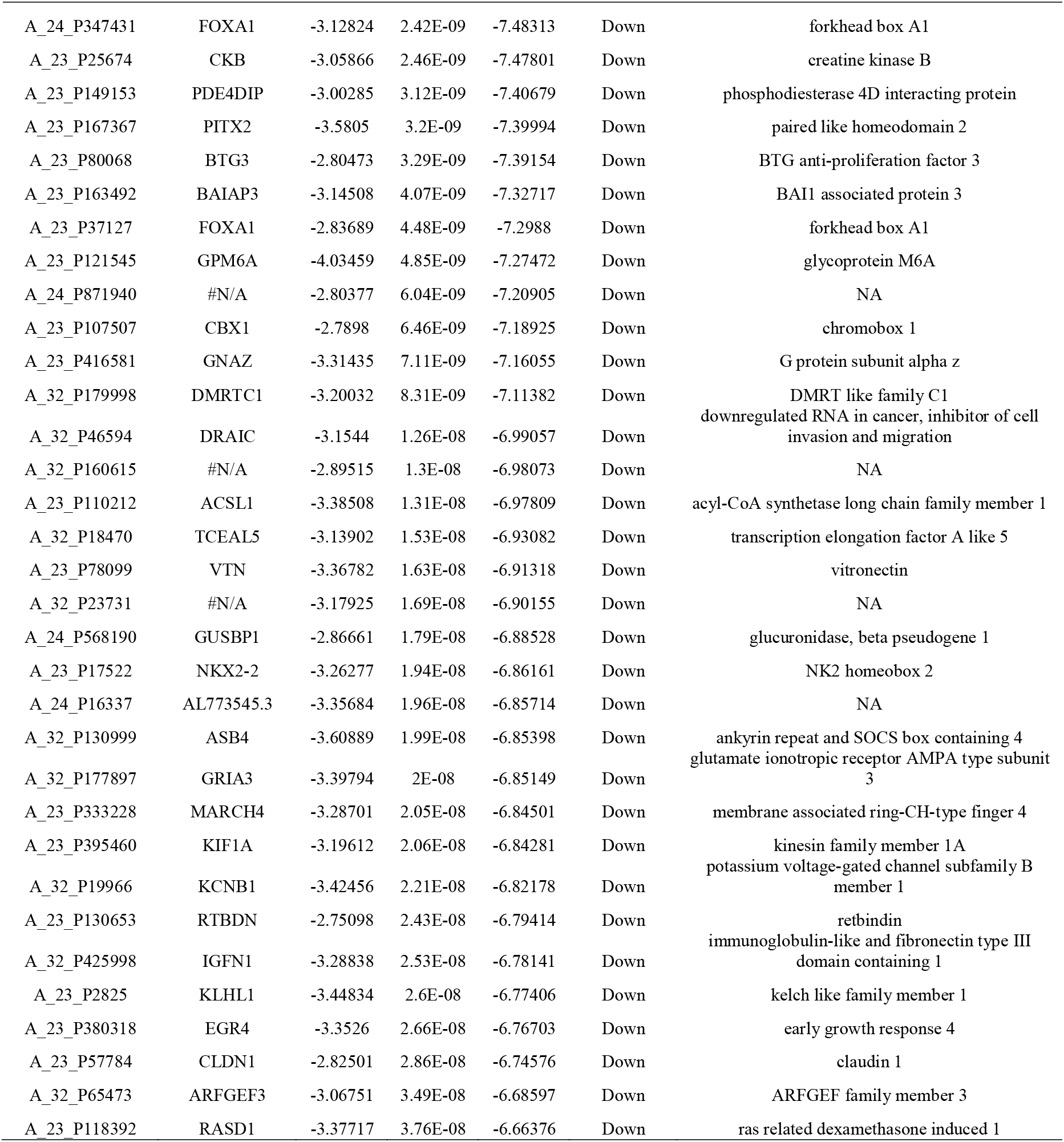
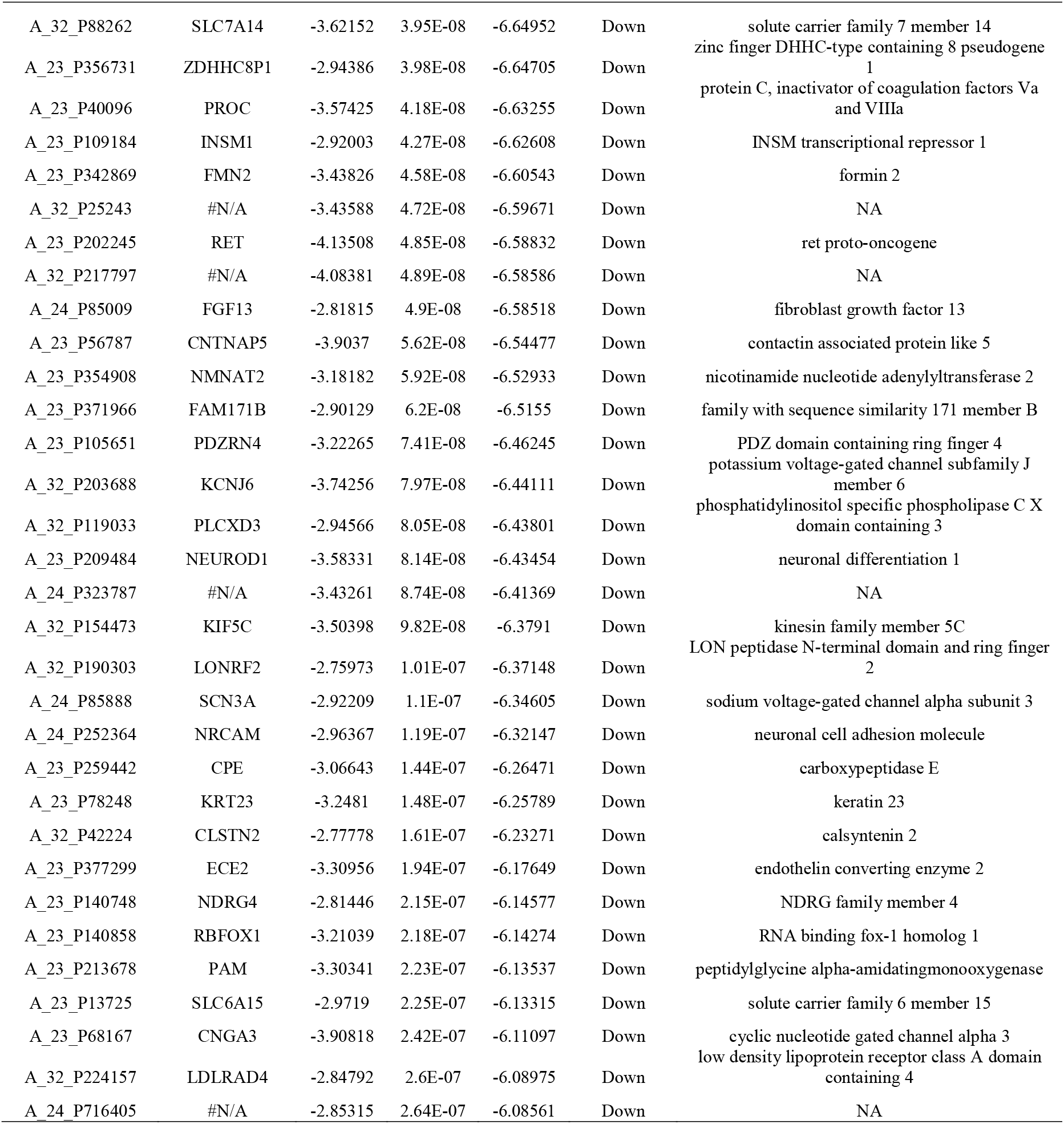
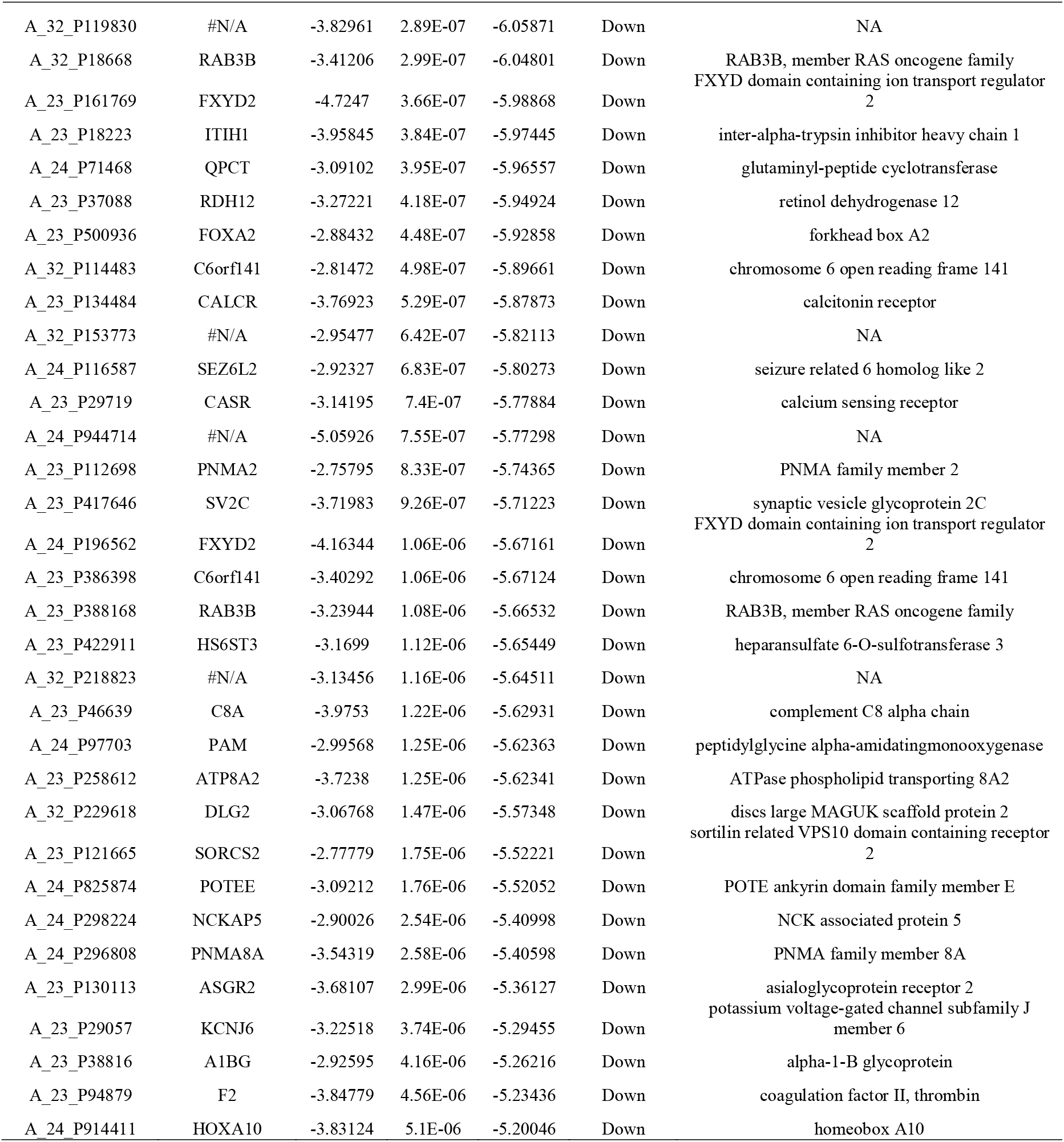
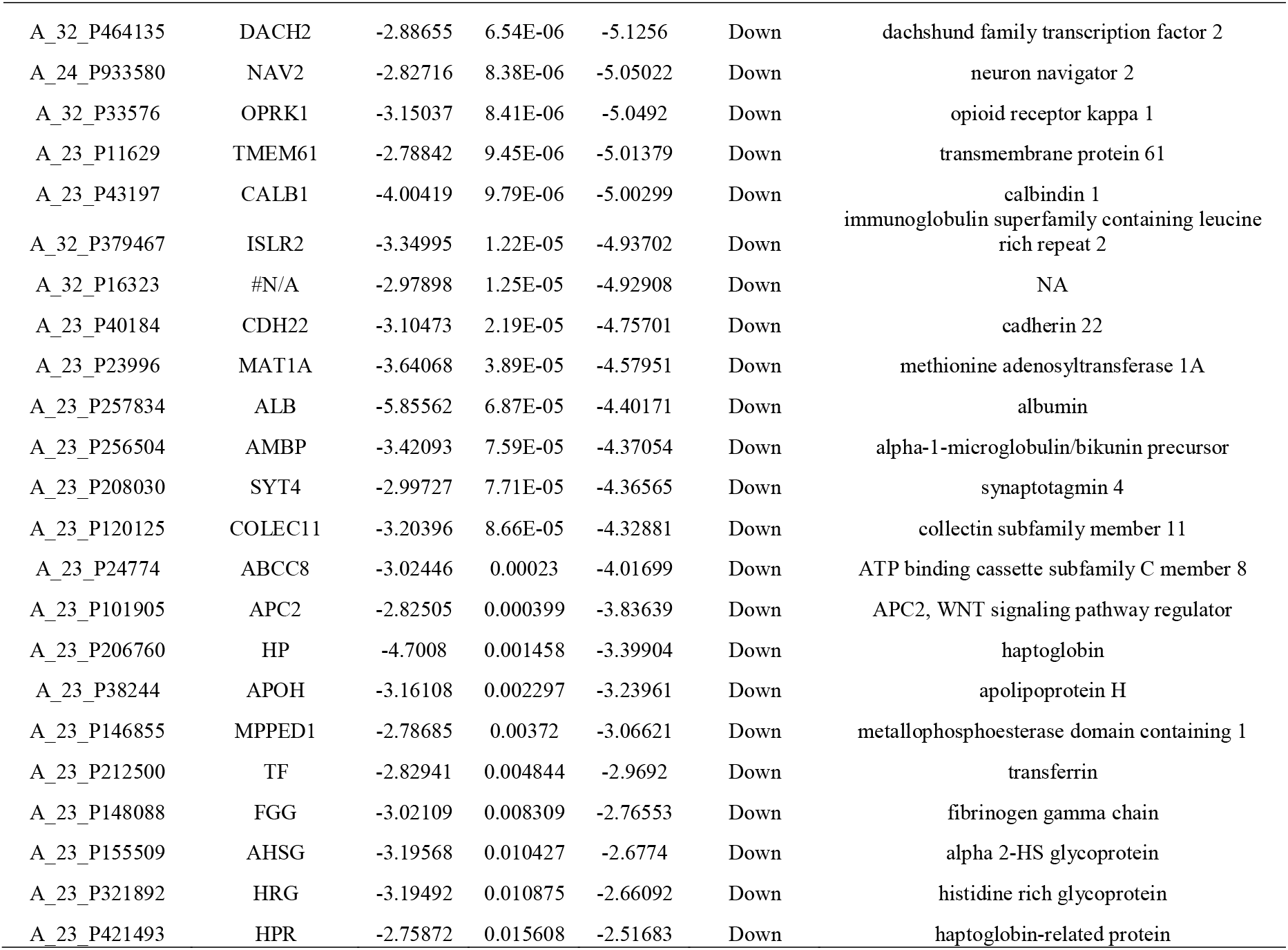
The statistical metrics for key differentially expressed genes (DEGs)

### Pathway enrichment analysis

To further analyze the biological function of the DEGs we processed pathway enrichment analysis with ToppCluster. As we conducted the pathway enrichment analysis, we found that these up and down regulated genes were mainly enriched in sucrose degradation, glutathione-mediated detoxification, chemical carcinogenesis, carbohydrate digestion and absorption, FOXA2 and FOXA3 transcription factor networks, a6b1 and a6b4 integrin signaling, transmembrane transport of small molecules, glutathione conjugation, Blood group glycolipid biosynthesis lact series, glycerolipid metabolism, genes encoding proteins affiliated structurally or functionally to extracellular matrix proteins, mechanism of gene regulation by peroxisome proliferators via PPARa (alpha), aginine biosynthesis, 2-arachidonoylglycerol biosynthesis, lipoprotein metabolic, renin-angiotensin system signaling, gluconeogenesis, arginine and proline metabolism, creatine biosynthesis, S-adenosyl-L-methionine biosynthesis, insulin secretion, complement and coagulation cascades, FOXA2 and FOXA3 transcription factor networks, FOXA transcription factor networks, platelet degranulation, cardiac conduction, urea cycle and metabolism of amino groups, extrinsic prothrombin activation pathway, acute myocardial infarction, blood coagulation, ionotropic glutamate receptor, insulin secretion pathway, coagulation cascade, lidocaine (antiarrhythmic) pathway and tranexamic acid pathway The pathways with p value <0.05 are shown in Table 3 and Table 4.

**Table 3.**
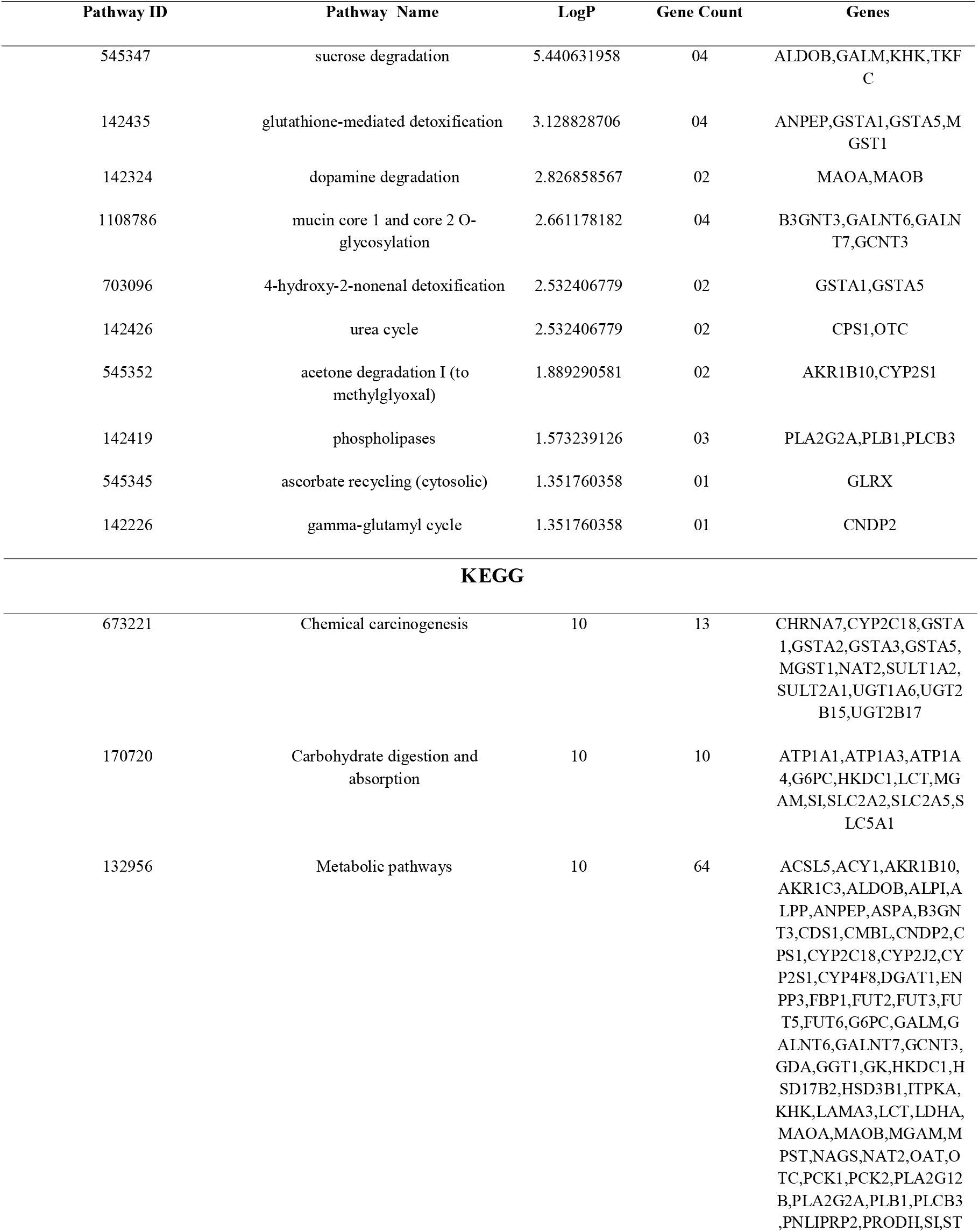
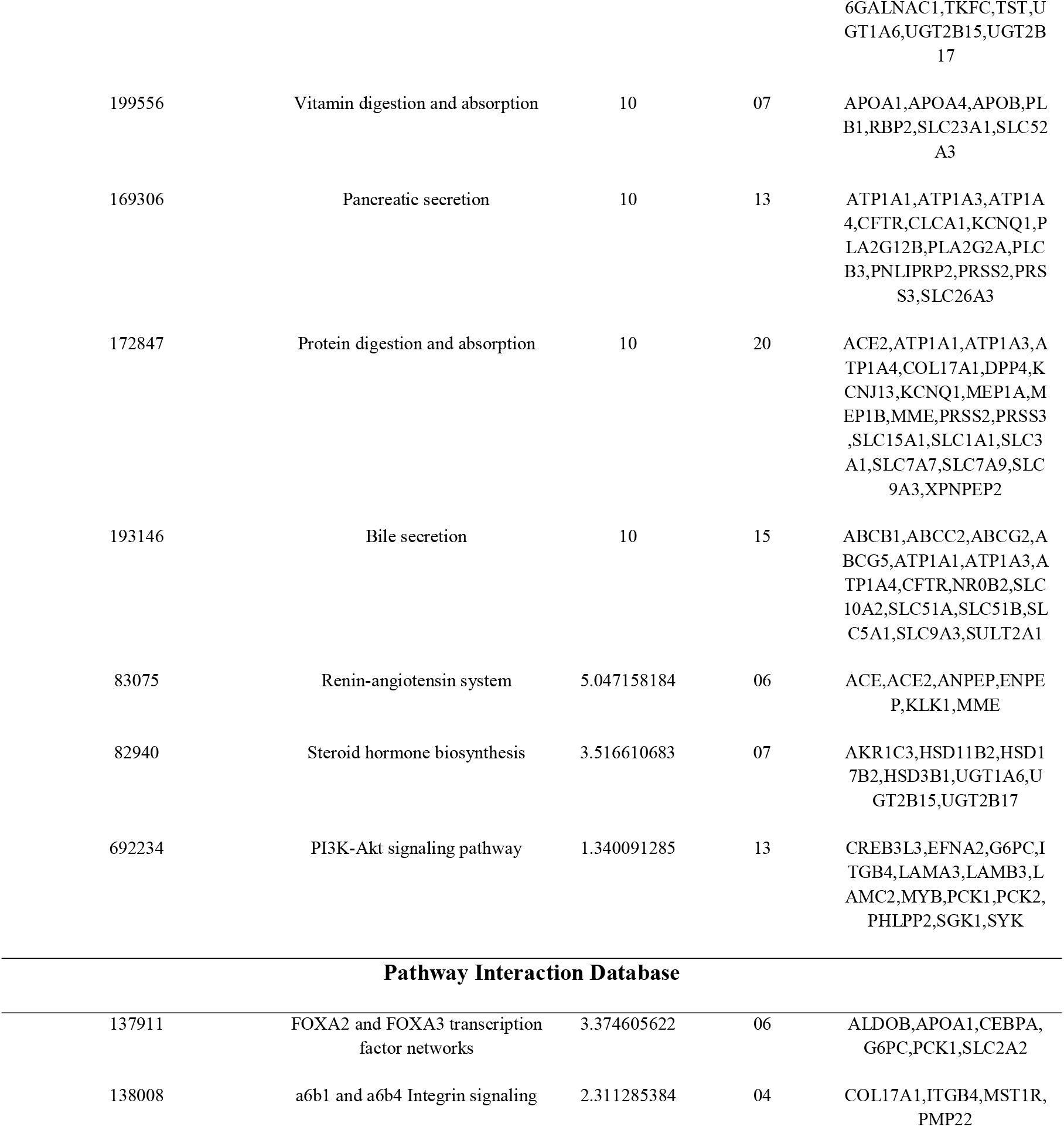
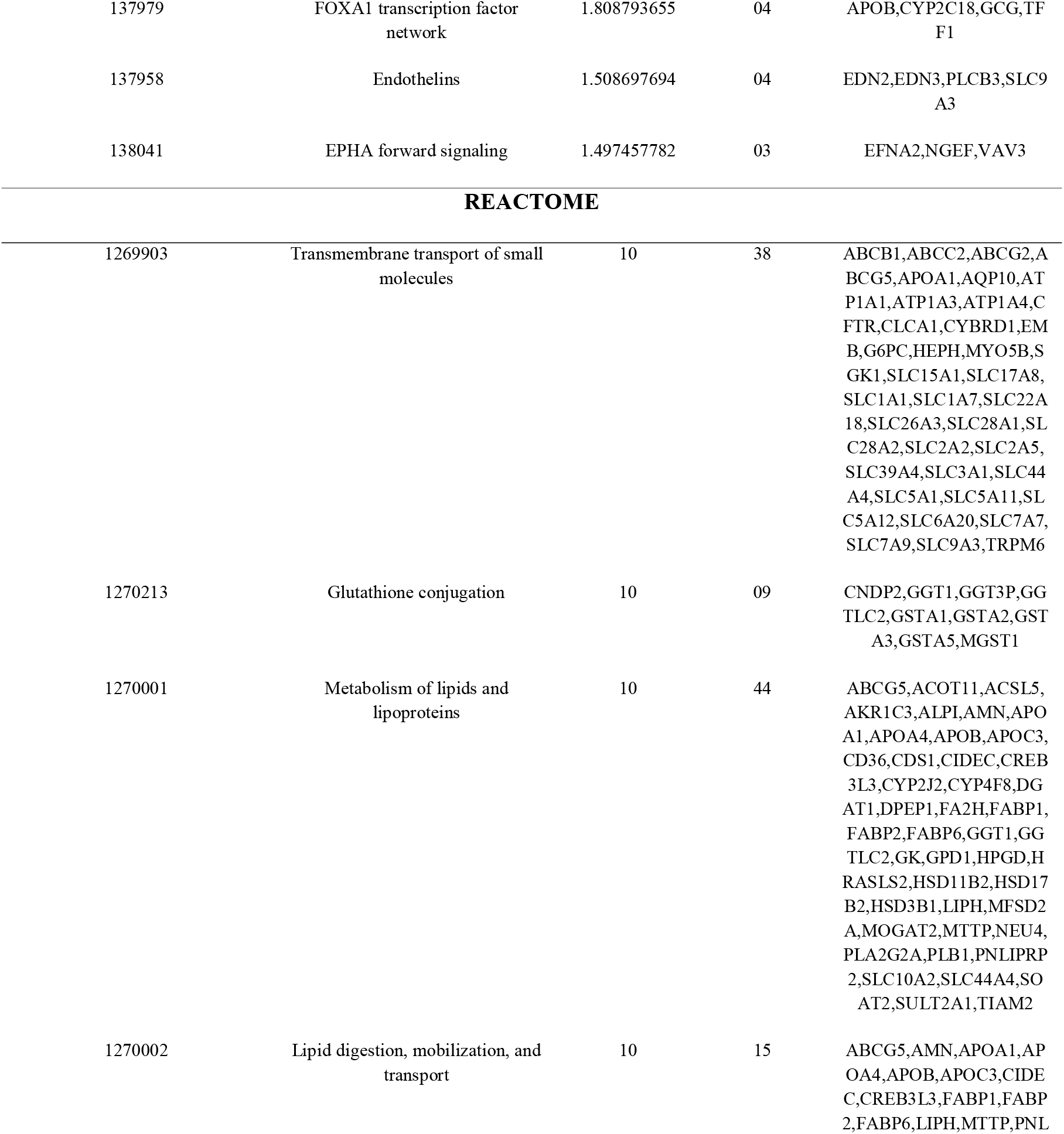
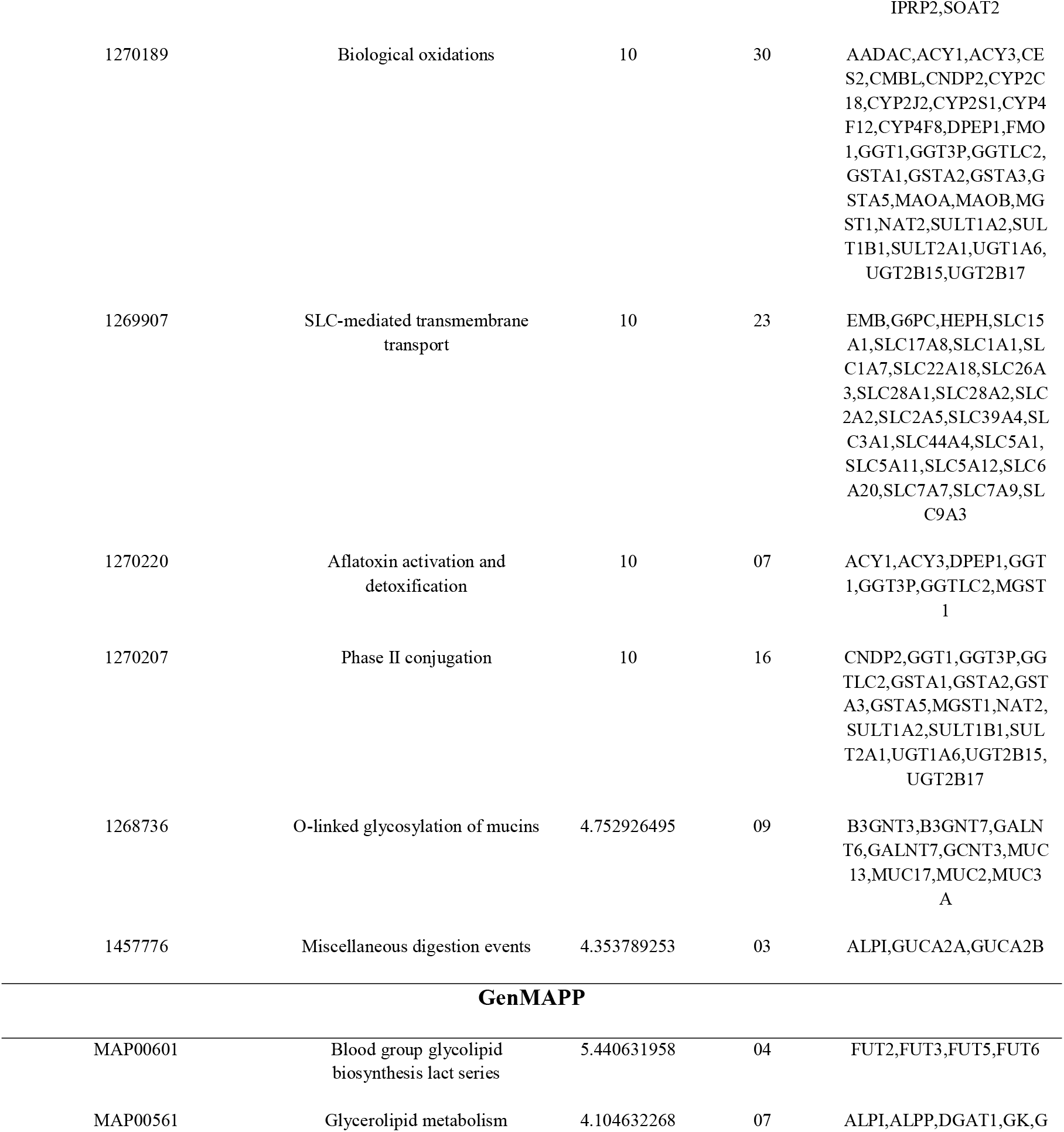
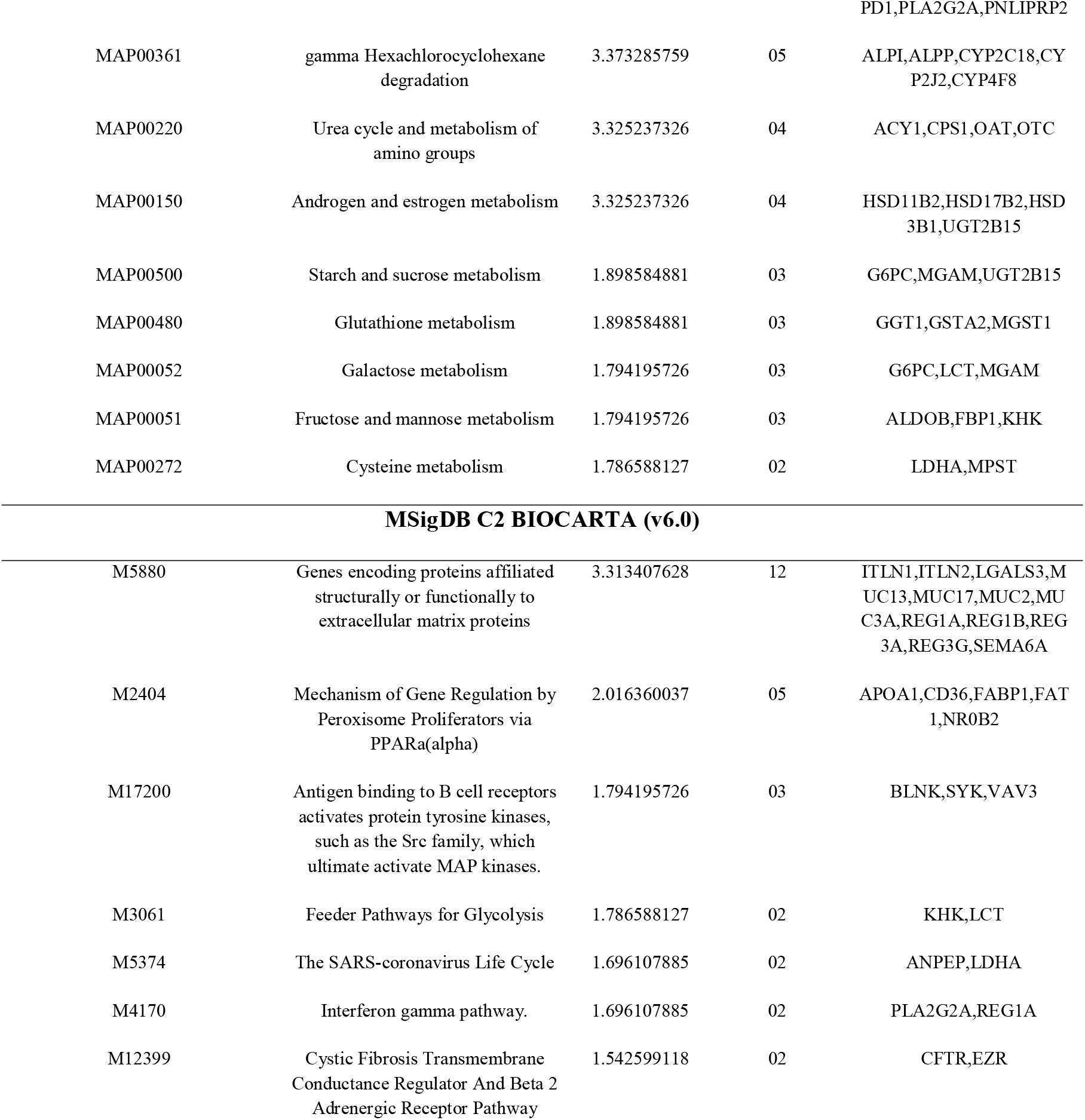
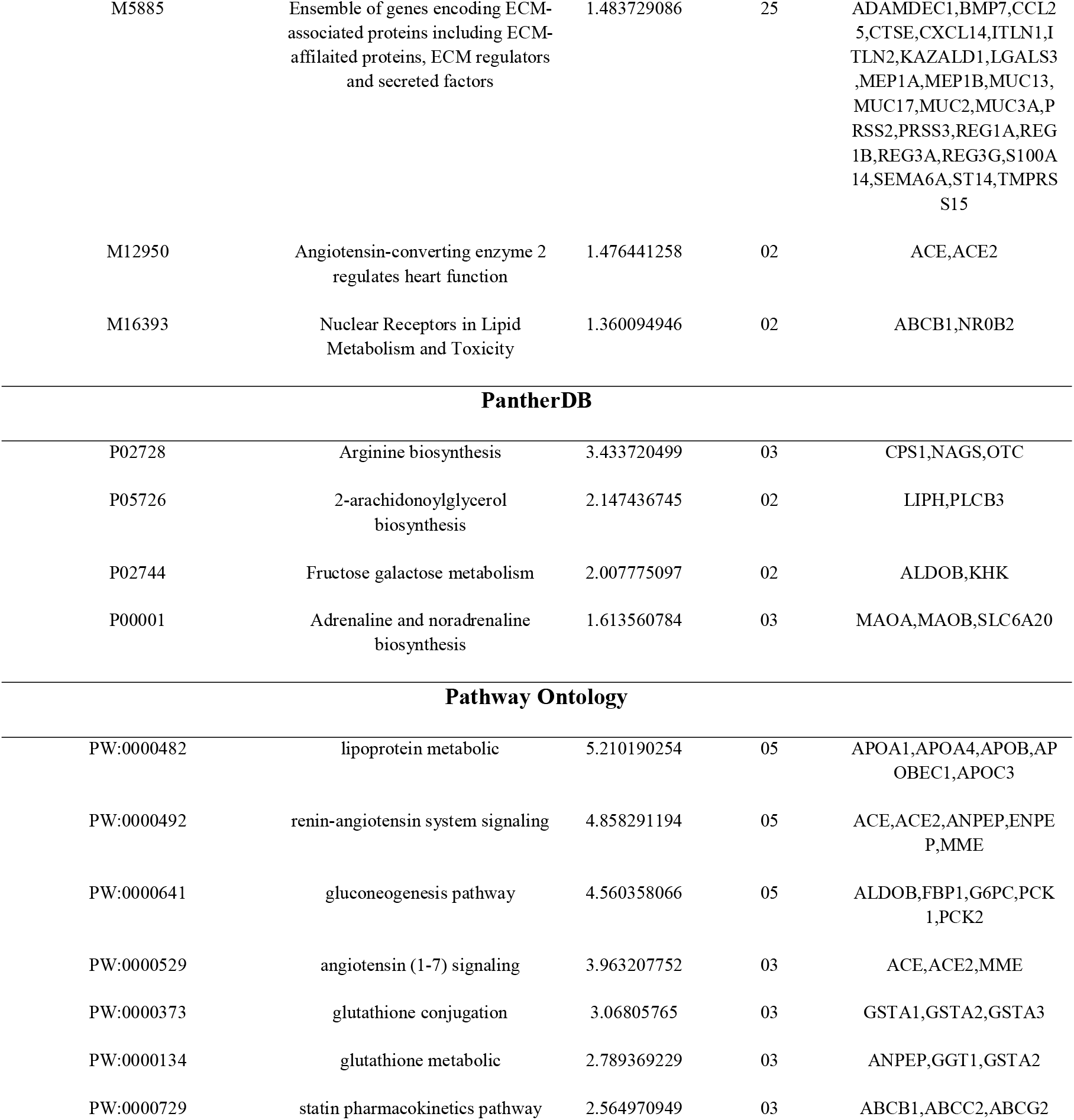
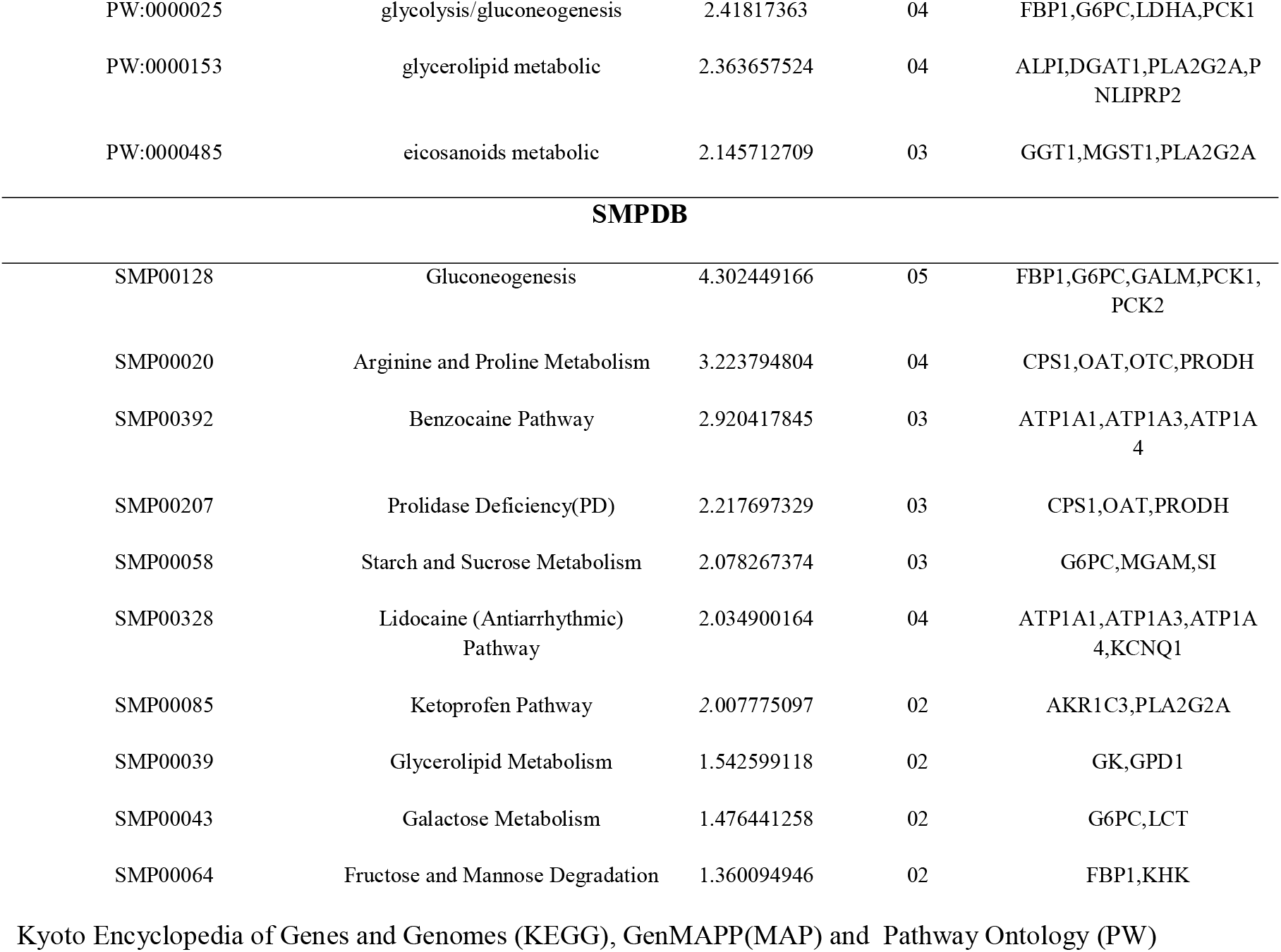
The enriched pathway terms of the up regulated differentially expressed genes

**Table 4.**
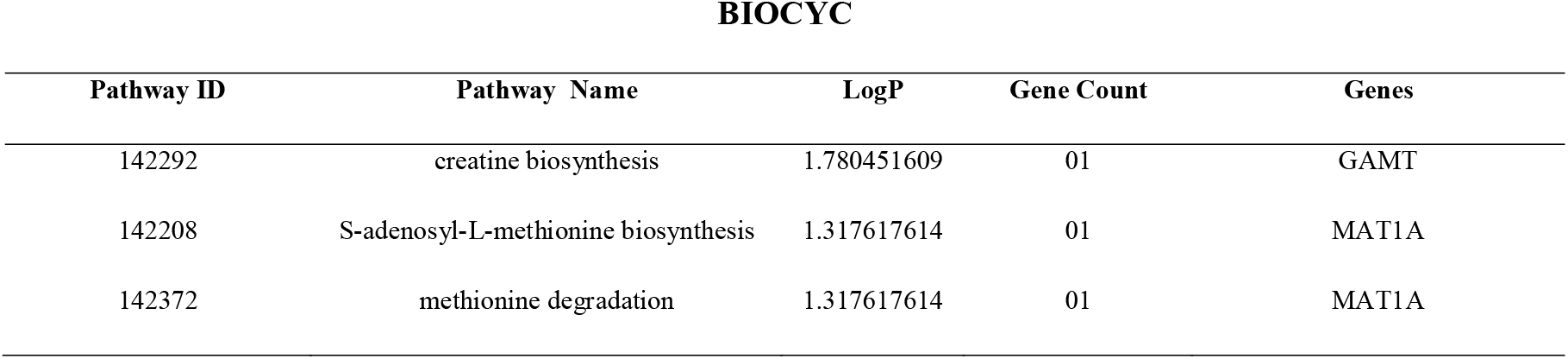
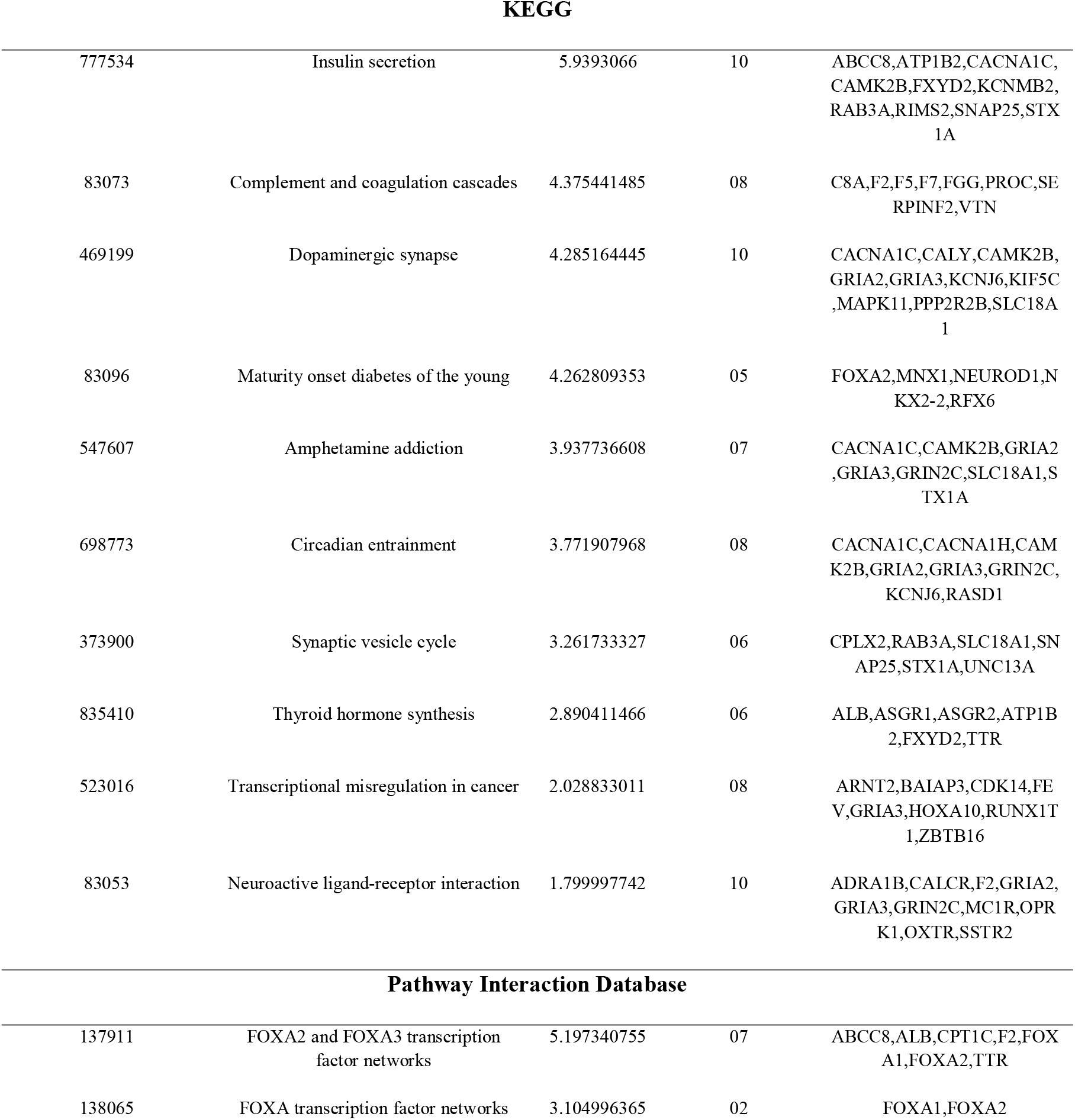
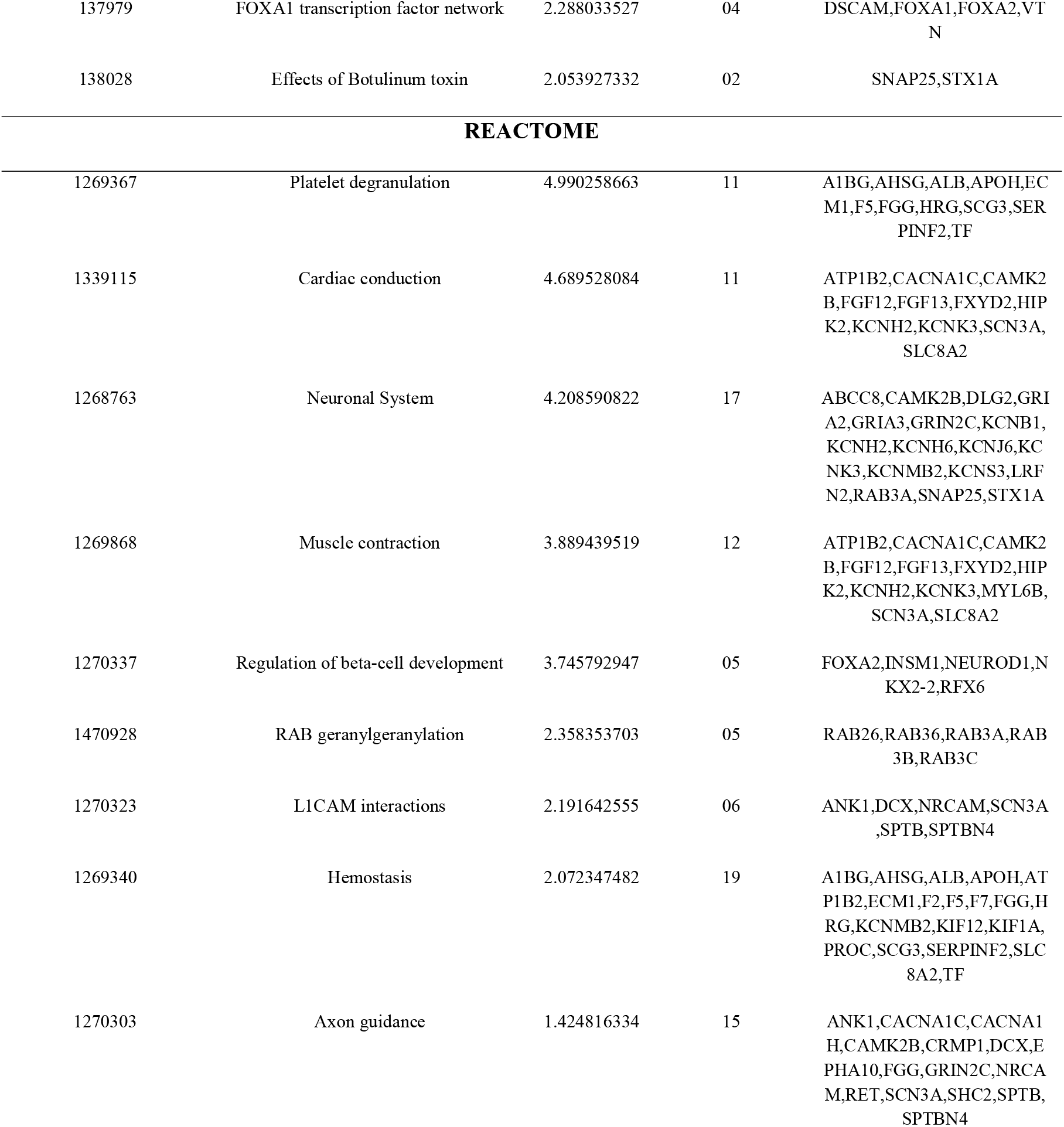
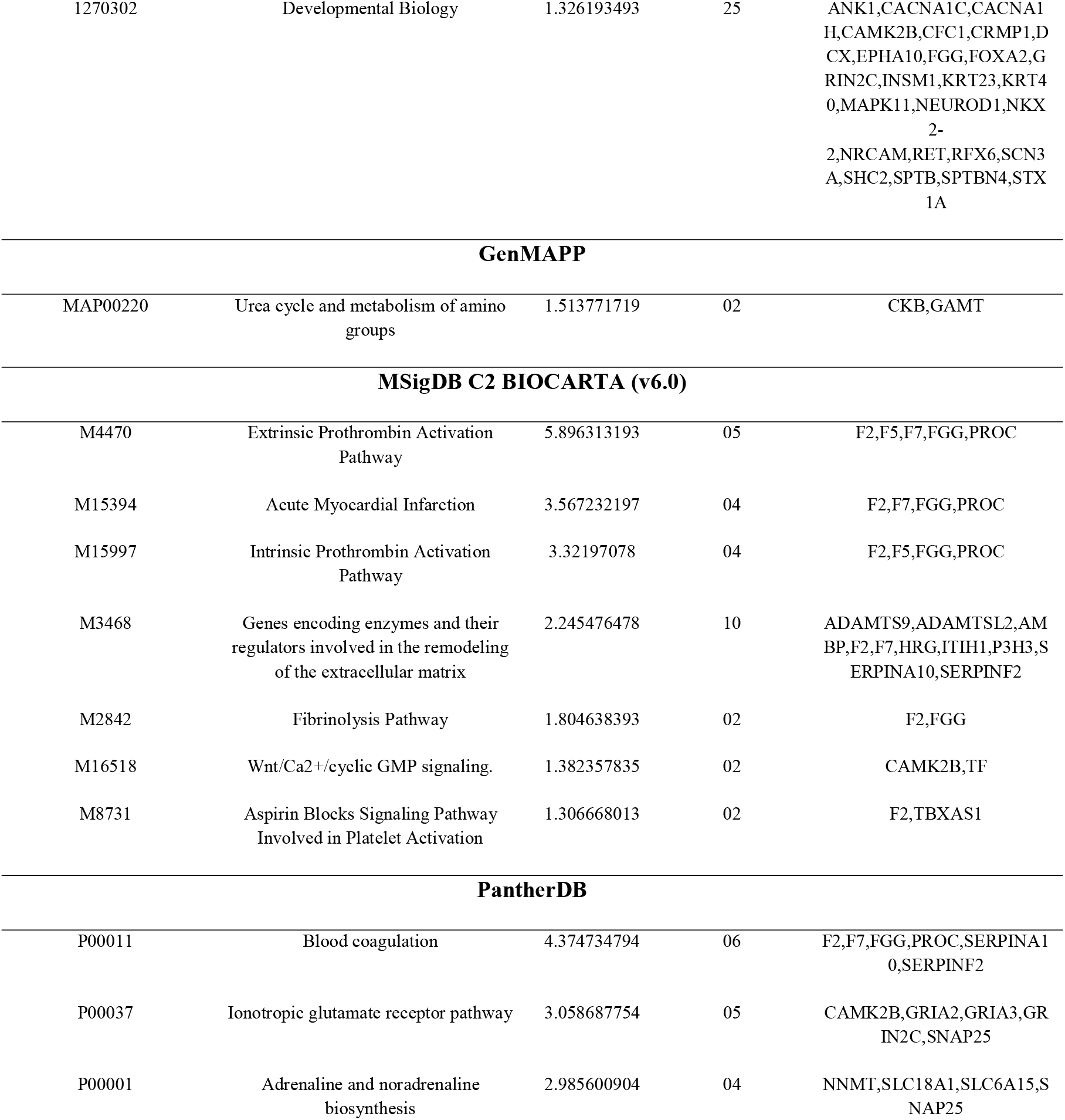
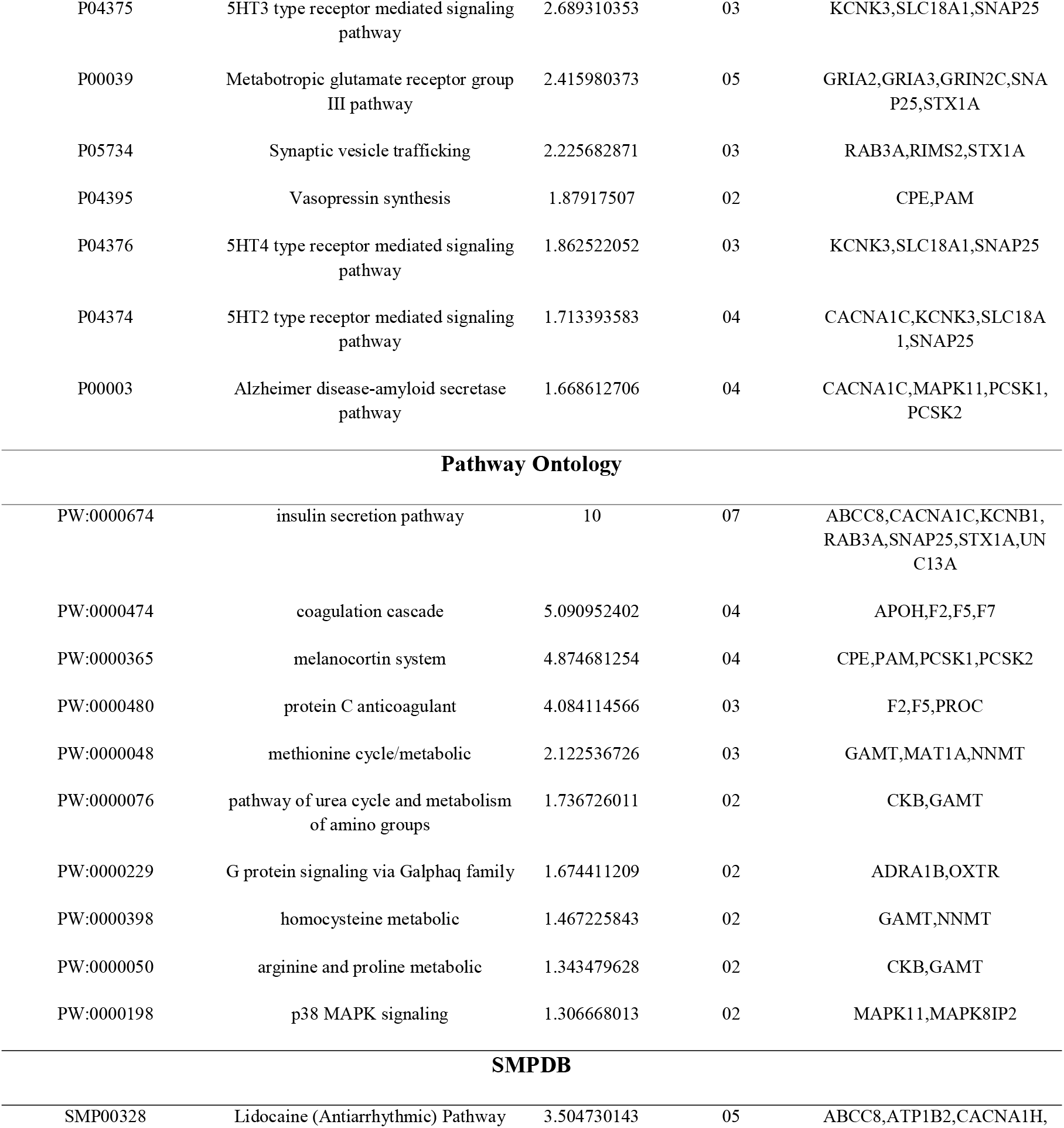
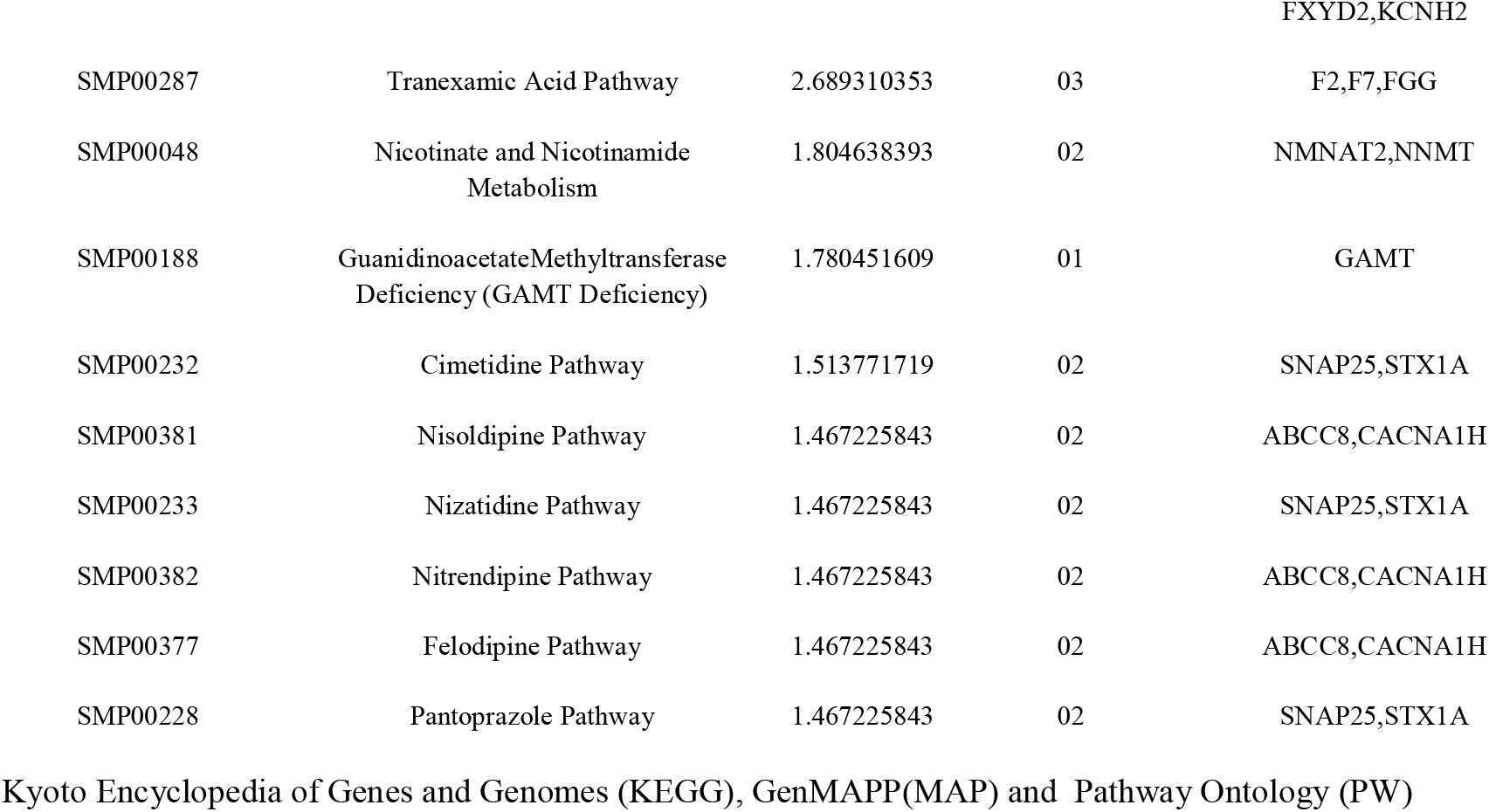
The enriched pathway terms of the down regulated differentially expressed genes

### Gene ontology (GO) enrichment analysis

The significant enrichment result of DEGs in BP, CC and MF are shown in Table 5 and Table 6. Up regulated genes were enriched in anion transport, cellular response to xenobiotic stimulus, cluster of actin-based cell projections, membrane region, transmembrane transporter activity and carbohydrate binding. Meanwhile, down regulated genes were enriched in modulation of chemical synaptic transmission, anterograde trans-synaptic signaling, secretory granule, vesicle membrane, cytoskeletal protein binding and metal ion transmembrane transporter activity.

**Table 5.**
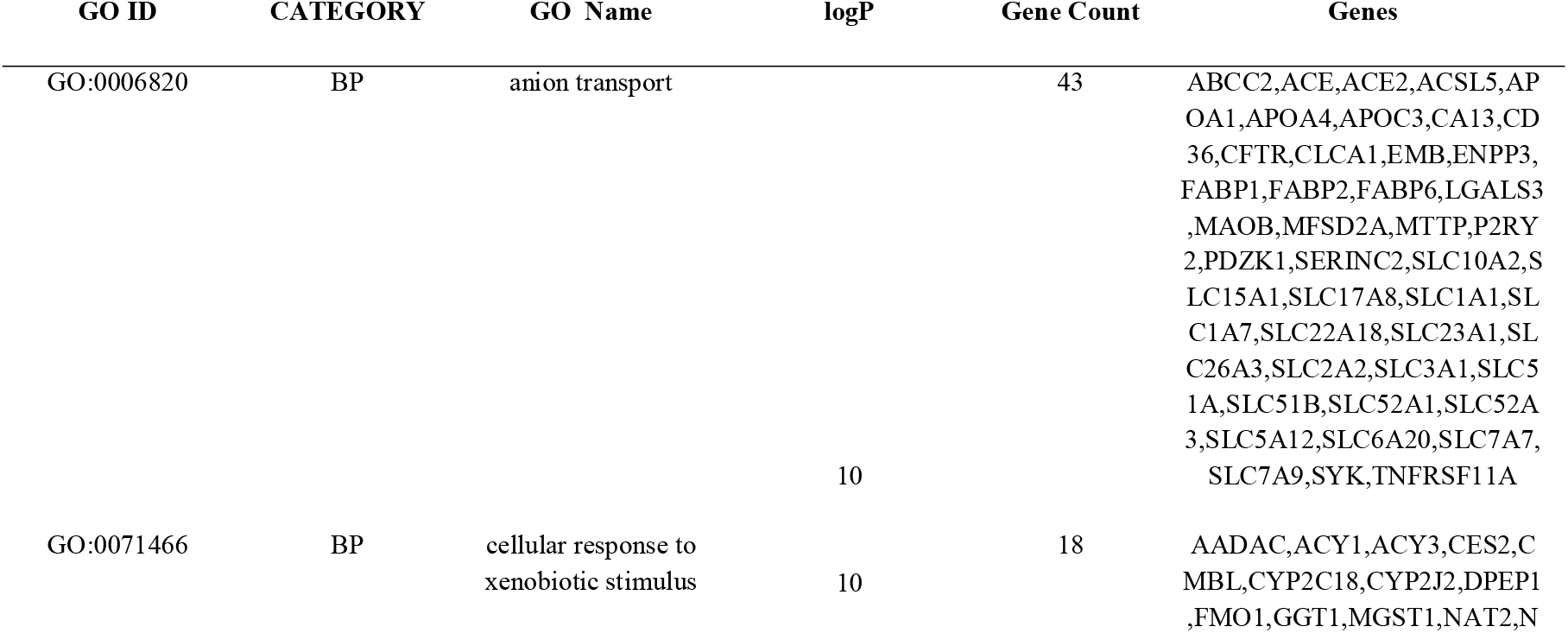
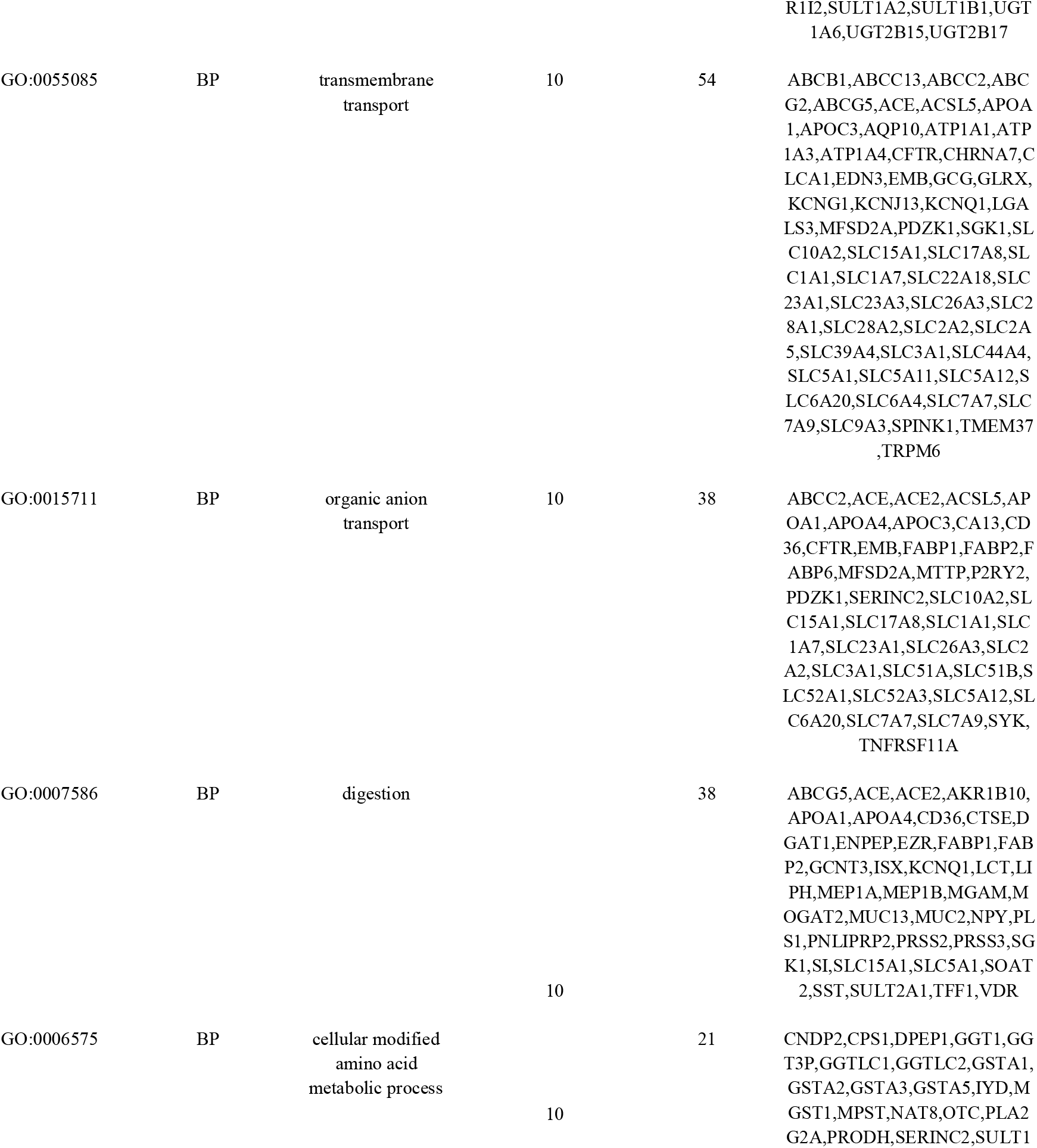
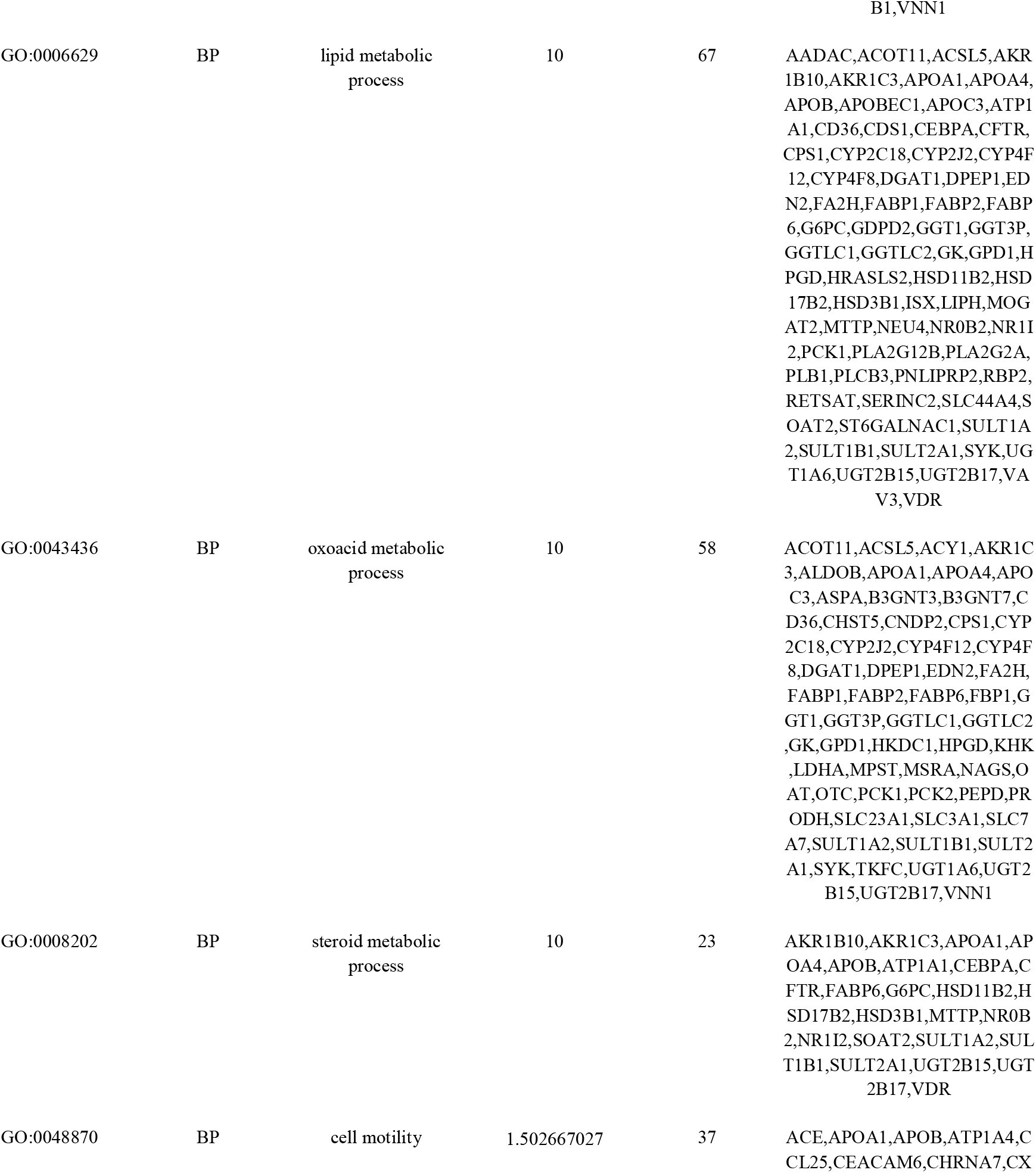
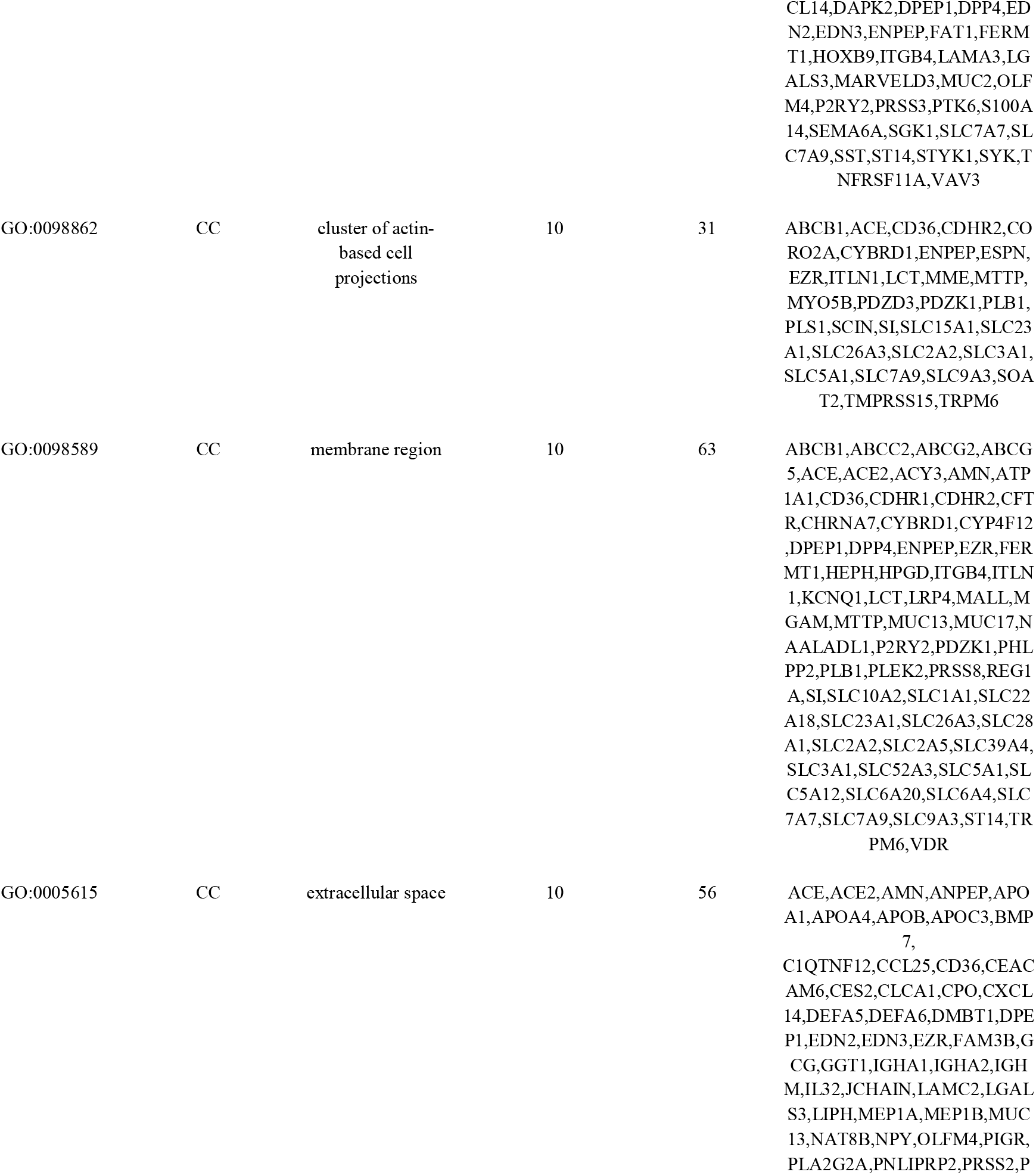
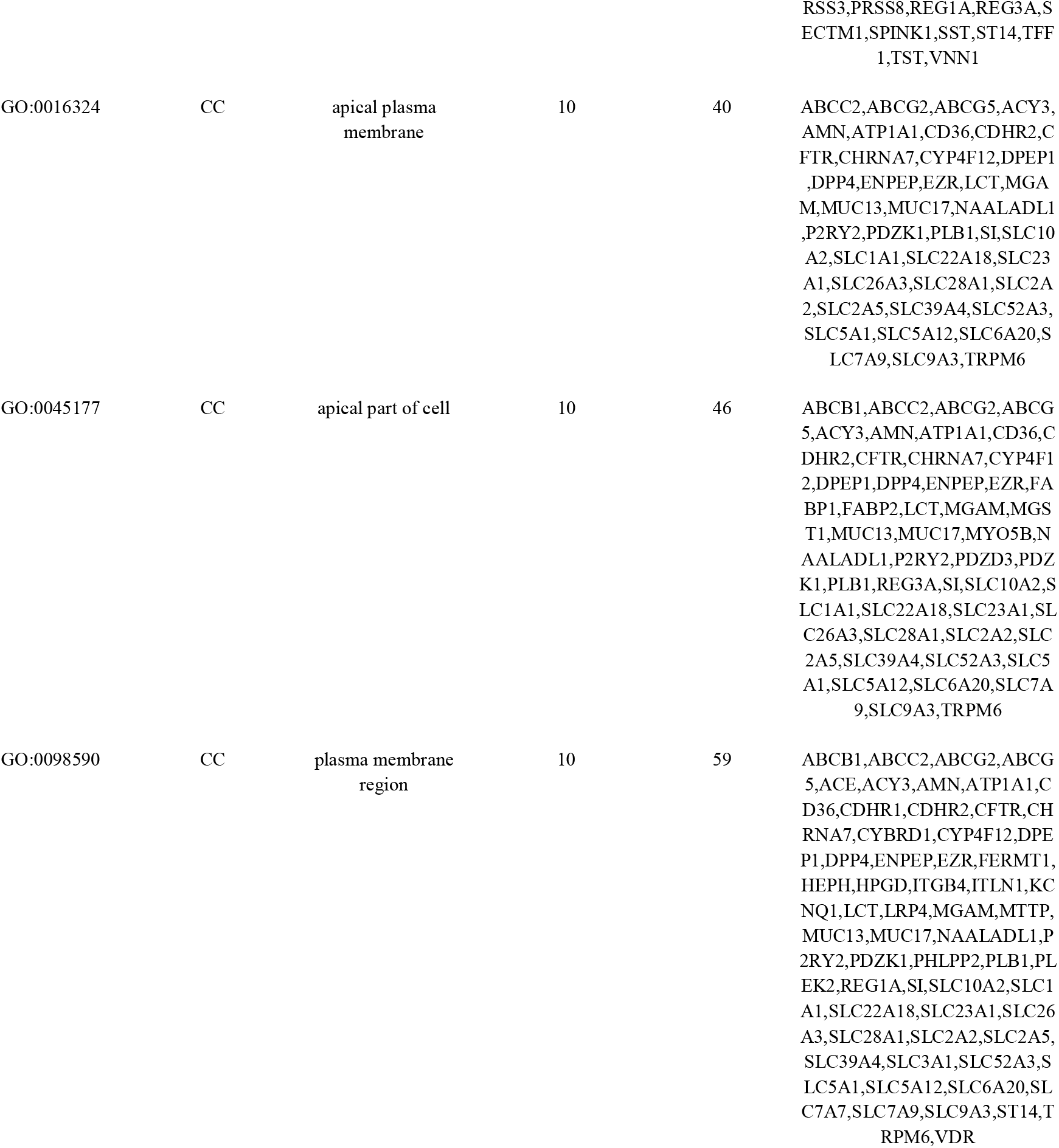
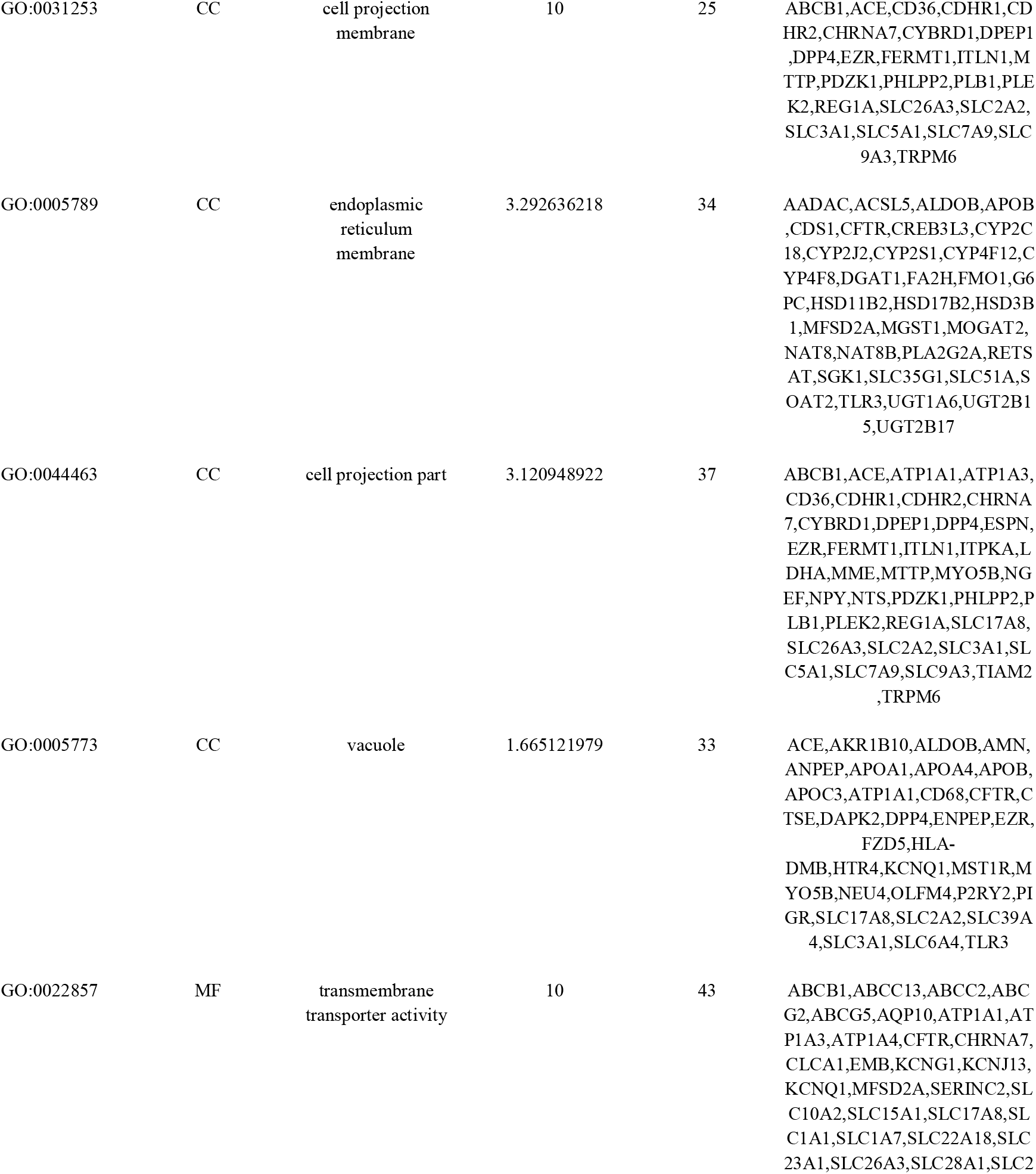
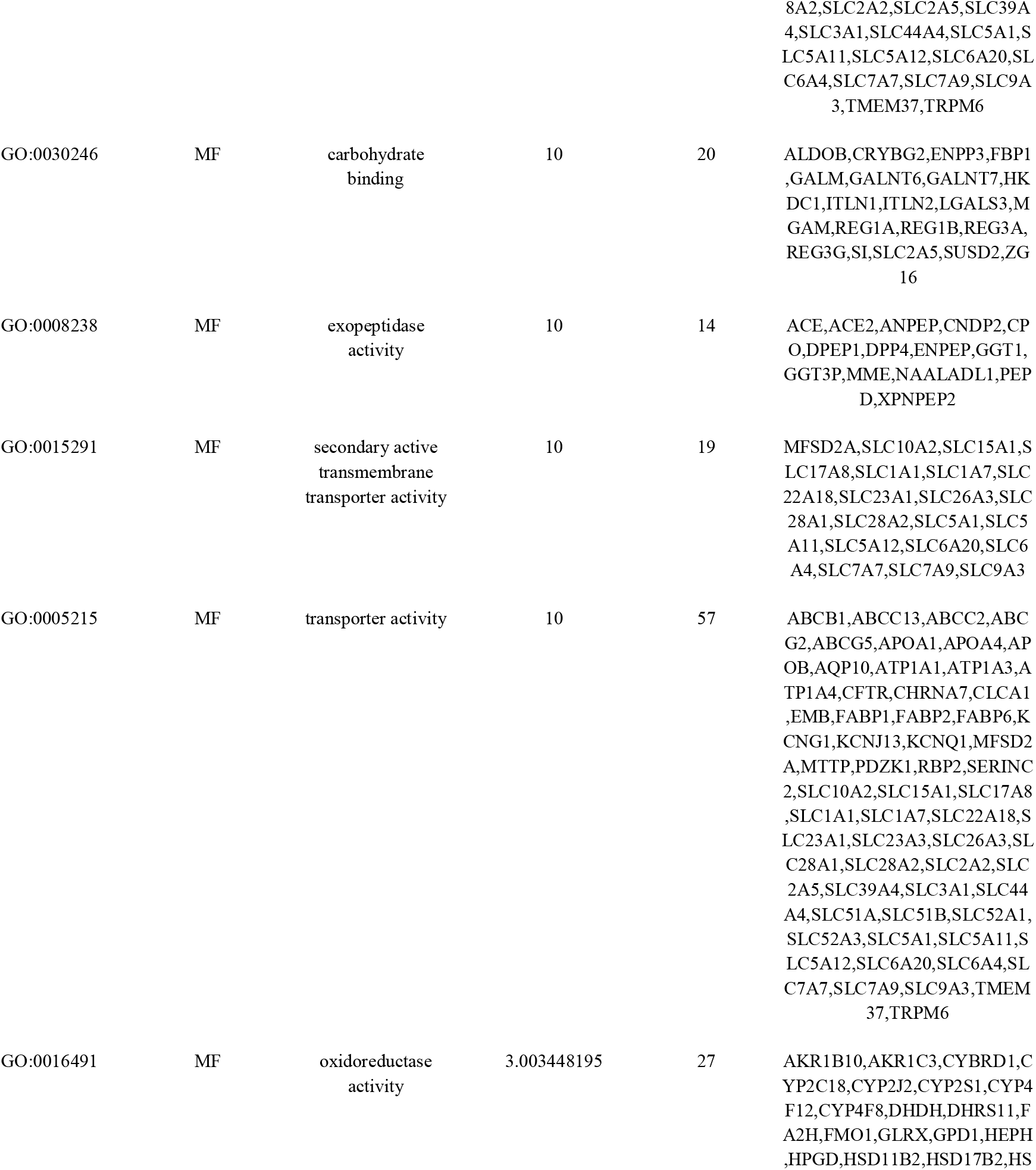
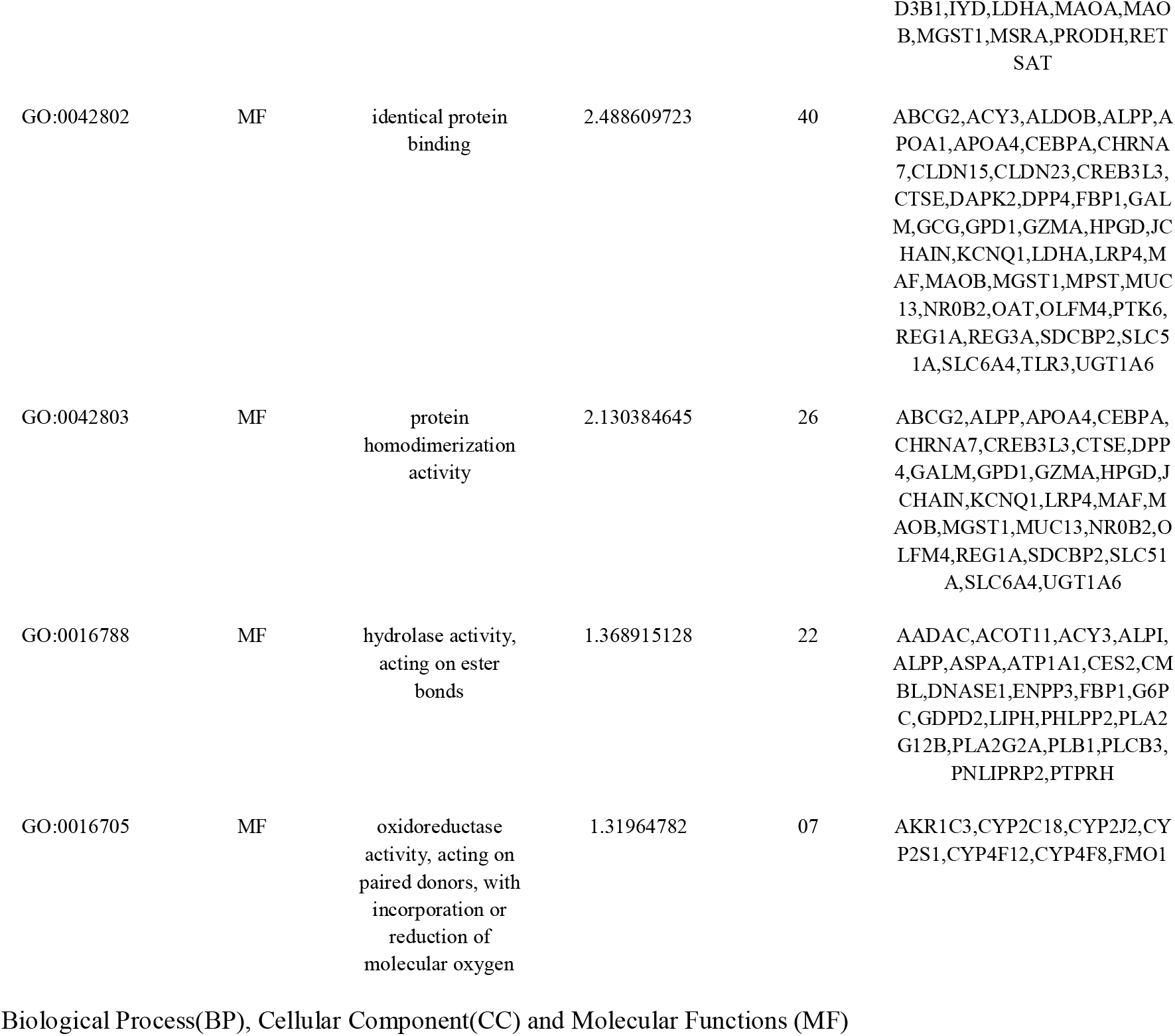
The enriched GO terms of the up-regulated differentially expressed genes

**Table 6.**
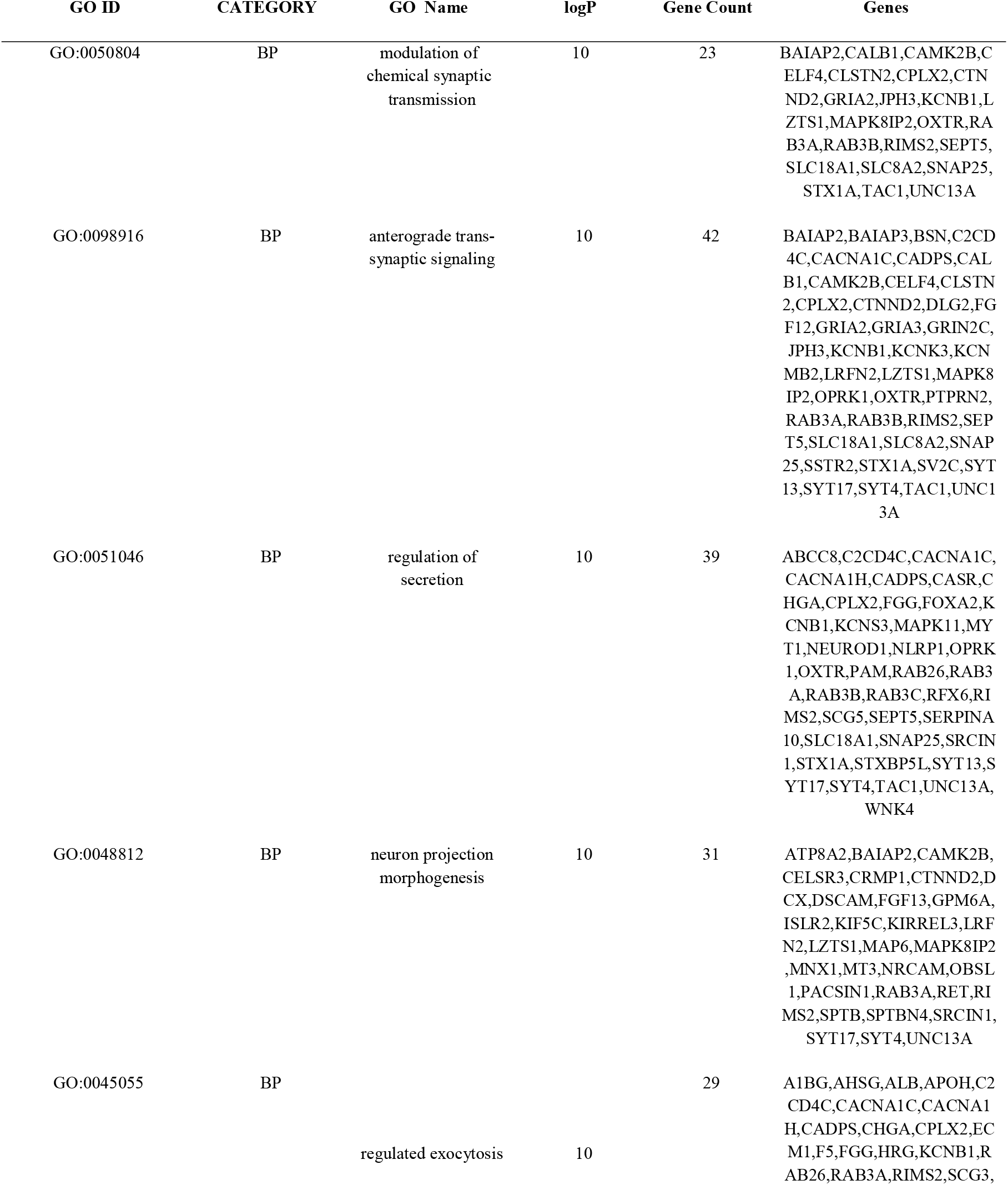
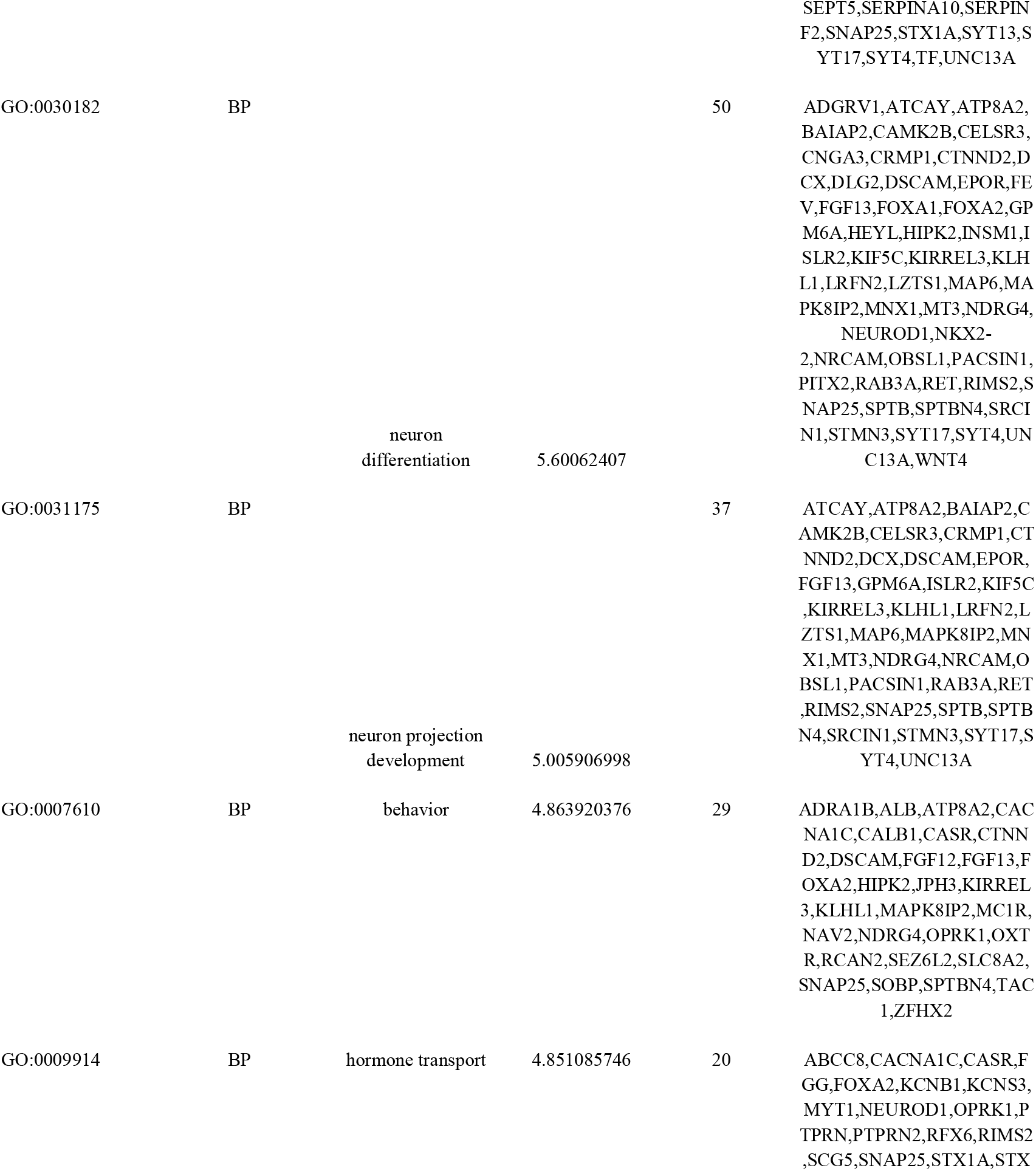
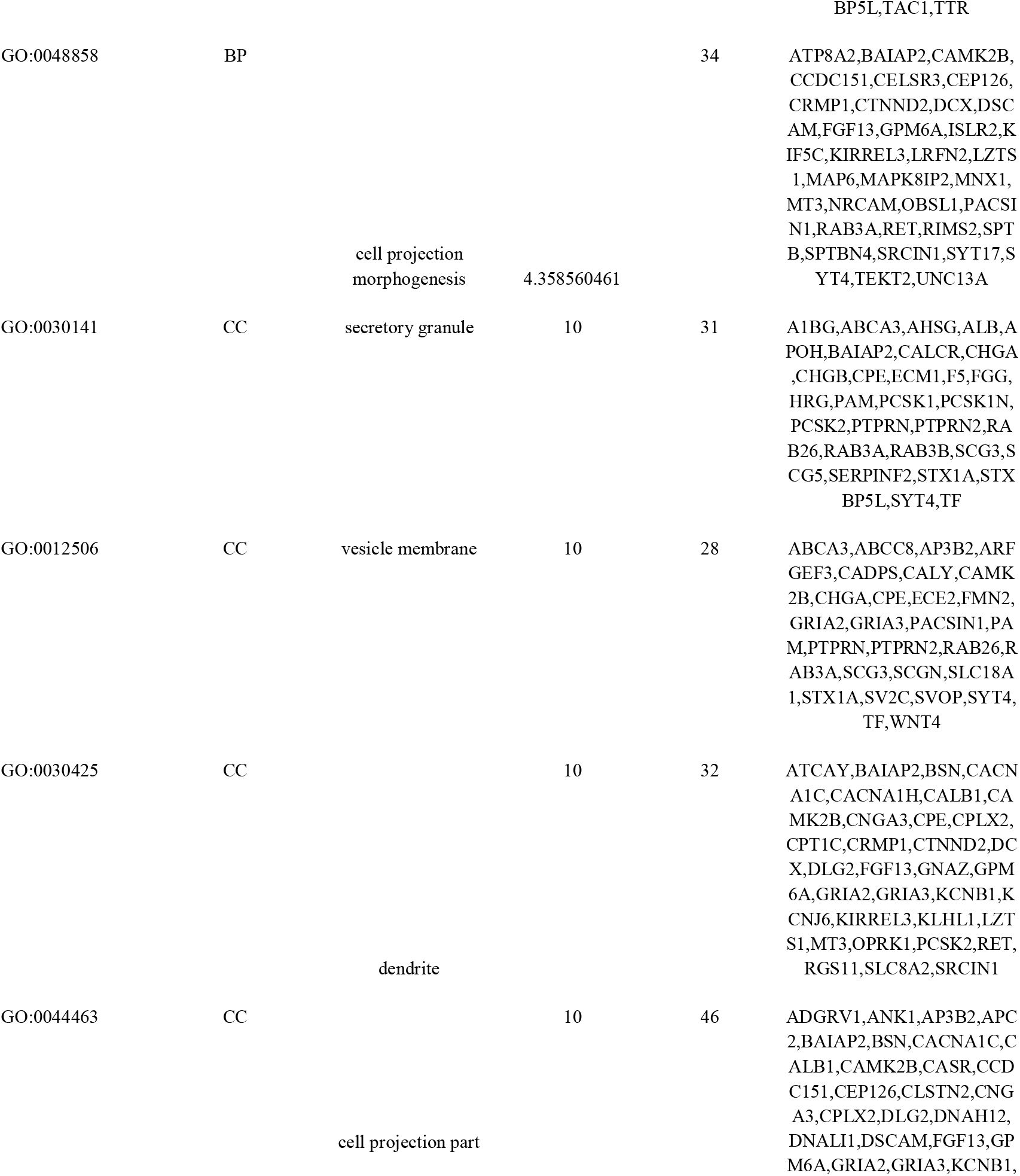
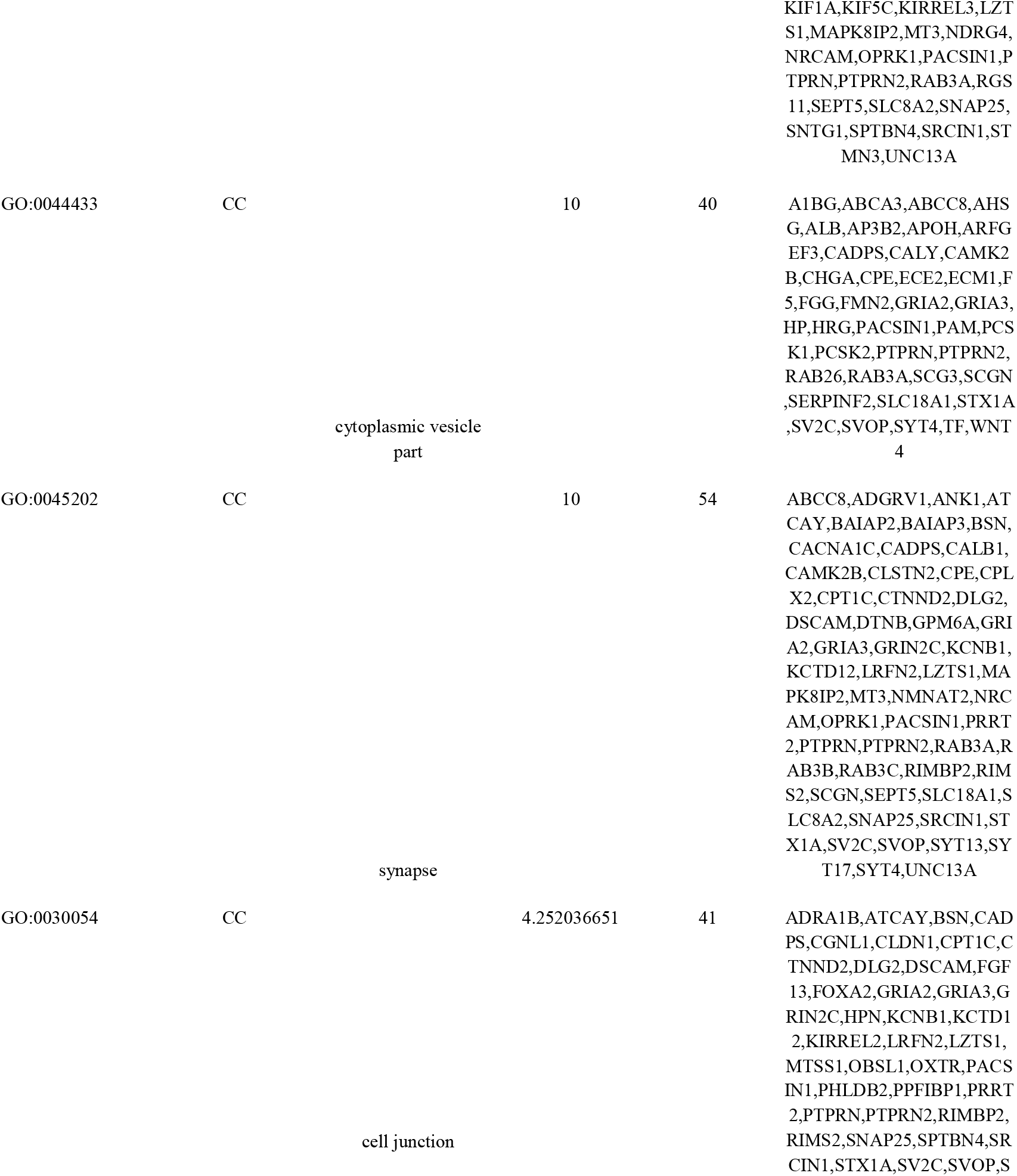
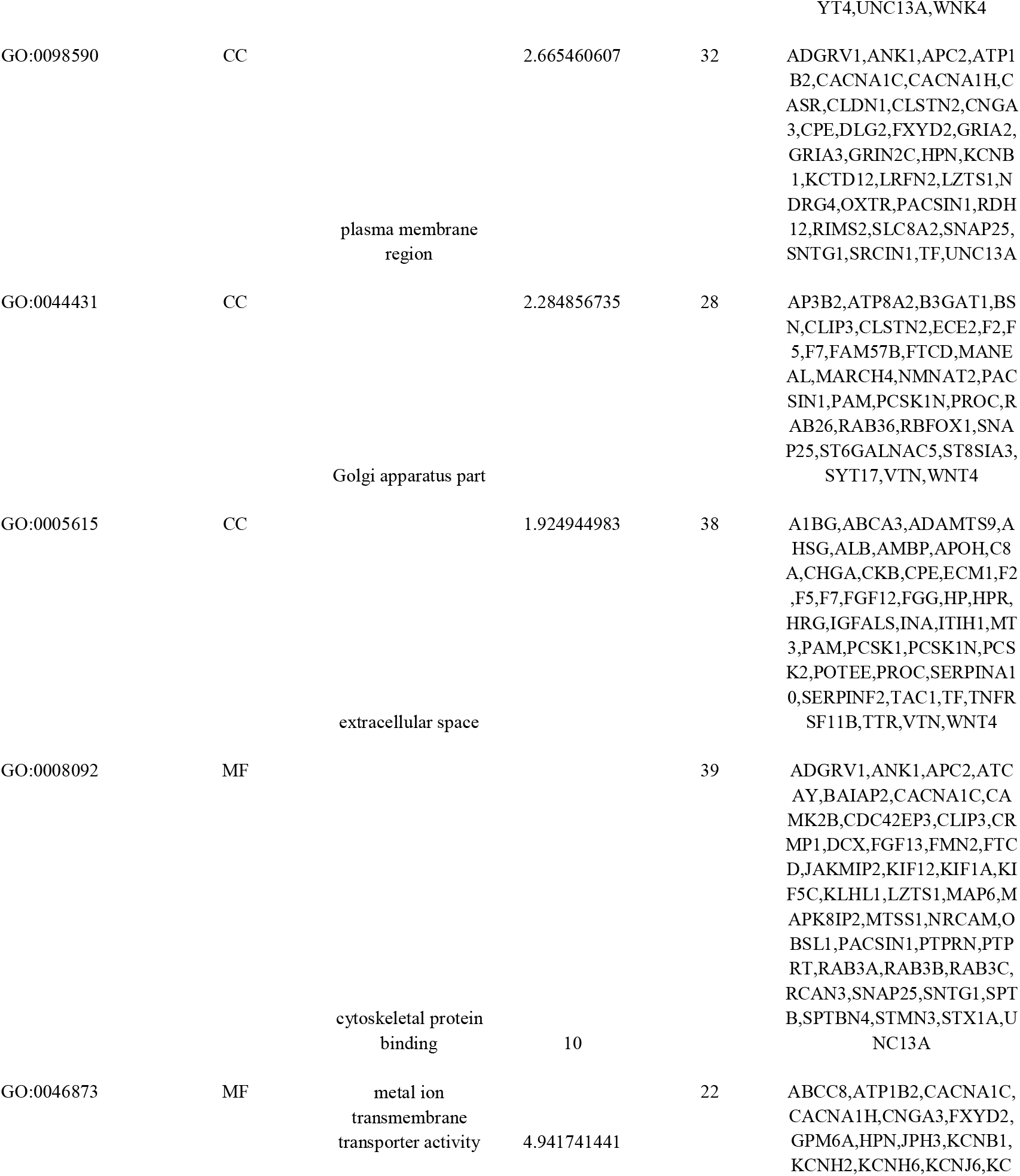
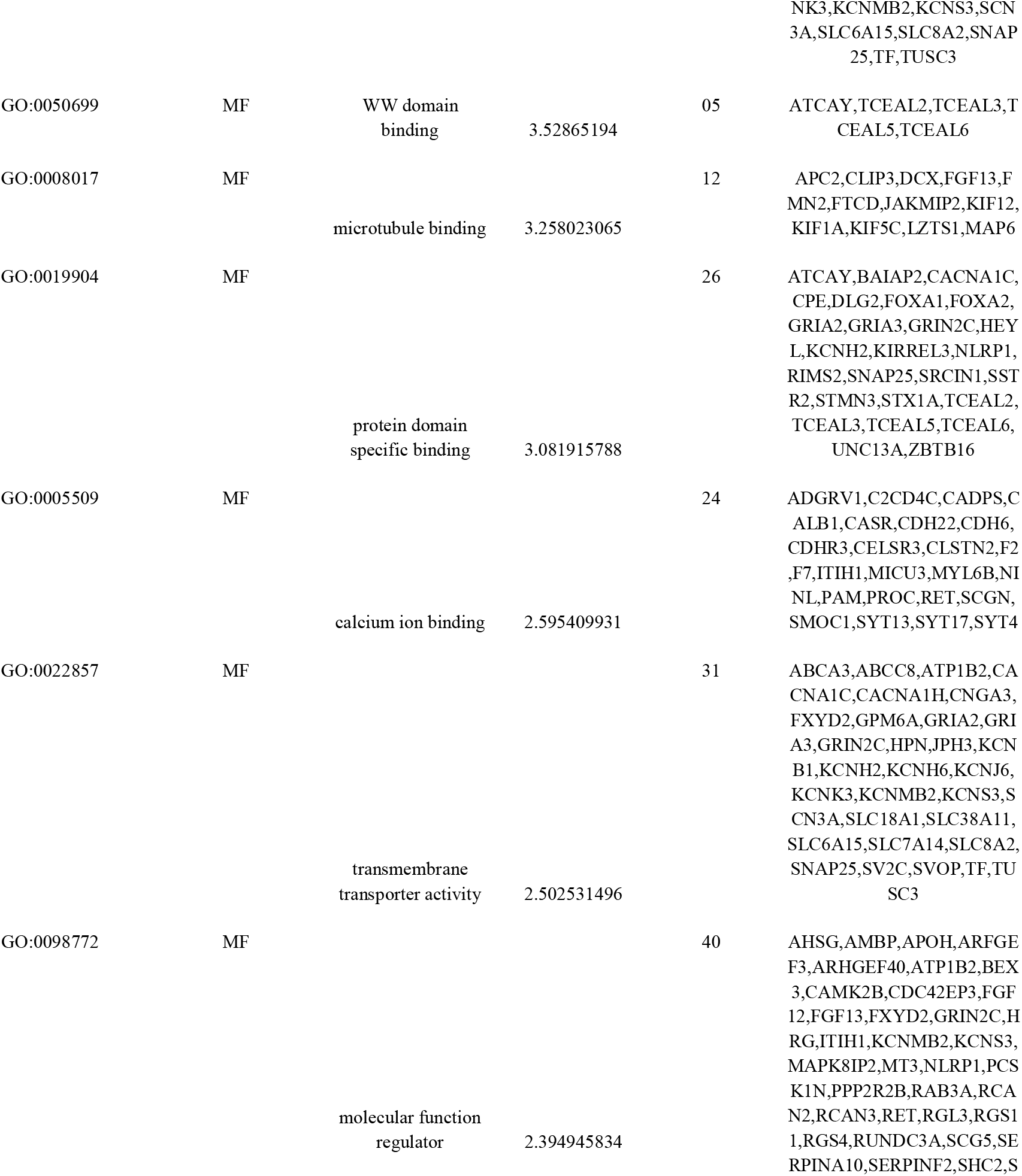
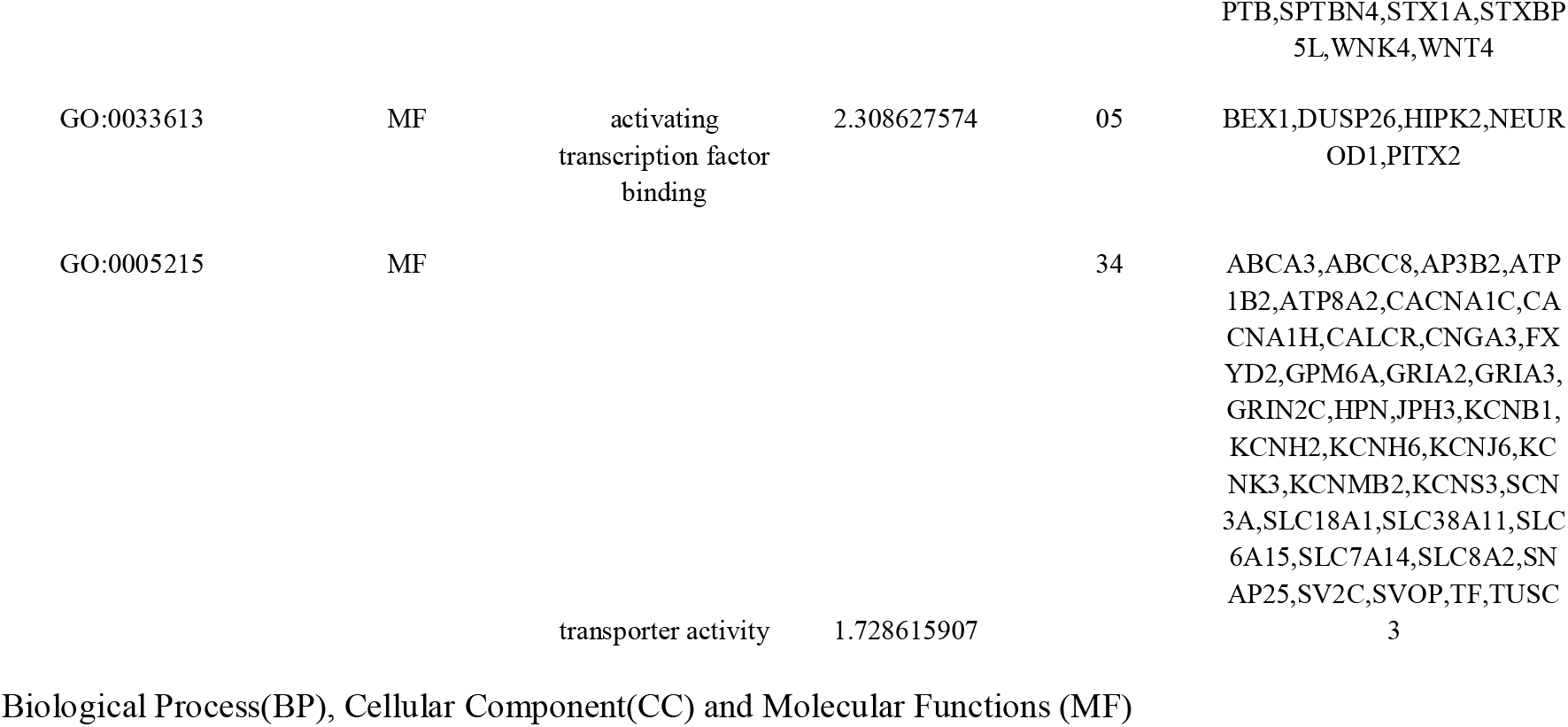
The enriched GO terms of the down-regulated differentially expressed genes

### Protein–protein interaction (PPI) network construction and module analysis

The constructed PPI network of up regulated gene is shown in Fig. 4A. In total, 4933 nodes and 8183 interacting protein pairs were contained in the PPI network. The up regulated hub genes were ranked by node degree distribution, betweenness centrality, stress centrality, closeness centrality and clustring coefficient. CFTR, EZR, ATP1A1, TUBA1C, LGALS3, C15orf48, LDHA, G6PC, ACOT11, PLA2G12B, HSD17B2 and DHDH were considered as up regulated hub genes in NET are listed in Table 7. These up regulated hub genes enriched in pancreatic secretion, cystic fibrosis transmembrane conductance regulator and beta 2 adrenergic receptor pathway, carbohydrate digestion and absorption, genes encoding proteins affiliated structurally or functionally to extracellular matrix proteins, metabolic pathways, carbohydrate digestion and absorption, metabolism of lipids and lipoproteins, lipid metabolic process, steroid hormone biosynthesis and oxidoreductase activity. Meanwhile, constructed PPI network of down regulated gene is shown in Fig. 4B. In total, 5307 nodes and 9099 interacting protein pairs were contained in the PPI network. The down regulated hub genes were ranked by node degree distribution, betweenness centrality, stress centrality, closeness centrality and clustring coefficient. KRT40, FOXA1, WWOX, OBSL1, ALB, PPP2R2B, CBX1, PHLDB2, HIPK2, SLC18A1, PNMA2 and ABCA3 are listed in Table 7. These down regulated hub genes enriched in developmental biology, FOXA2 and FOXA3 transcription factor networks, neuron projection morphogenesis, thyroid hormone synthesis, dopaminergic synapse, cell junction, cardiac conduction, adrenaline and noradrenaline biosynthesis and secretory granule.

**Figure 4.**
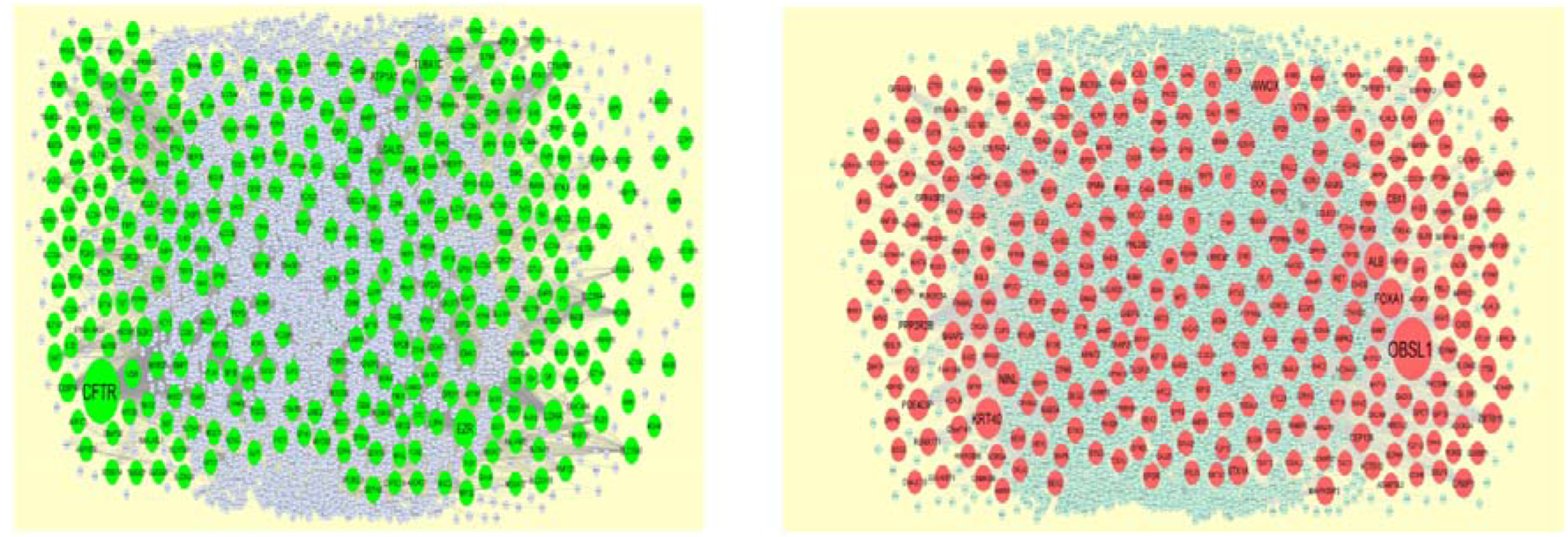
Protein–protein interaction network of (A) up regulated differentially expressed genes (B) down regulated differentially expressed genes. Green nodes denotes up regulated genes and red nodes denotes down regulated genes.

**Table 7.**
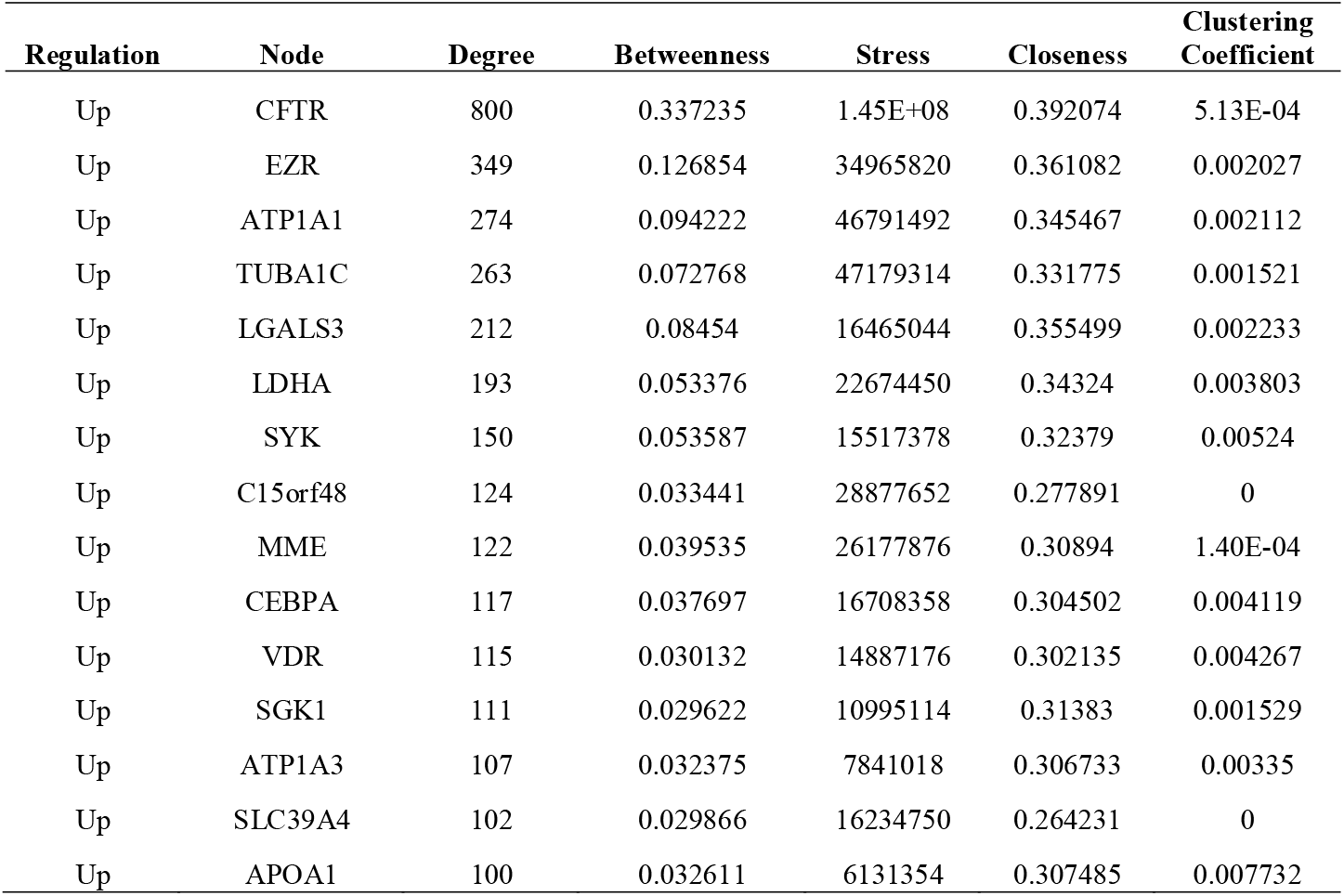
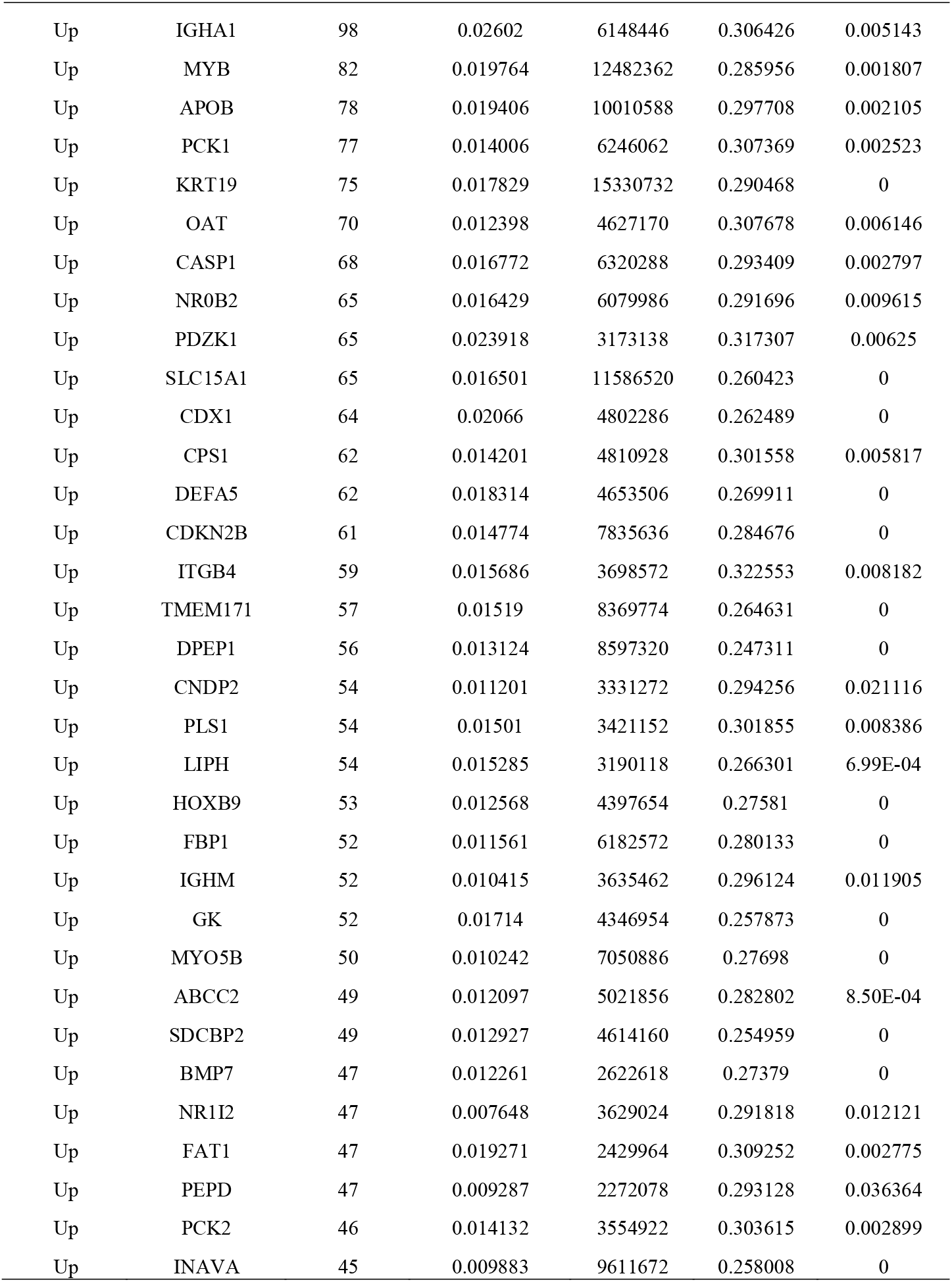
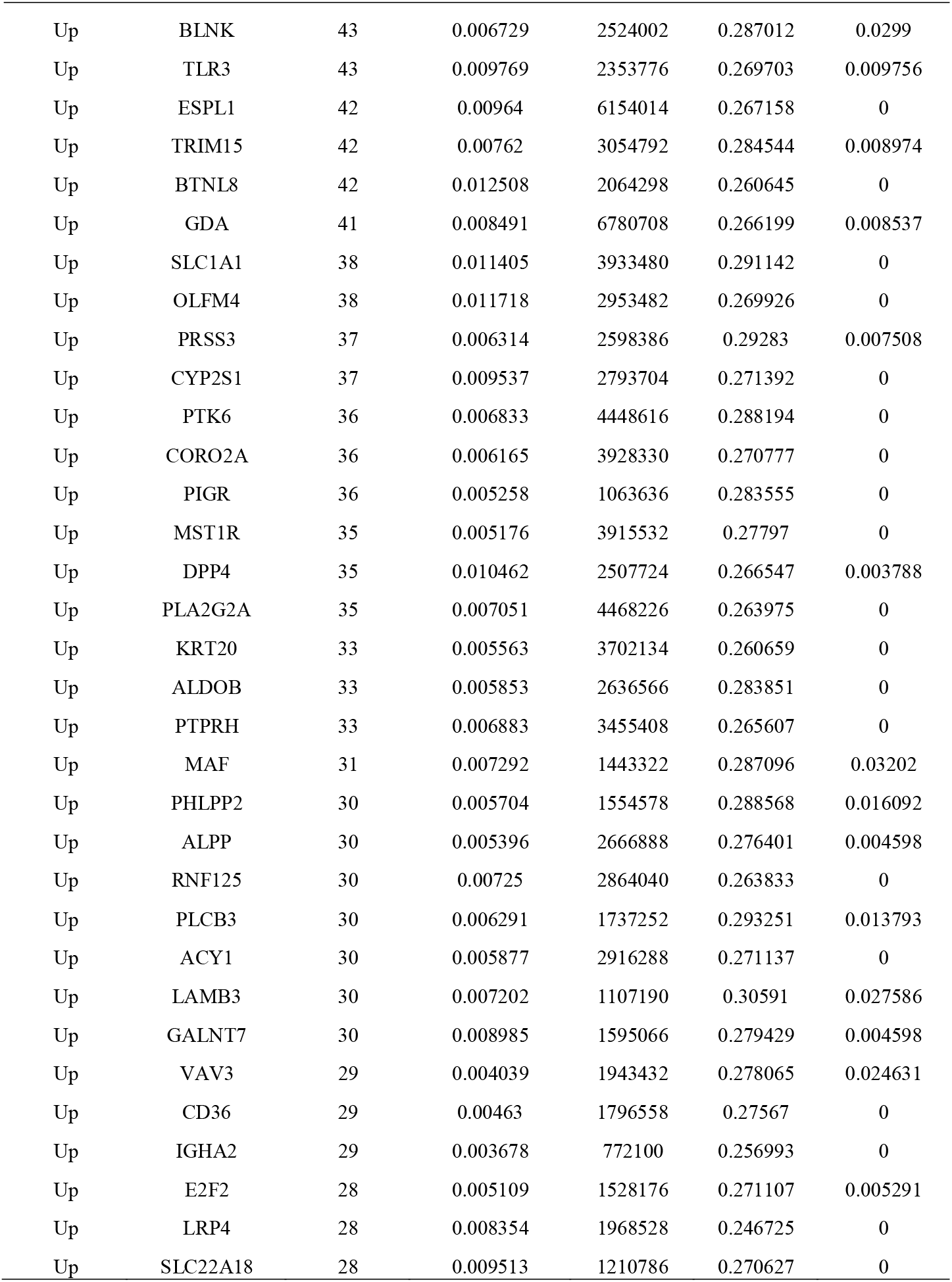
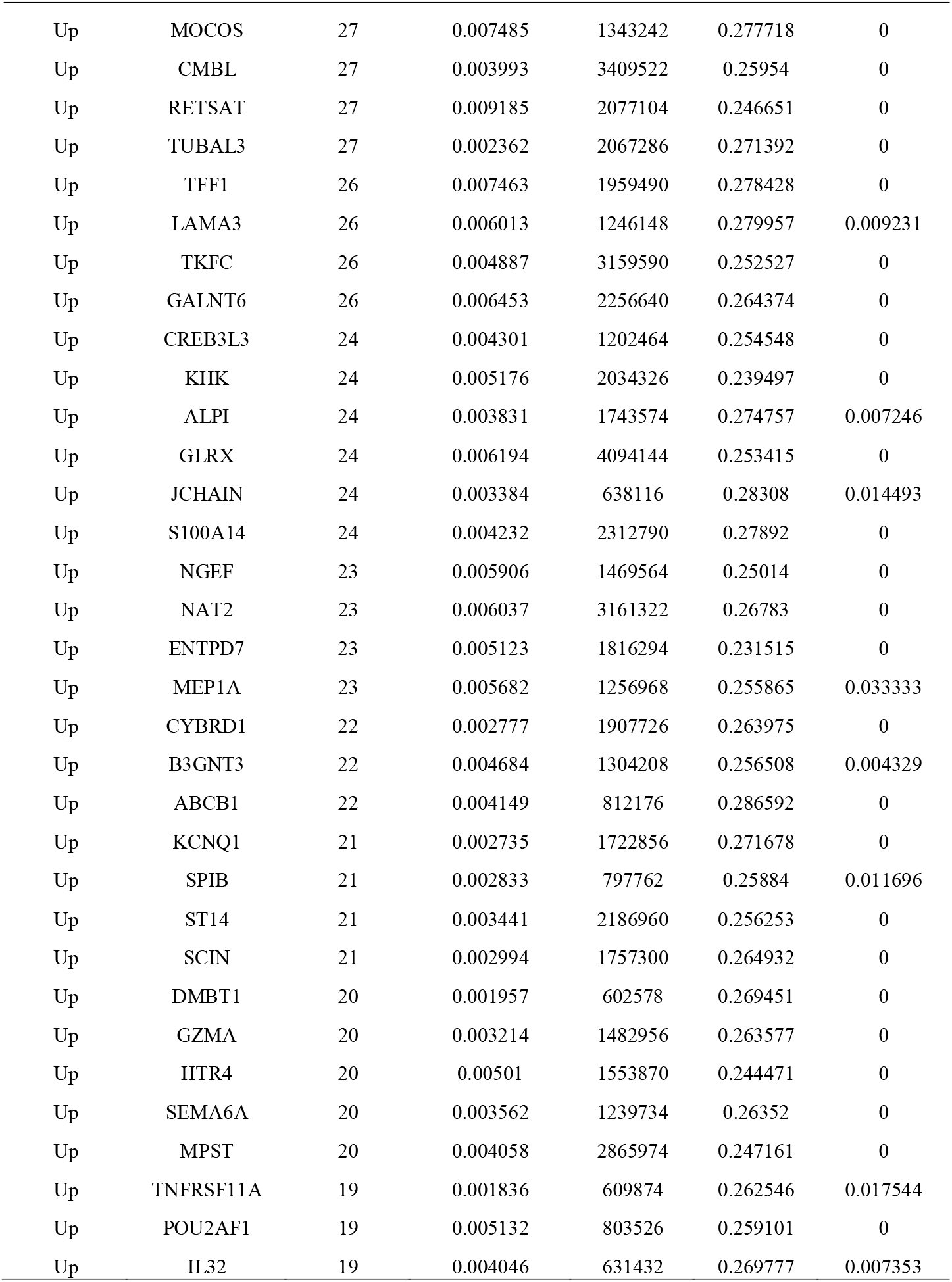
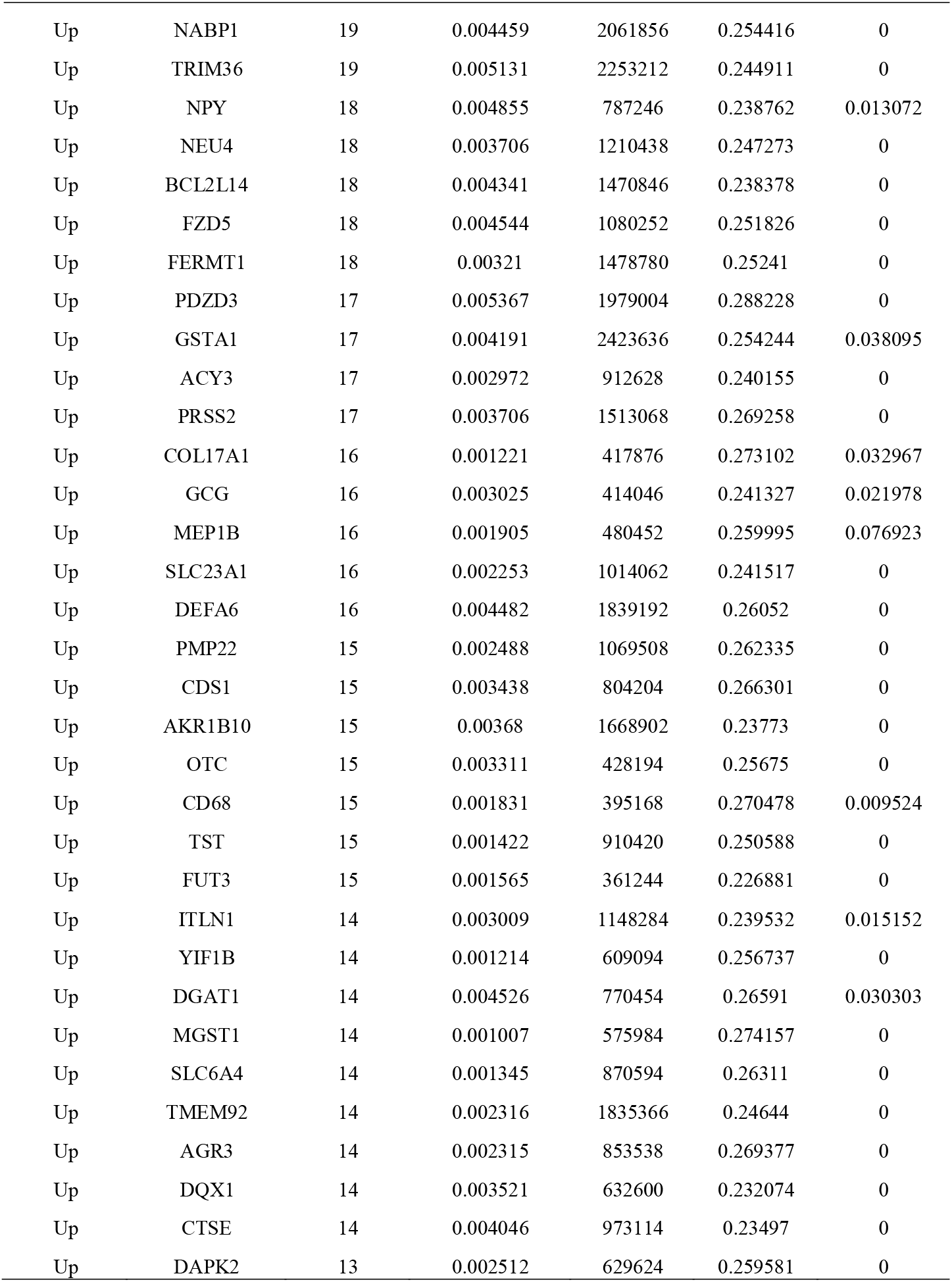
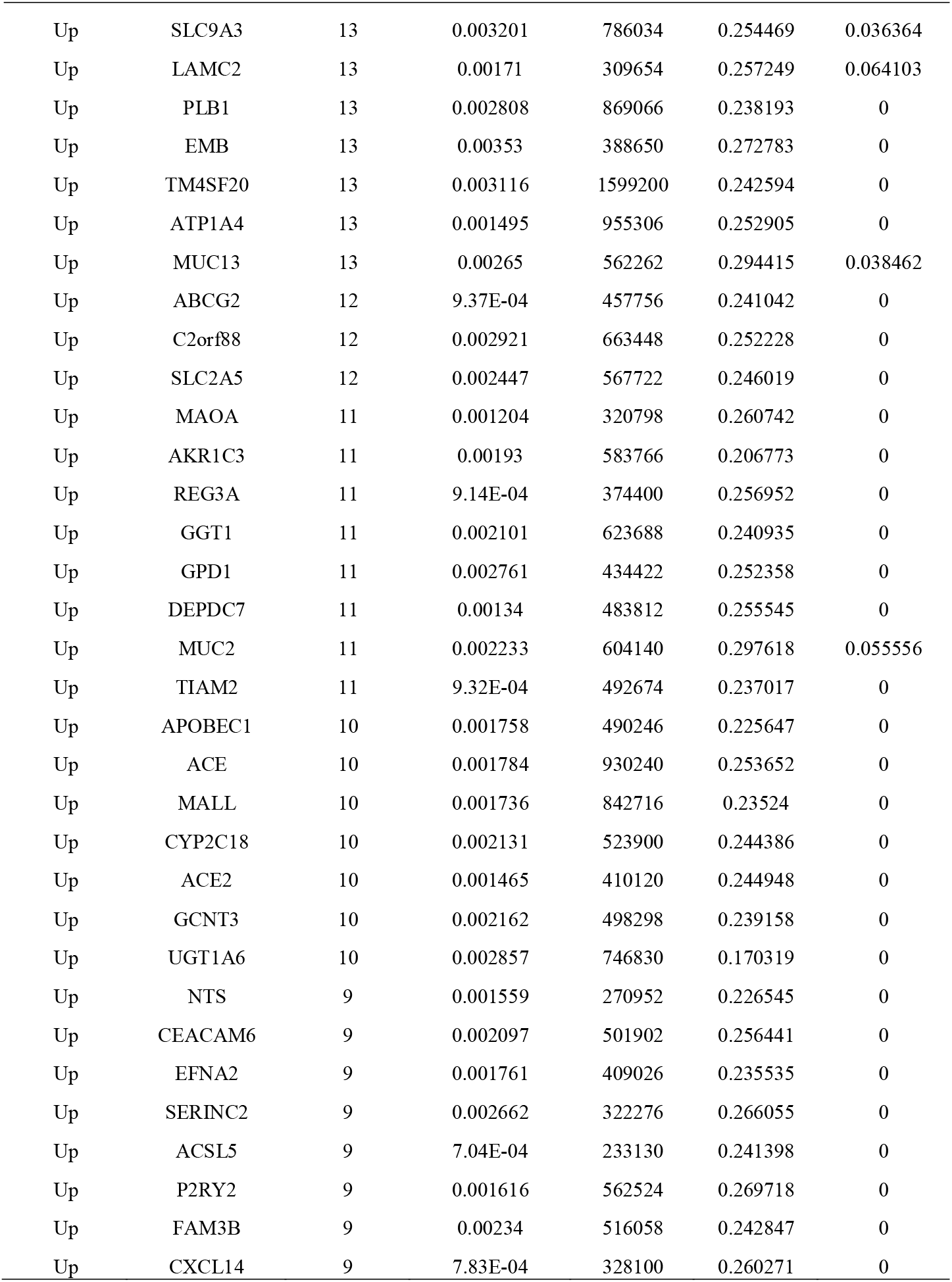
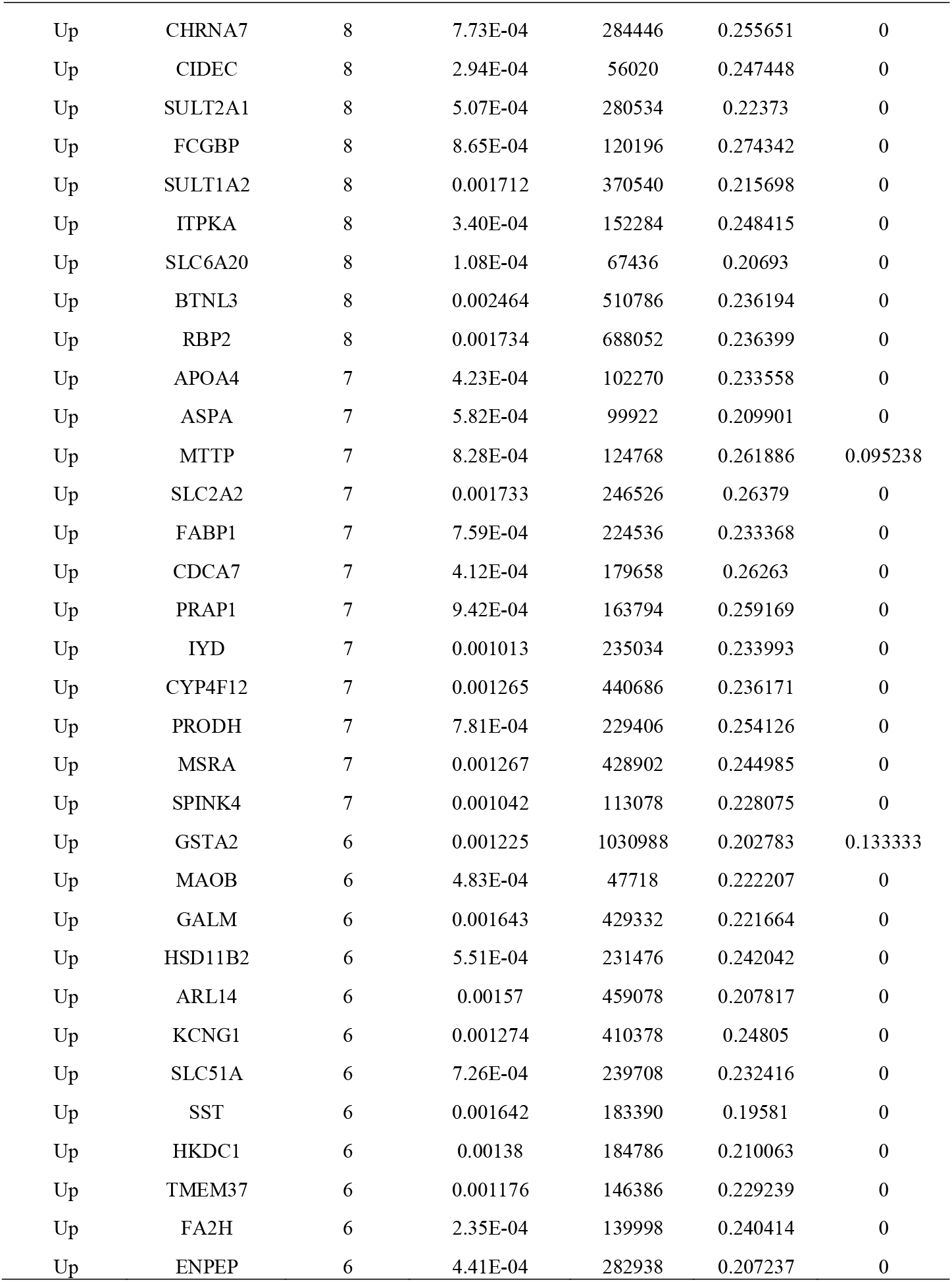
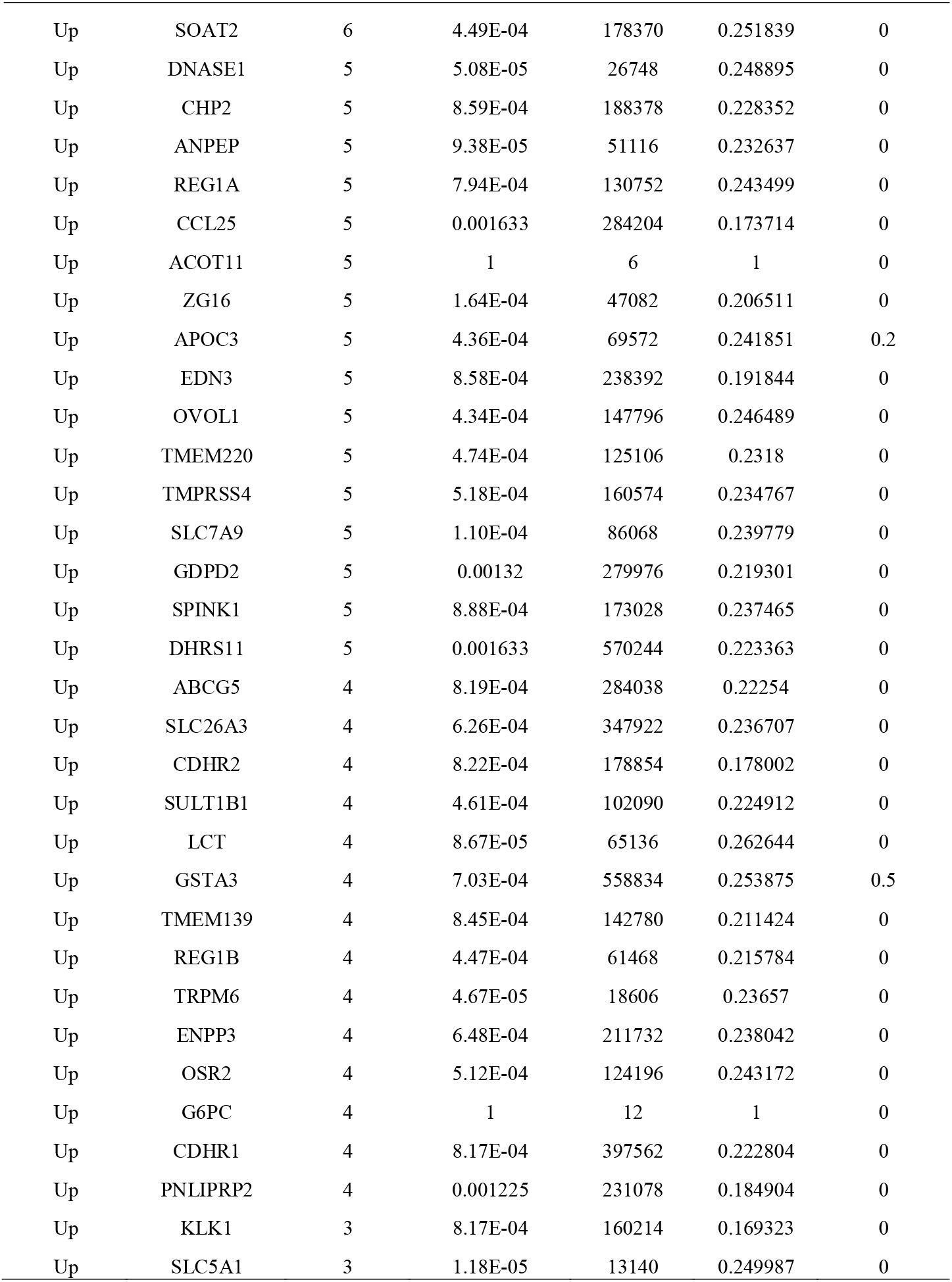
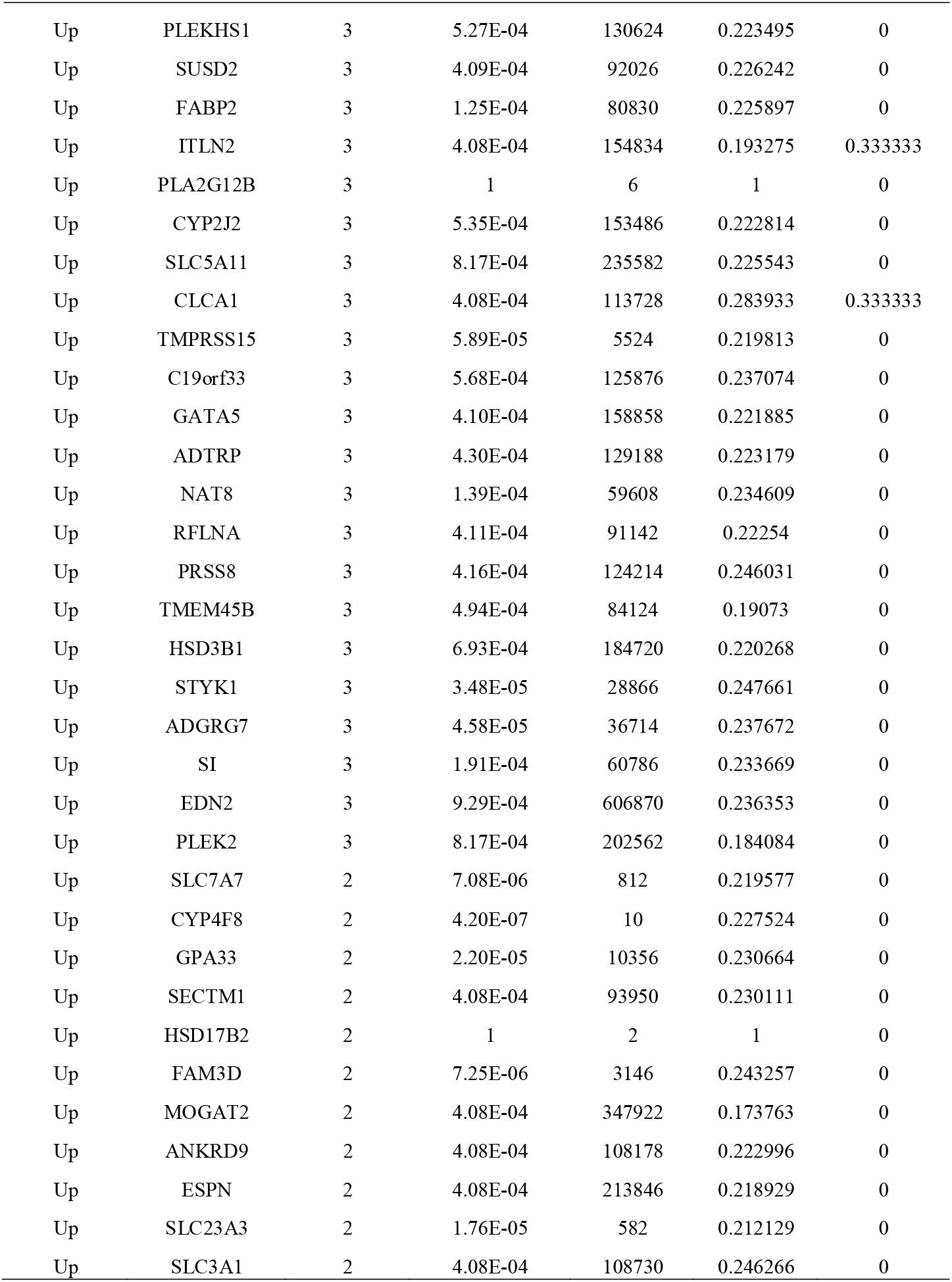
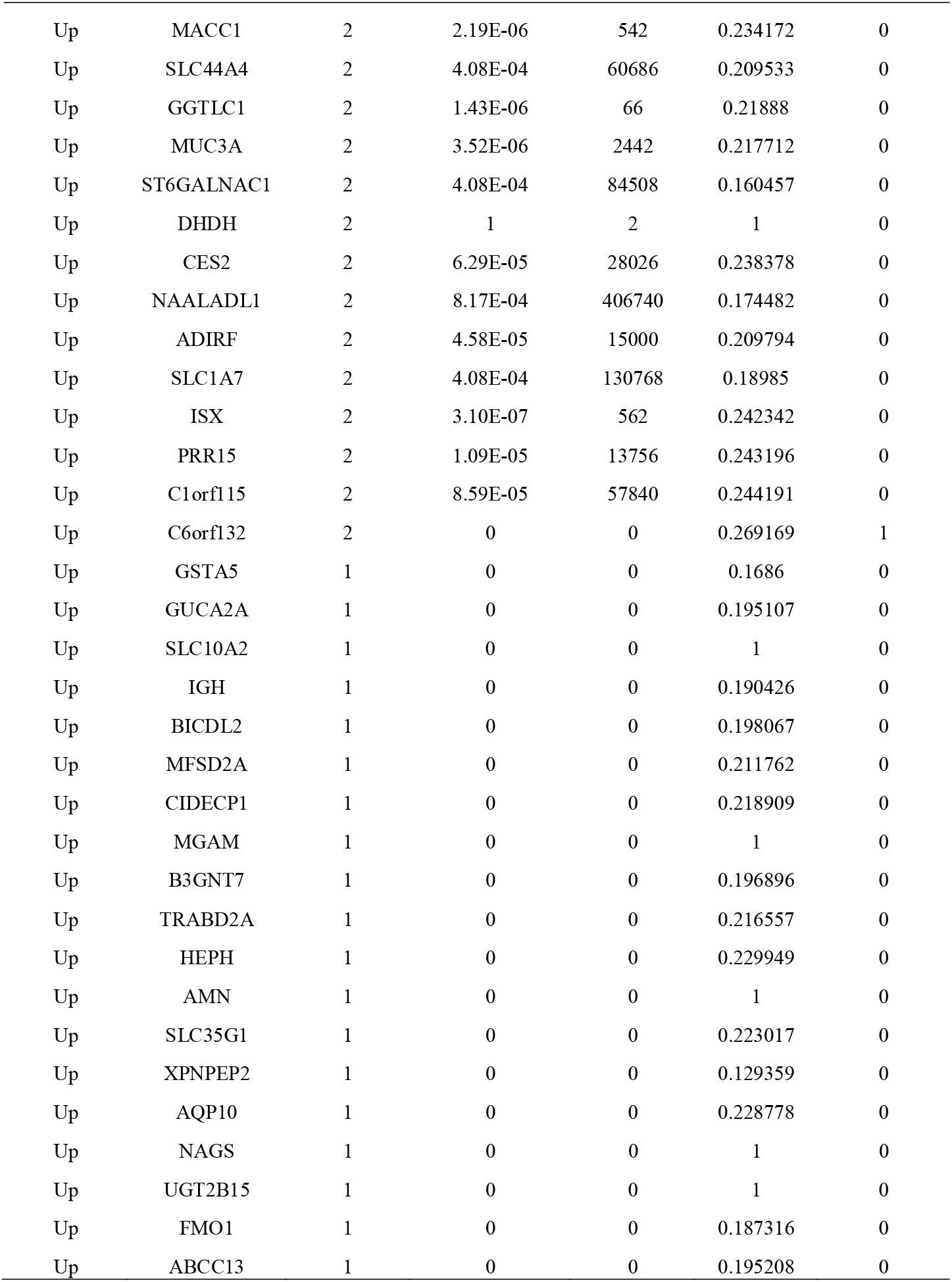
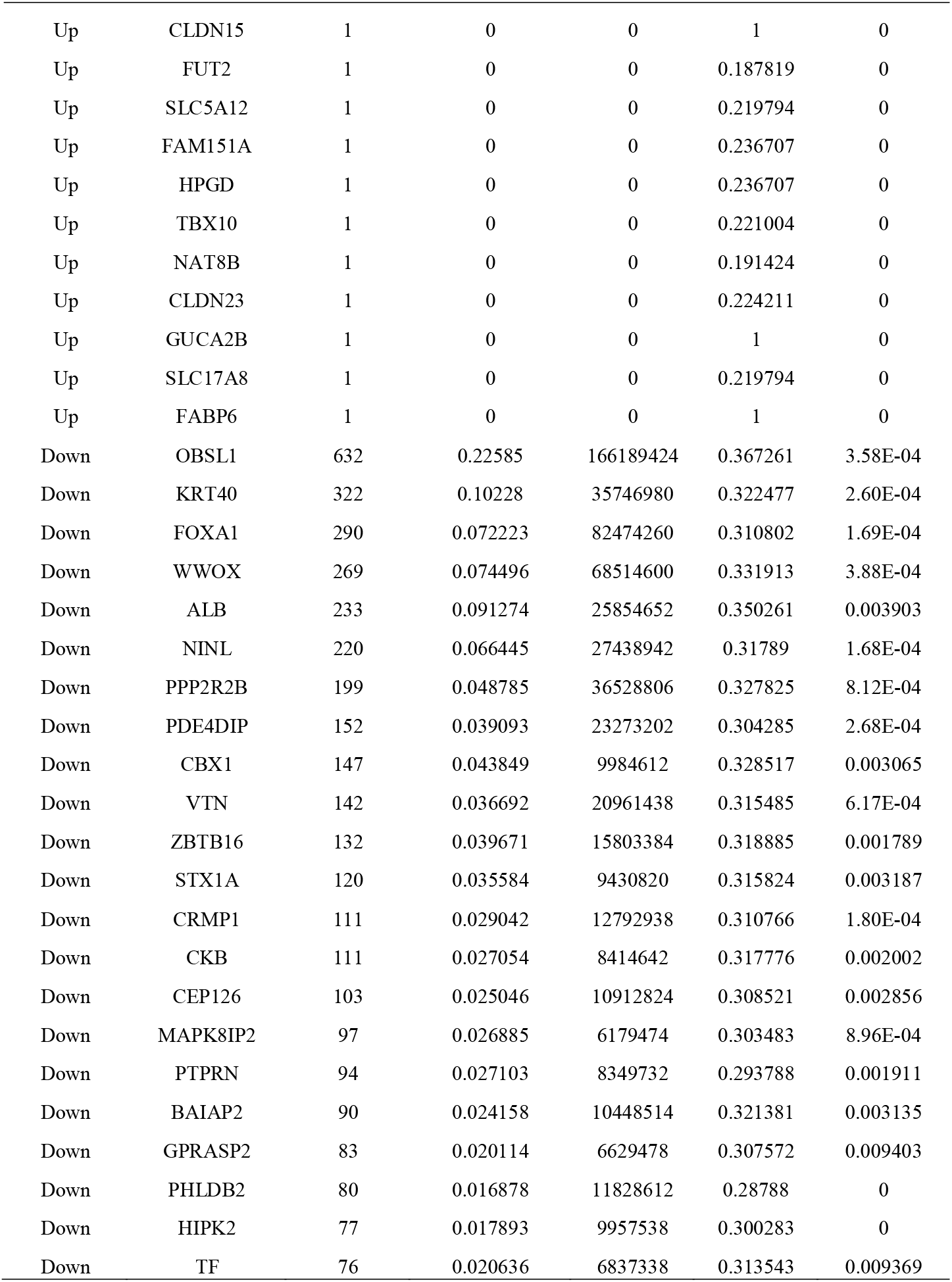
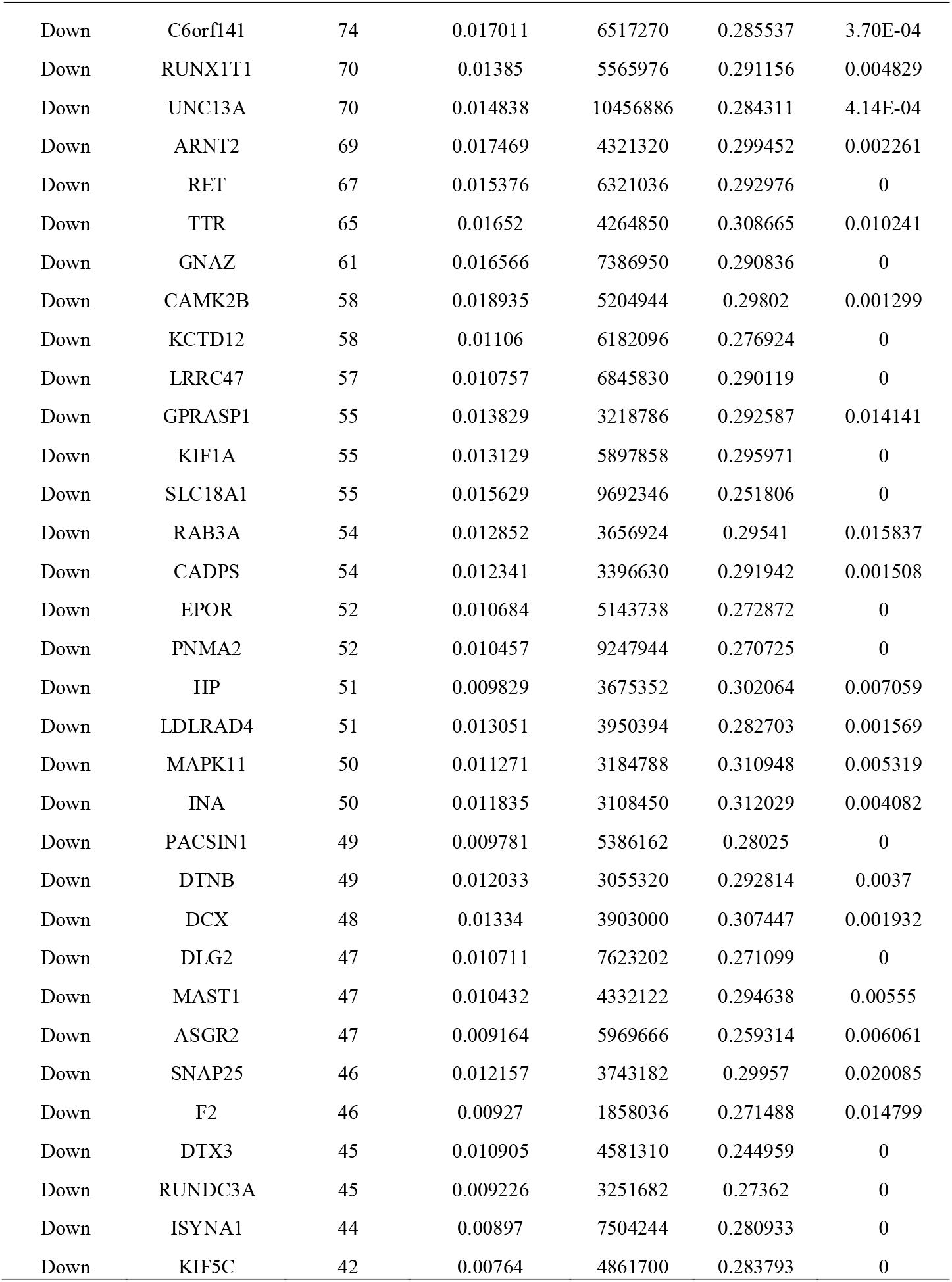
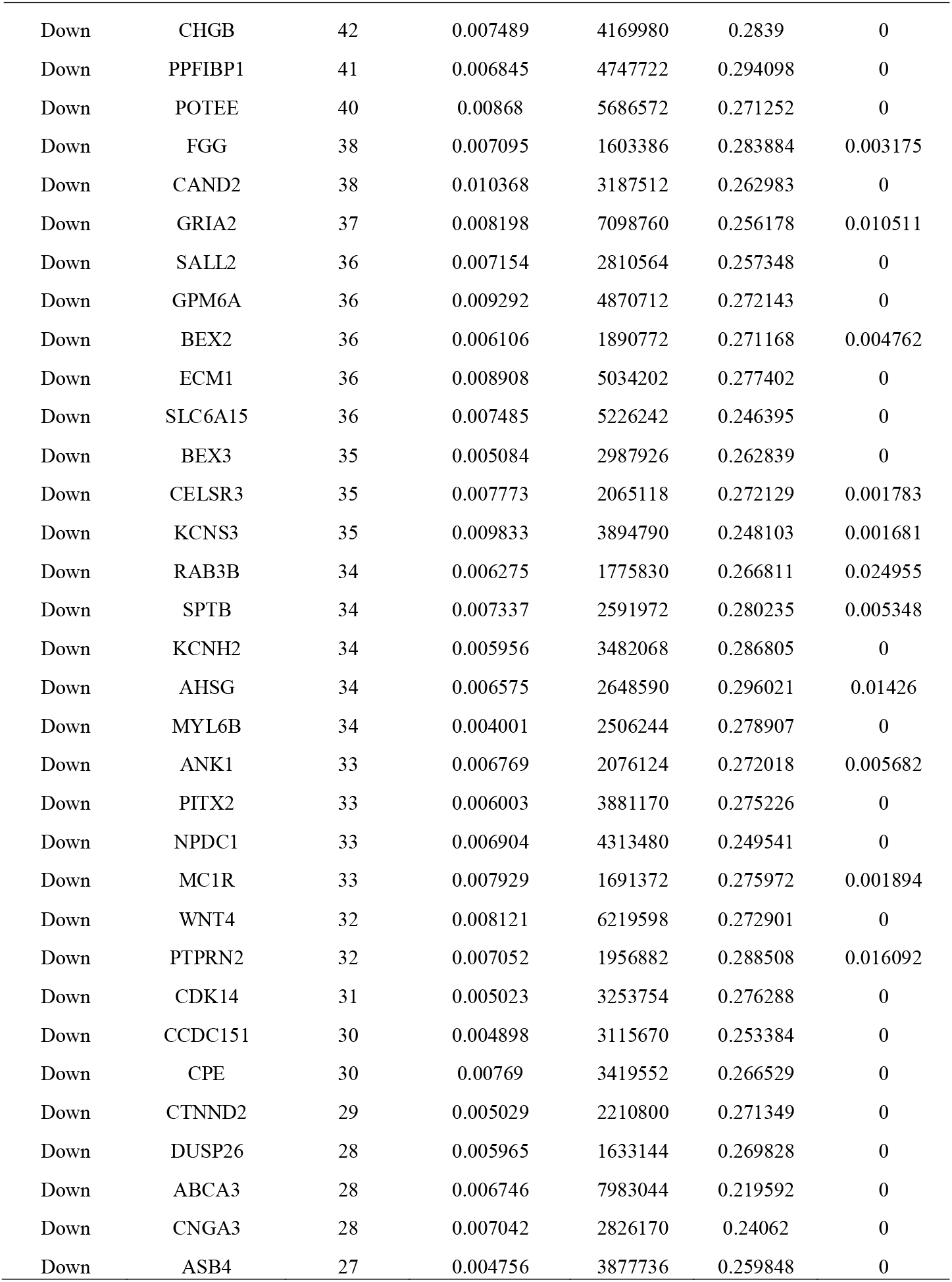
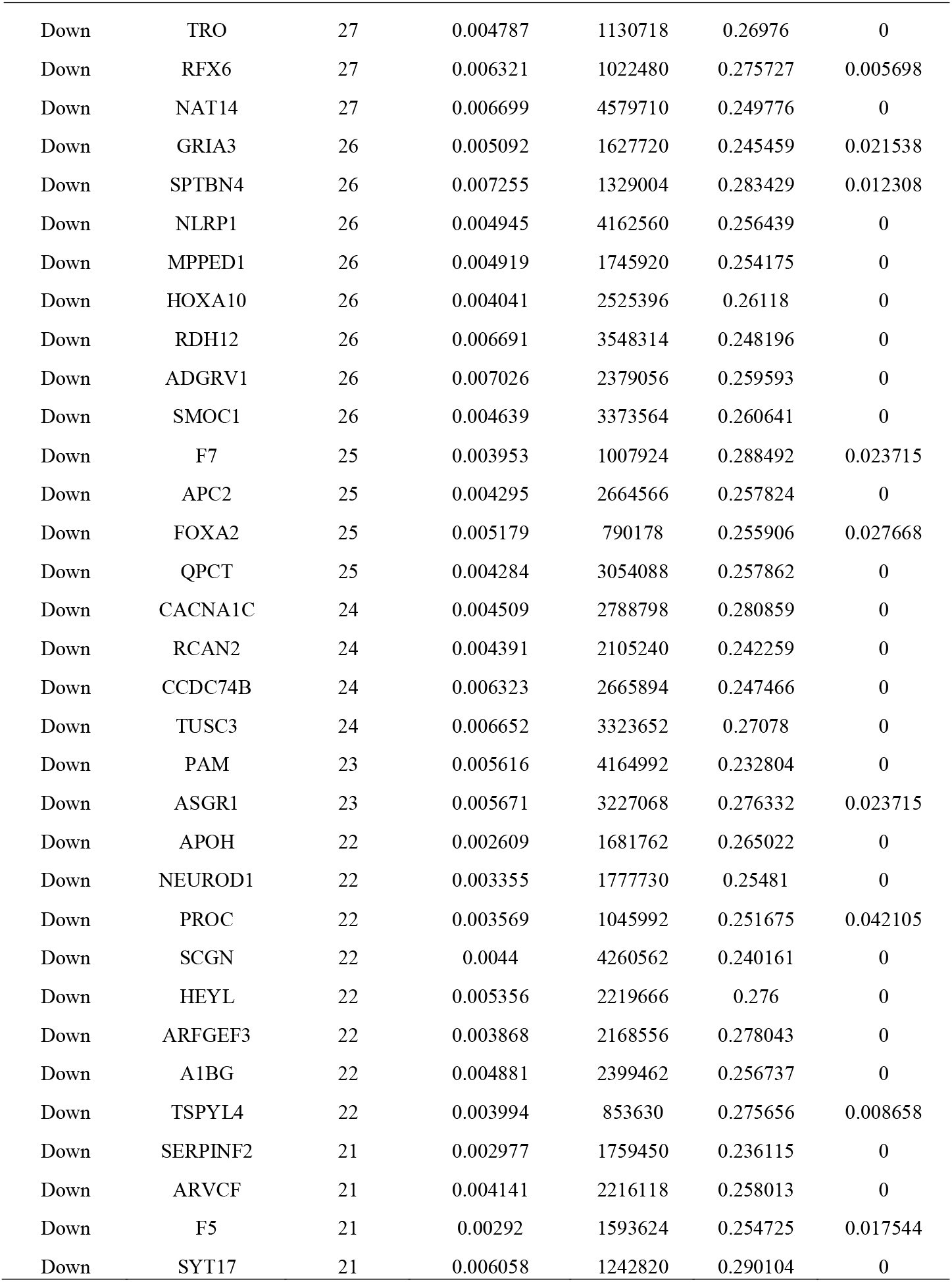
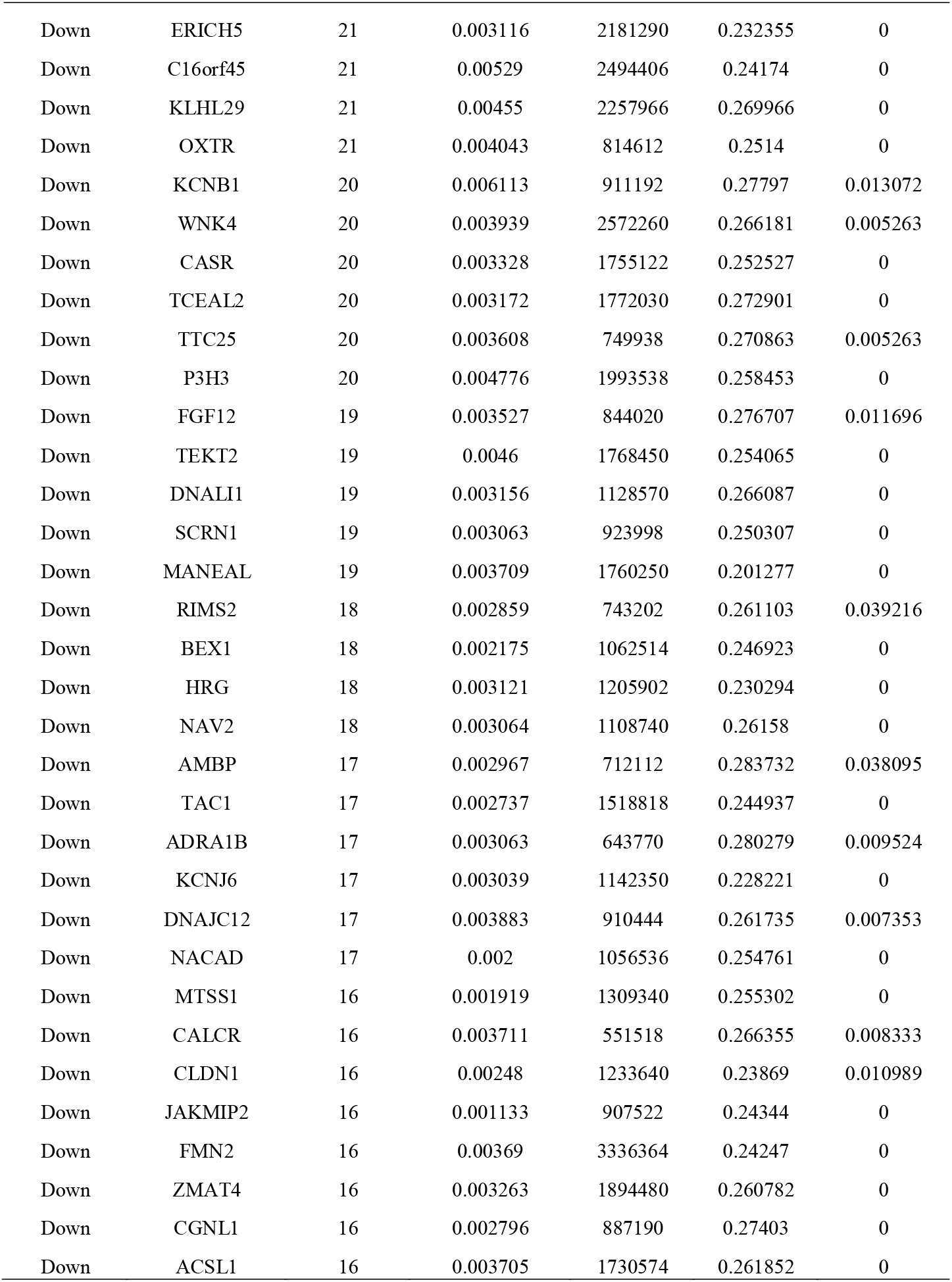
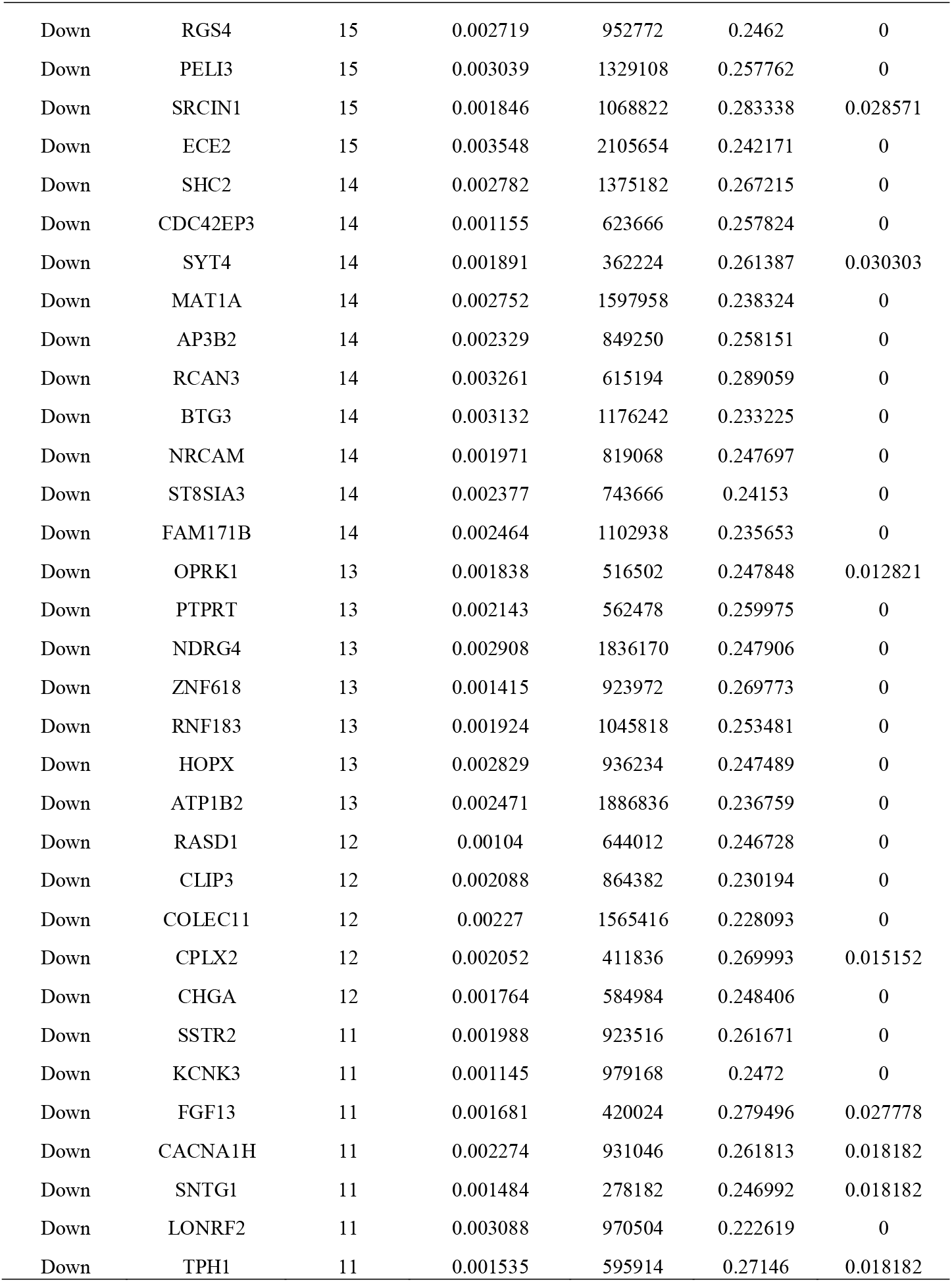
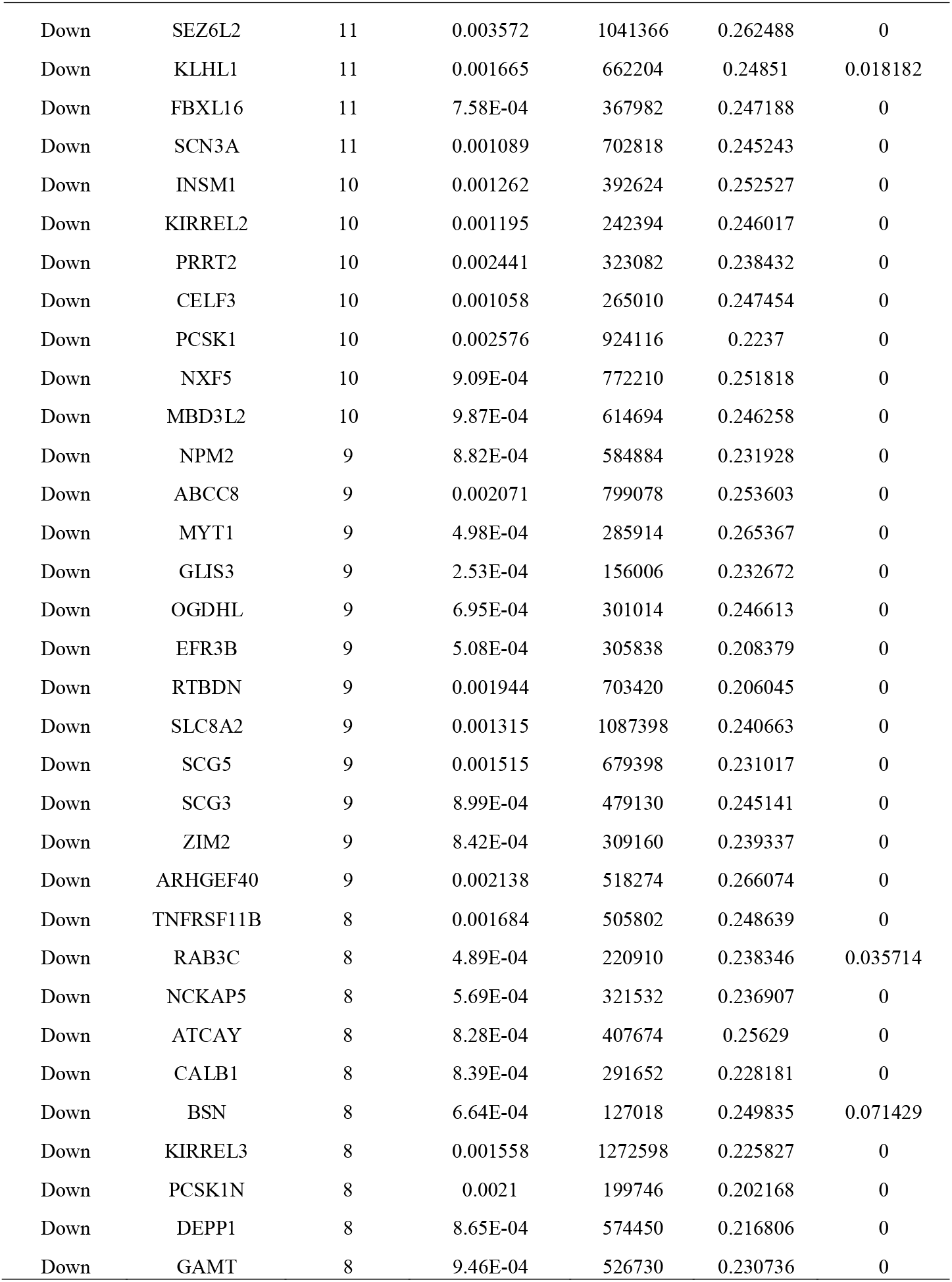
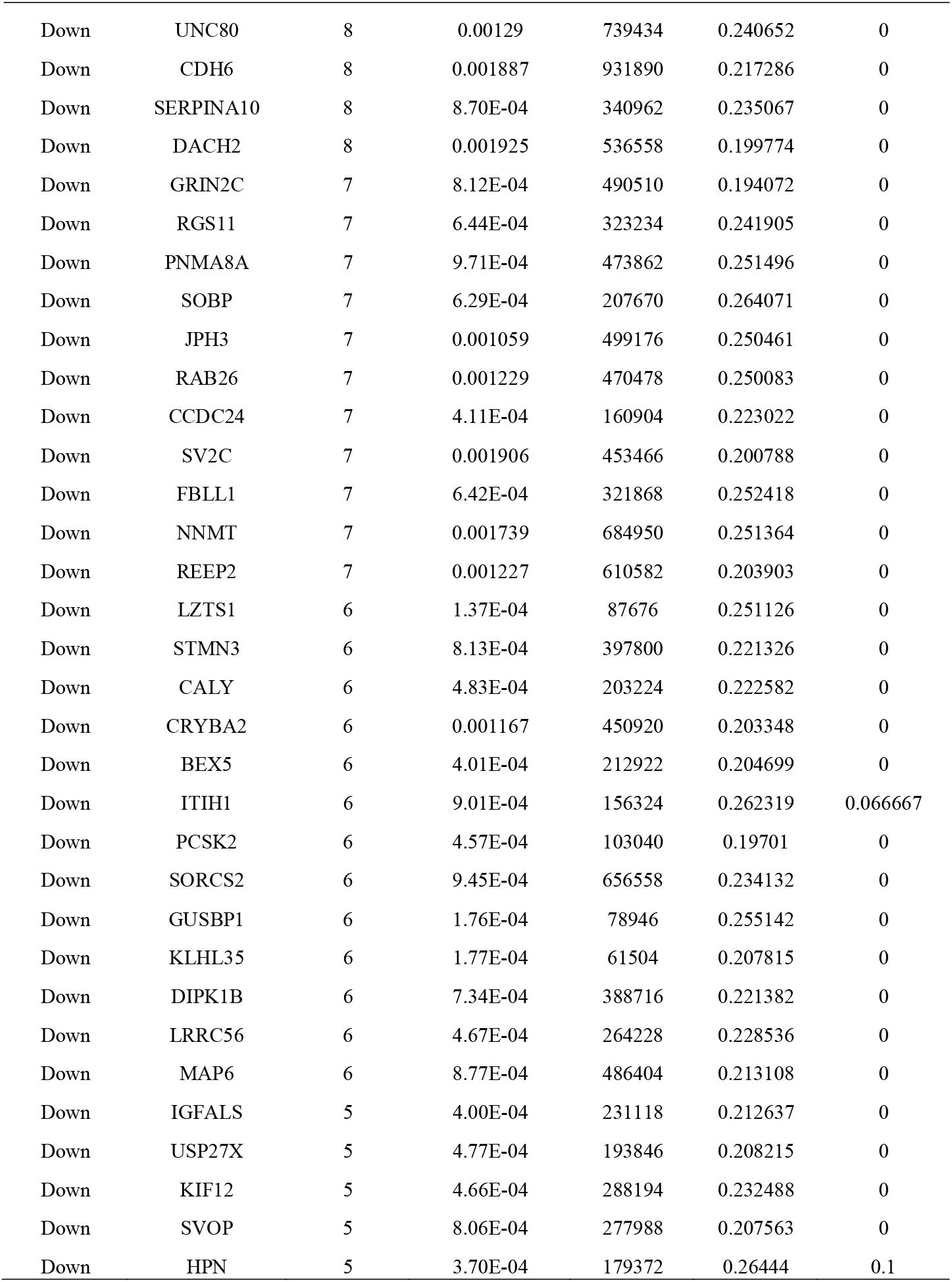
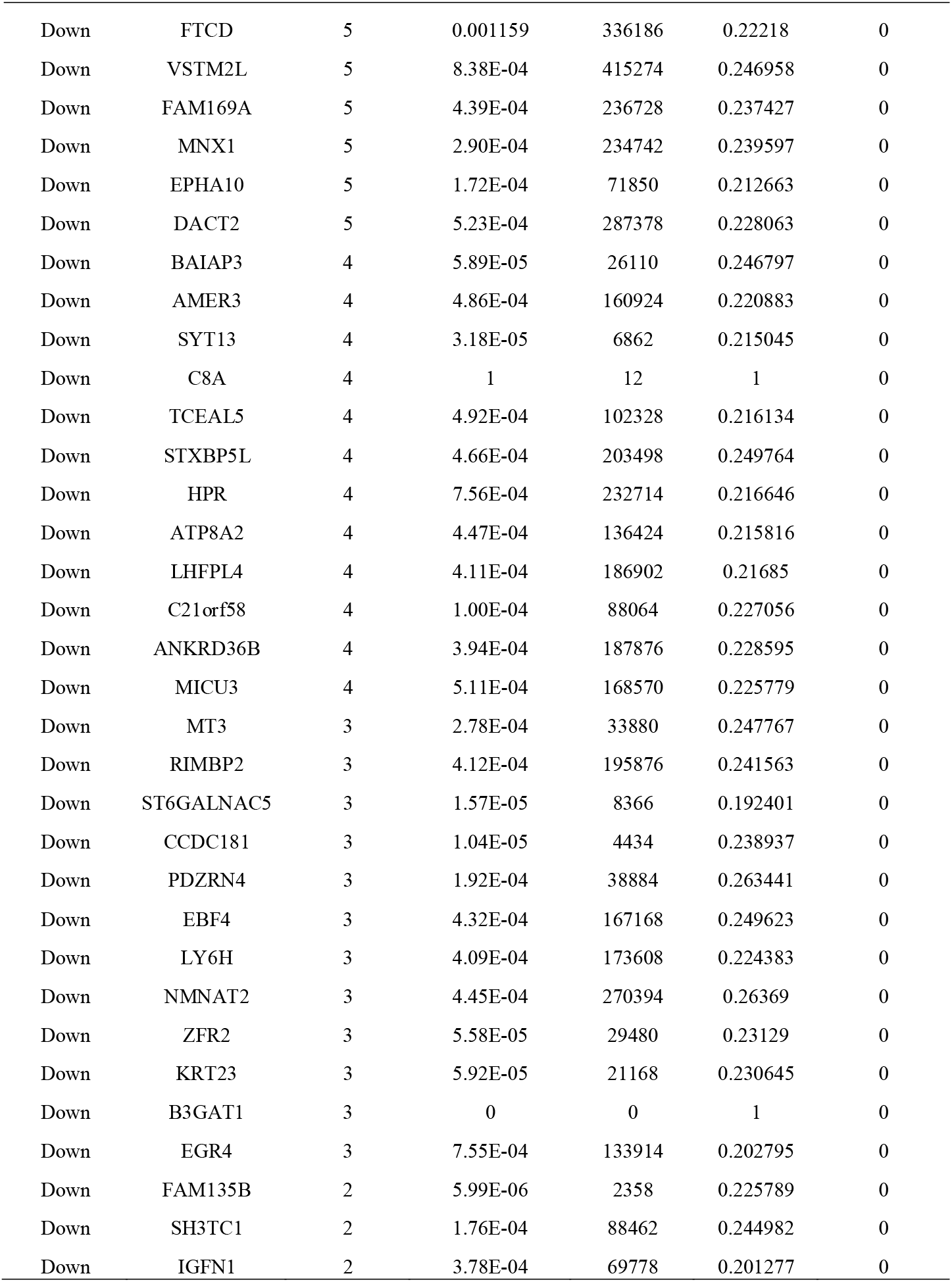
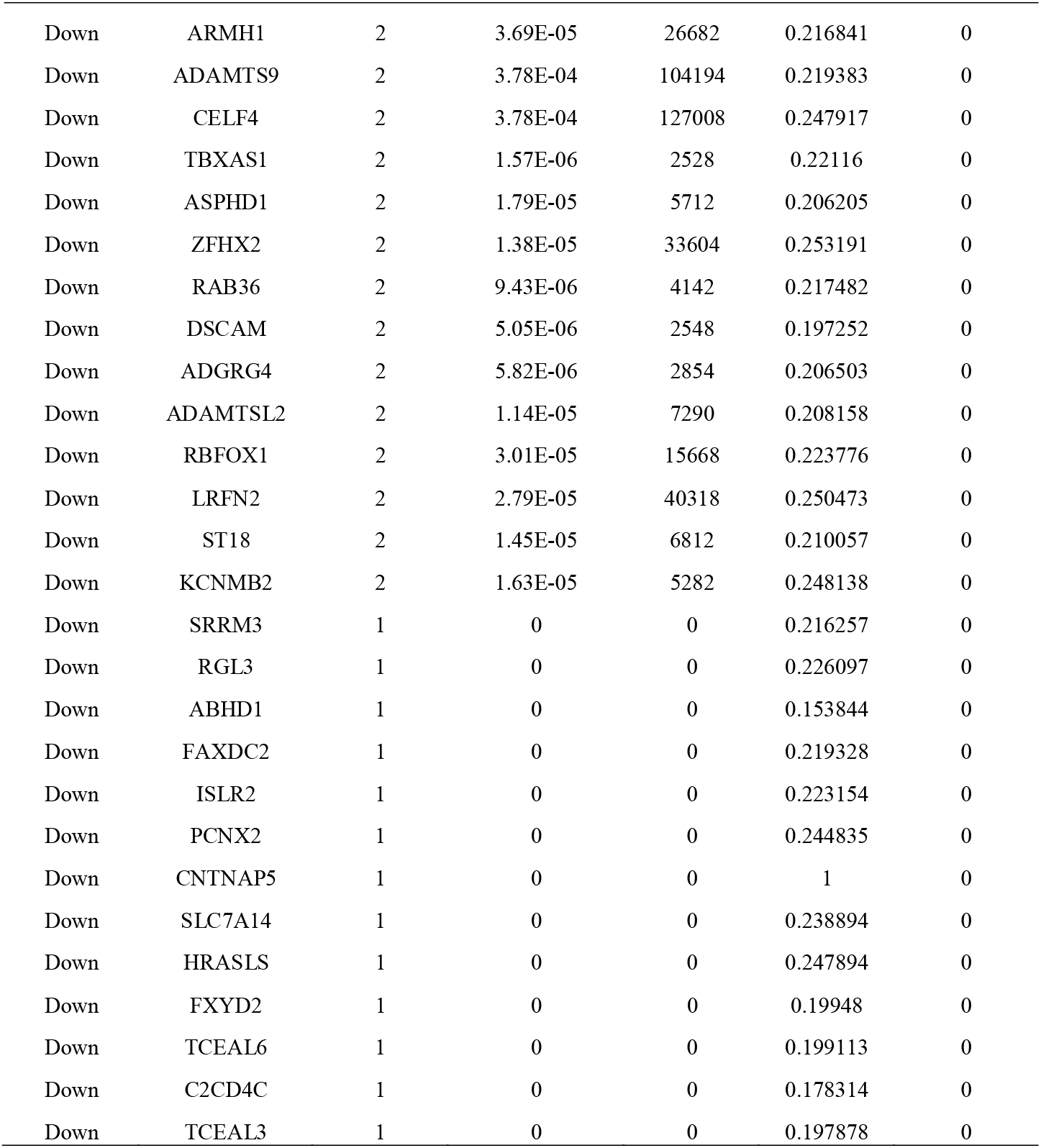
Topology table for up and down regulated genes

A total of 748 modules were obtained from PPI network of up regulated genes. The four most significant modules (module 1, module 3, module 5 and module 6) were obtained using PEWCC1 in Cytoscape (Fig. 5A). Module 1 had 59 nodes and 119 edges, module 3 had 24 nodes and 55 edges, module 5 had 23 nodes and 50 edges and module 6 had 21 nodes and 43 edges. Hub genes in these modules are enriched in pancreatic secretion, cystic fibrosis transmembrane conductance regulator and beta 2 adrenergic receptor pathway, anion transport, metabolic pathways, gamma-glutamyl cycle, oxoacid metabolic process, PI3K-Akt signaling pathway, EPHA forward signaling, antigen binding to B cell receptors activates protein tyrosine kinases, such as the Src family, which ultimate activate MAP kinases, genes encoding proteins affiliated structurally or functionally to extracellular matrix proteins and carbohydrate digestion and absorption. Meanwhile, total of 889 modules were extracted from PPI network of down regulated genes. The four most significant modules (module 1, module 2, module 4 and module 6) were obtained using PEWCC1 in Cytoscape (Fig. 5B). Module 1 had 23 nodes and 49 edges, module 2 had 22 nodes and 51 edges, module 4 had 18 nodes and 36 edges and module 6 had 14 nodes and 27 edges. Hub genes in these modules are enriched in cell projection morphogenesis, neuroactive ligand-receptor interaction, thyroid hormone synthesis, hormone transport, secretory granule, insulin secretion, modulation of chemical synaptic transmission and Synaptic vesicle trafficking.

**Figure 5.**
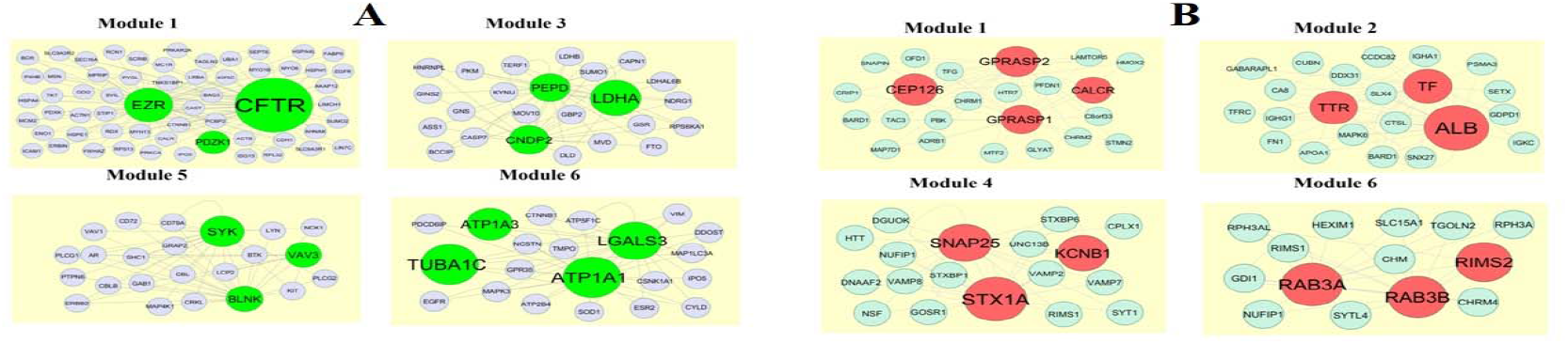
Modules in PPI network. (A) Green nodes denote the up regulated genes (B) Red nodes denote the down regulated genes

### Construction of the target gene - miRNA network

Prediction of miRNAs was aimed to further understand which miRNAs and no of miRNAs controls the up and down regulated genes (Fig. 6A and Fig. 6B). Top five up regulated targeted genes such as VAV3 interacts with 164 miRNAs, SULT1B1 interacts with 131 miRNAs, MACC1 interacts with 97 miRNAs, E2F2 interacts with 78 miRNAs and SLC28A1 interacts with 75 miRNAs are listed in Table 8. These target genes in this network were enriched in EPHA forward signaling, biological oxidations and transmembrane transport of small molecules. Meanwhile, top five down regulated targeted genes such as SVOP interacts with 107 miRNAs, KCNJ6 interacts with 90 miRNAs, SORCS2 interacts with 88 miRNAs, HEYL interacts with 83 miRNAs and SMOC1 interacts with 80 miRNAs are listed in Table 8. These target genes in this network were enriched in vesicle membrane, dopaminergic synapse, neuron differentiation and calcium ion binding.

**Figure 6.**
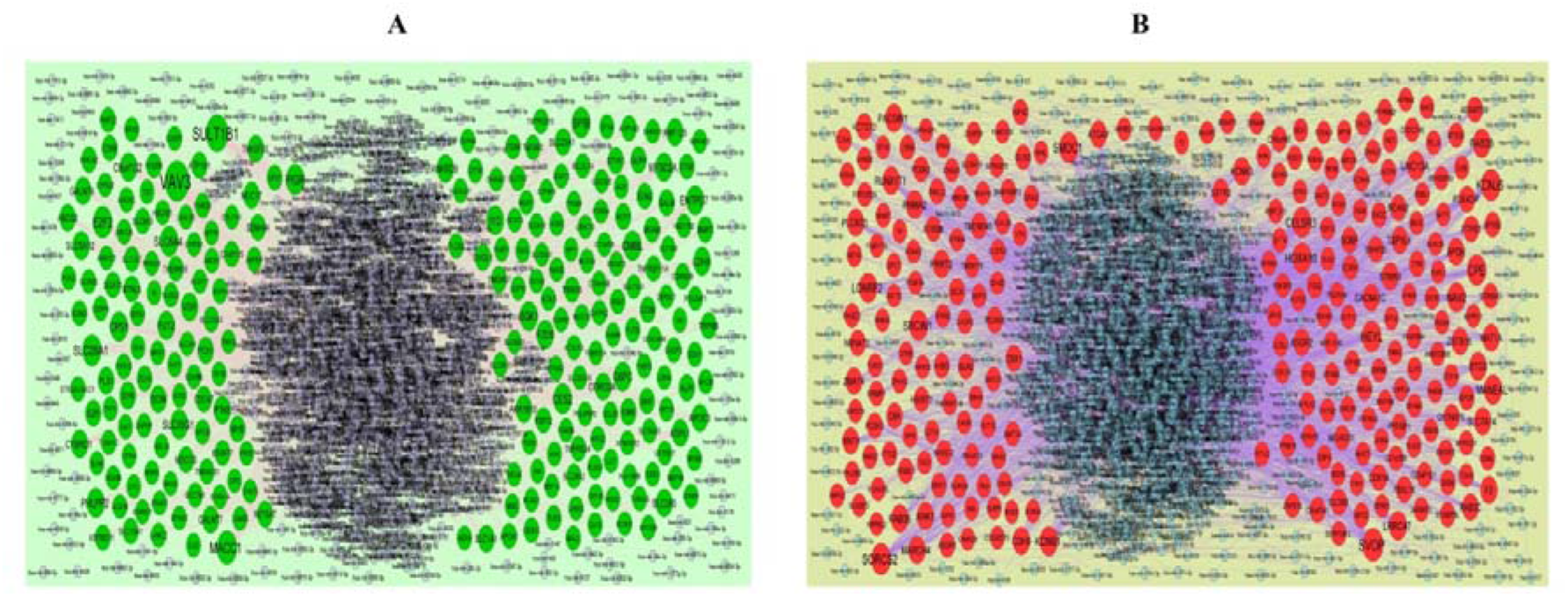
(A) The network of up regulated genes and their related miRNAs. The green circles nodes are the up regulated DEGs and purple diamond nodes are the miRNAs (B) The network of down regulated genes and their related miRNAs. The red circle nodes are the down regulated DEGs and blue diamond nodes are the miRNAs

**Table 8.**
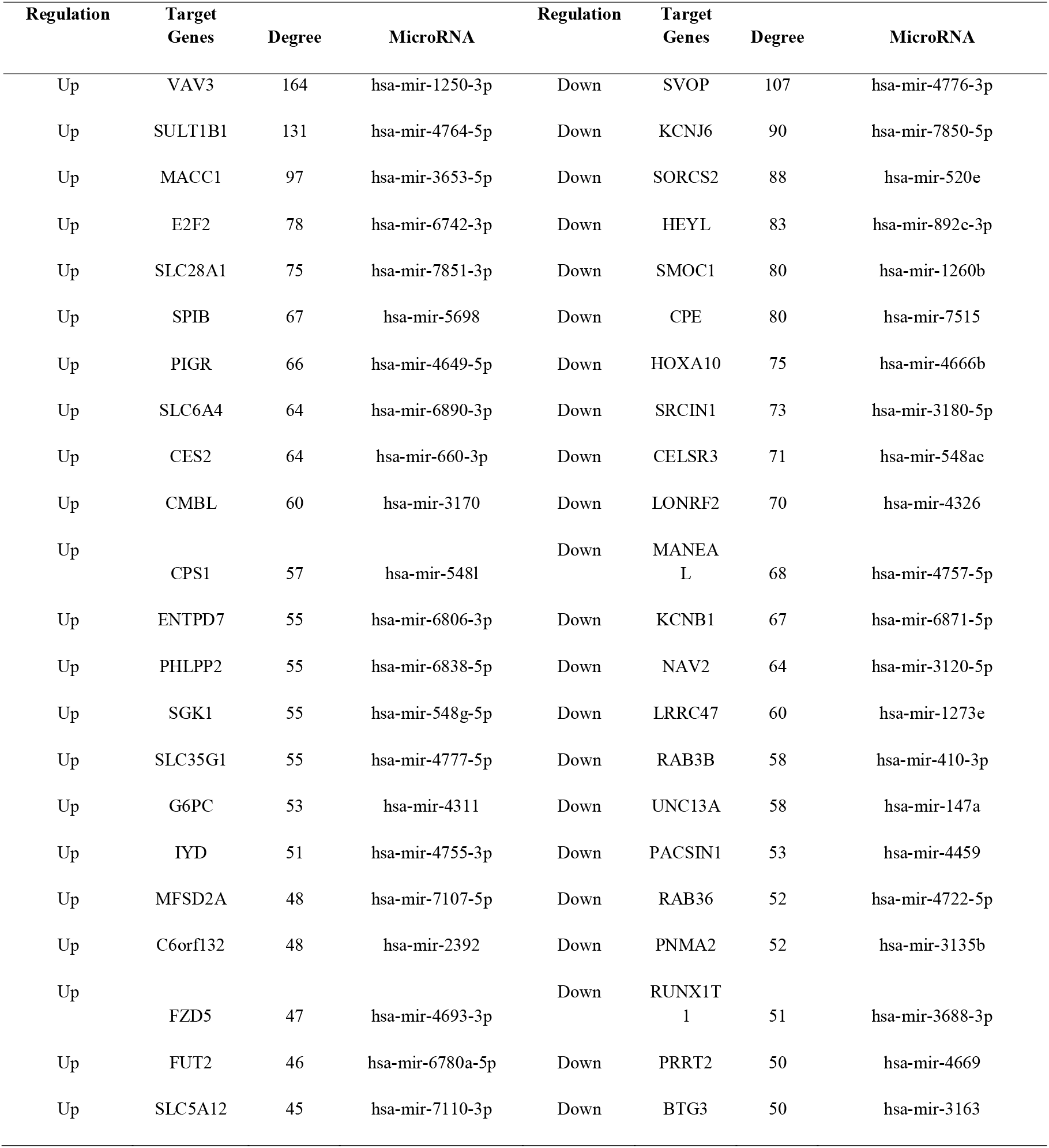
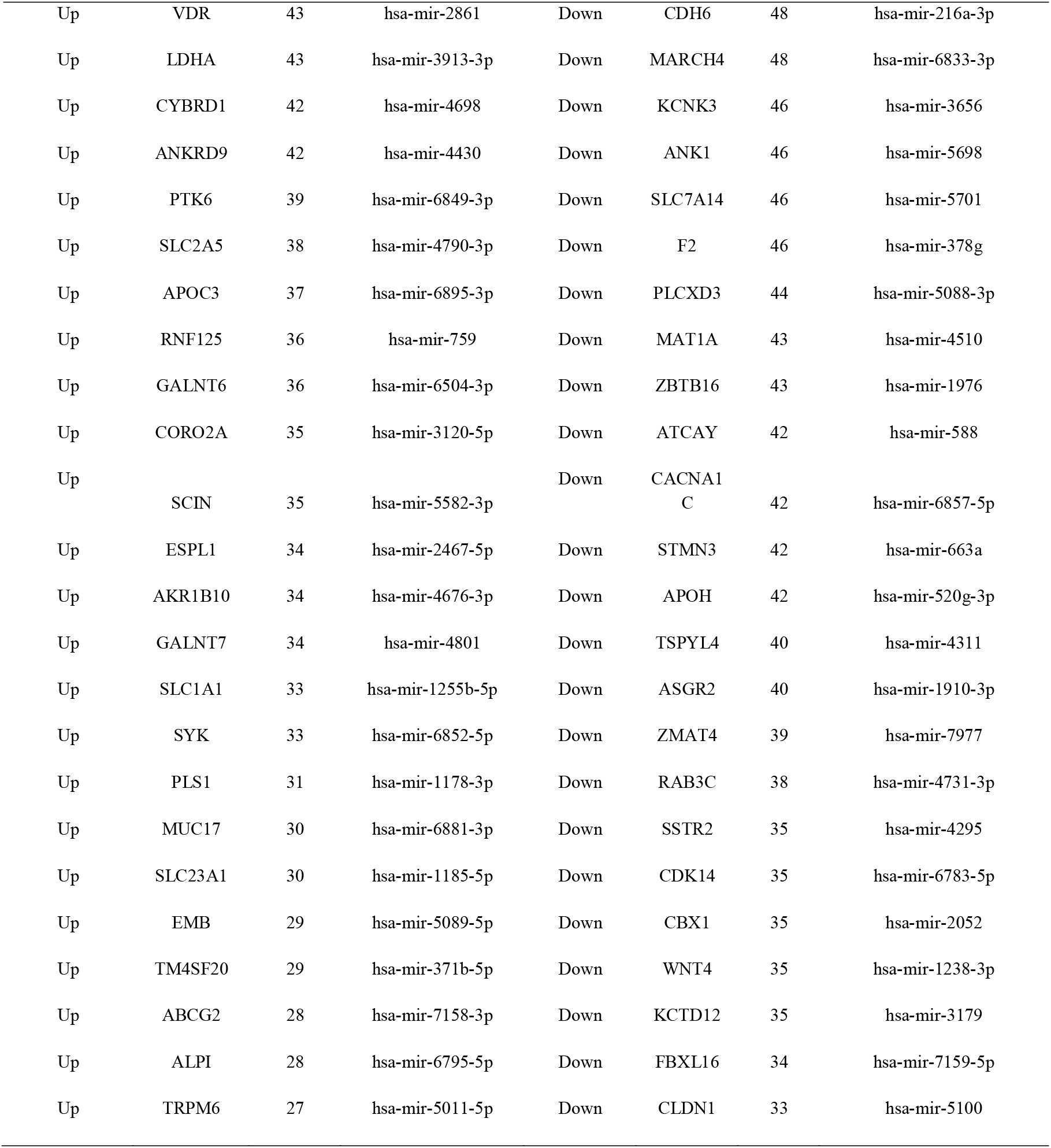
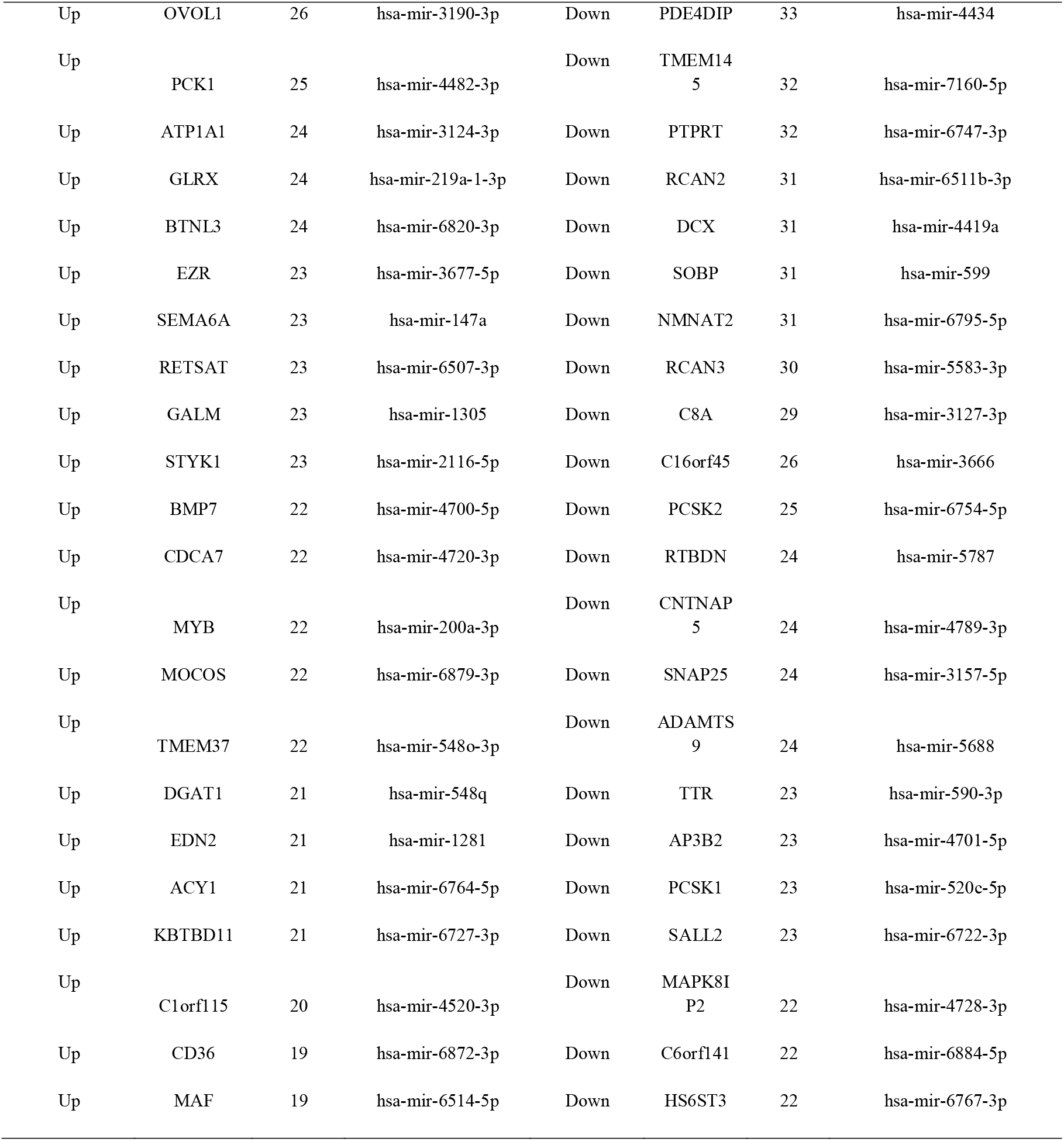
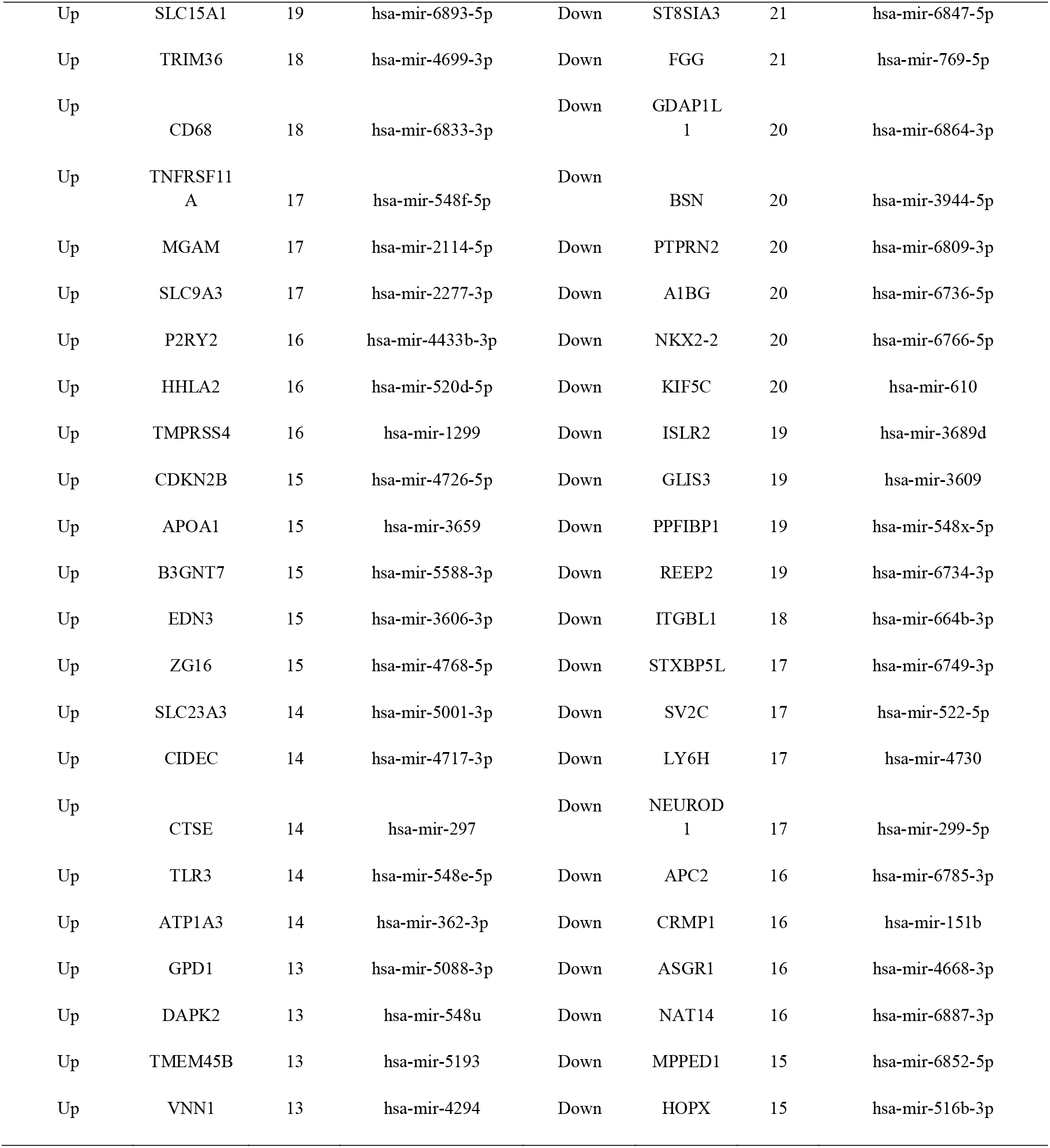
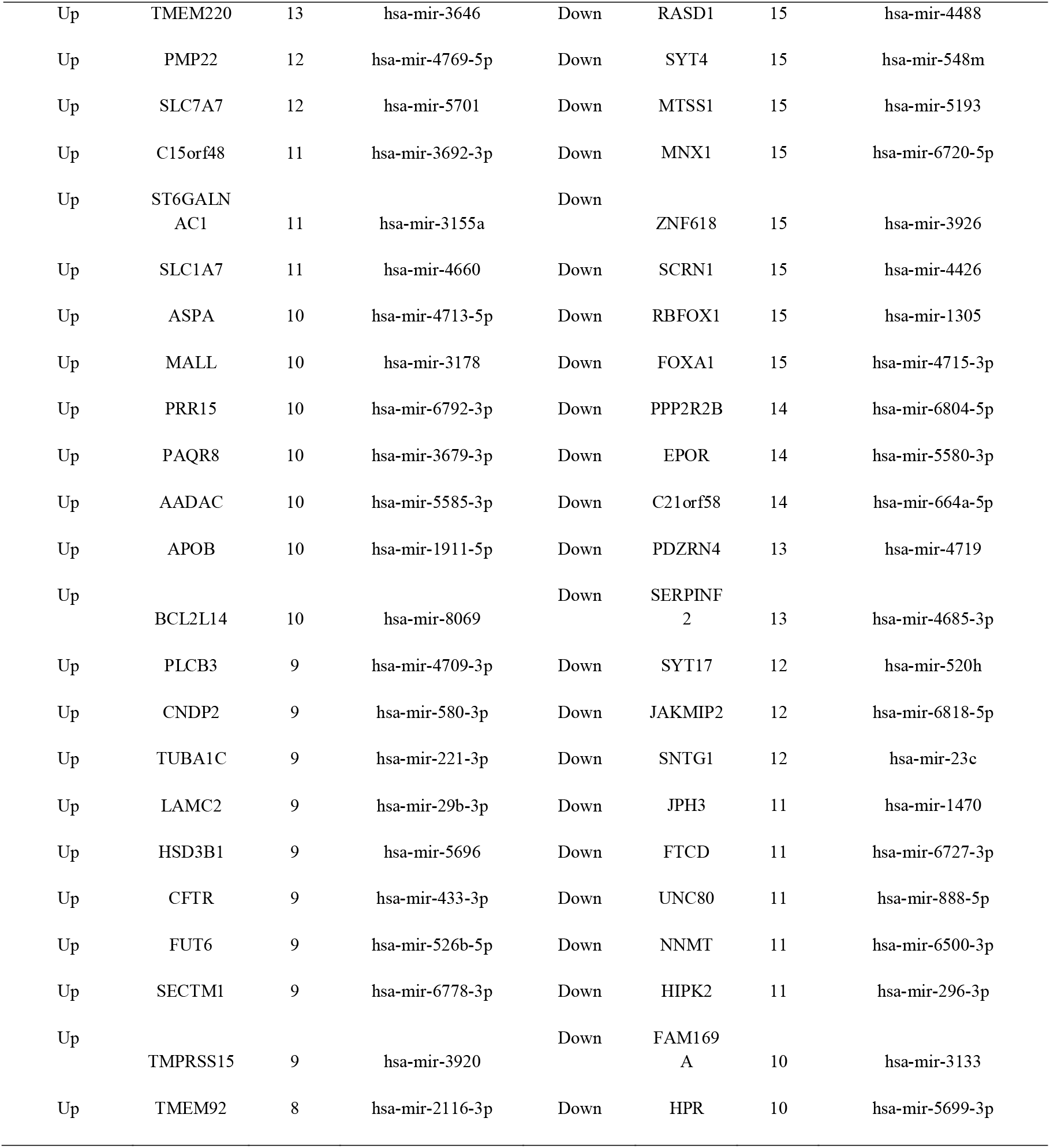
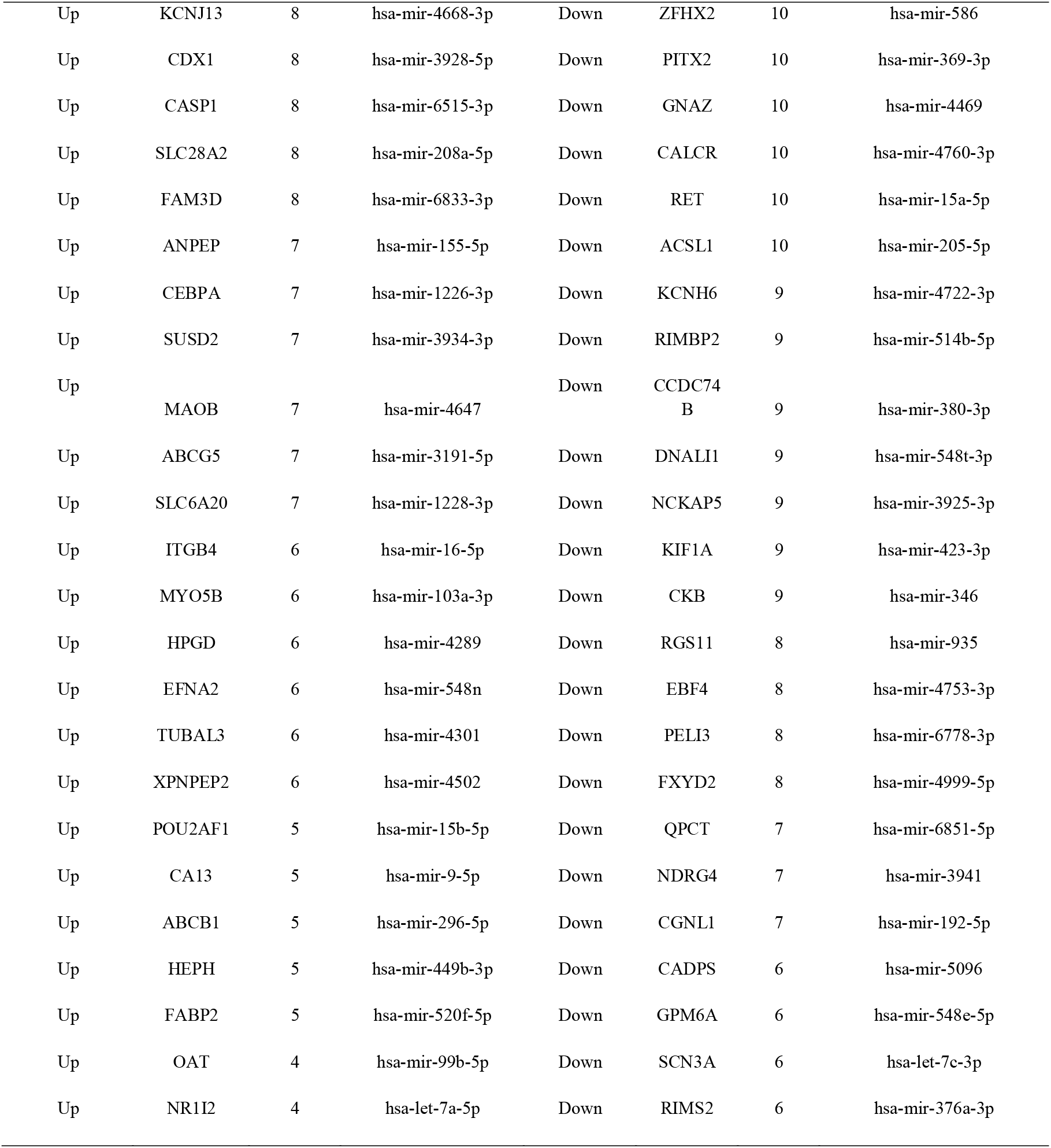
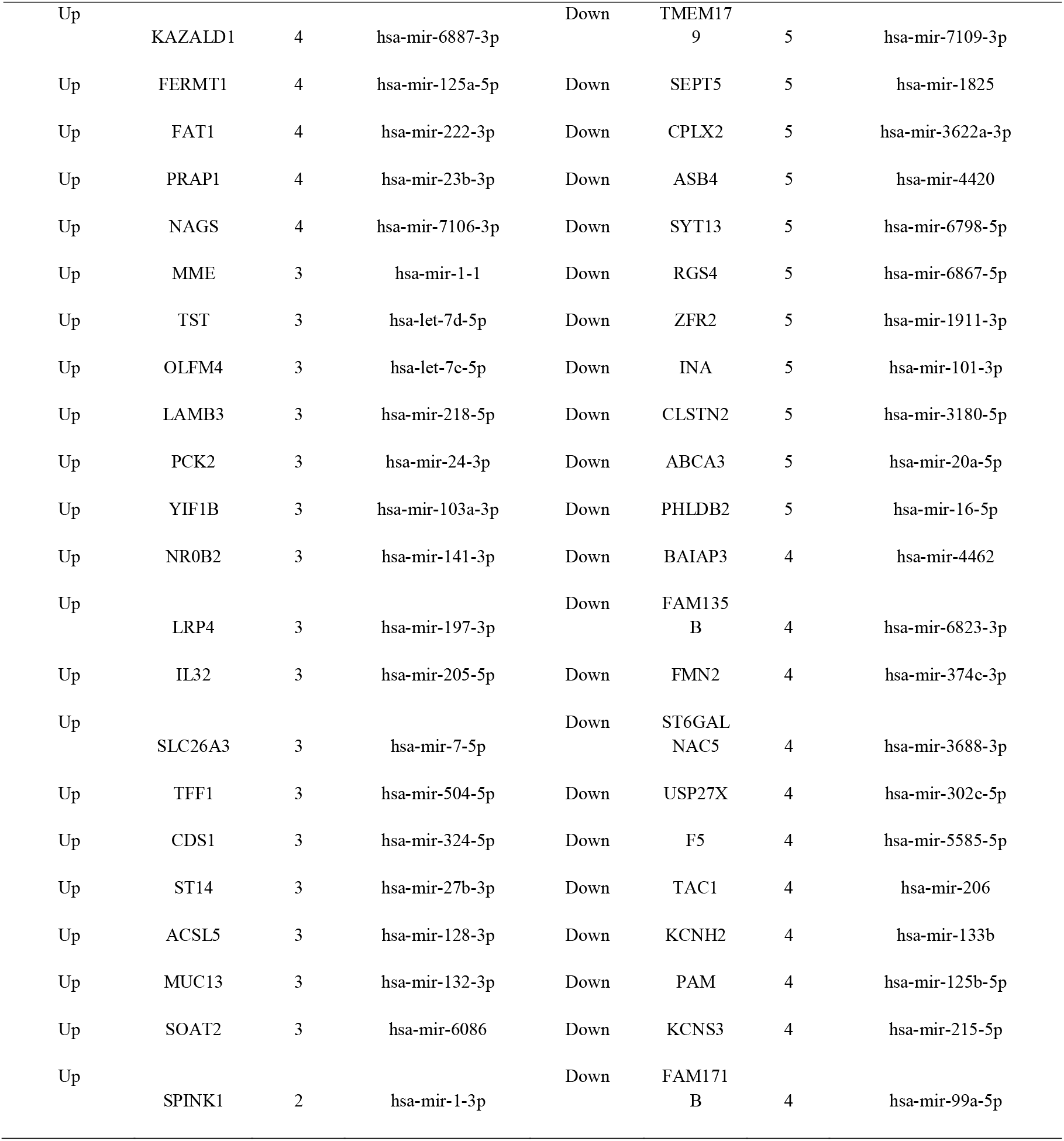
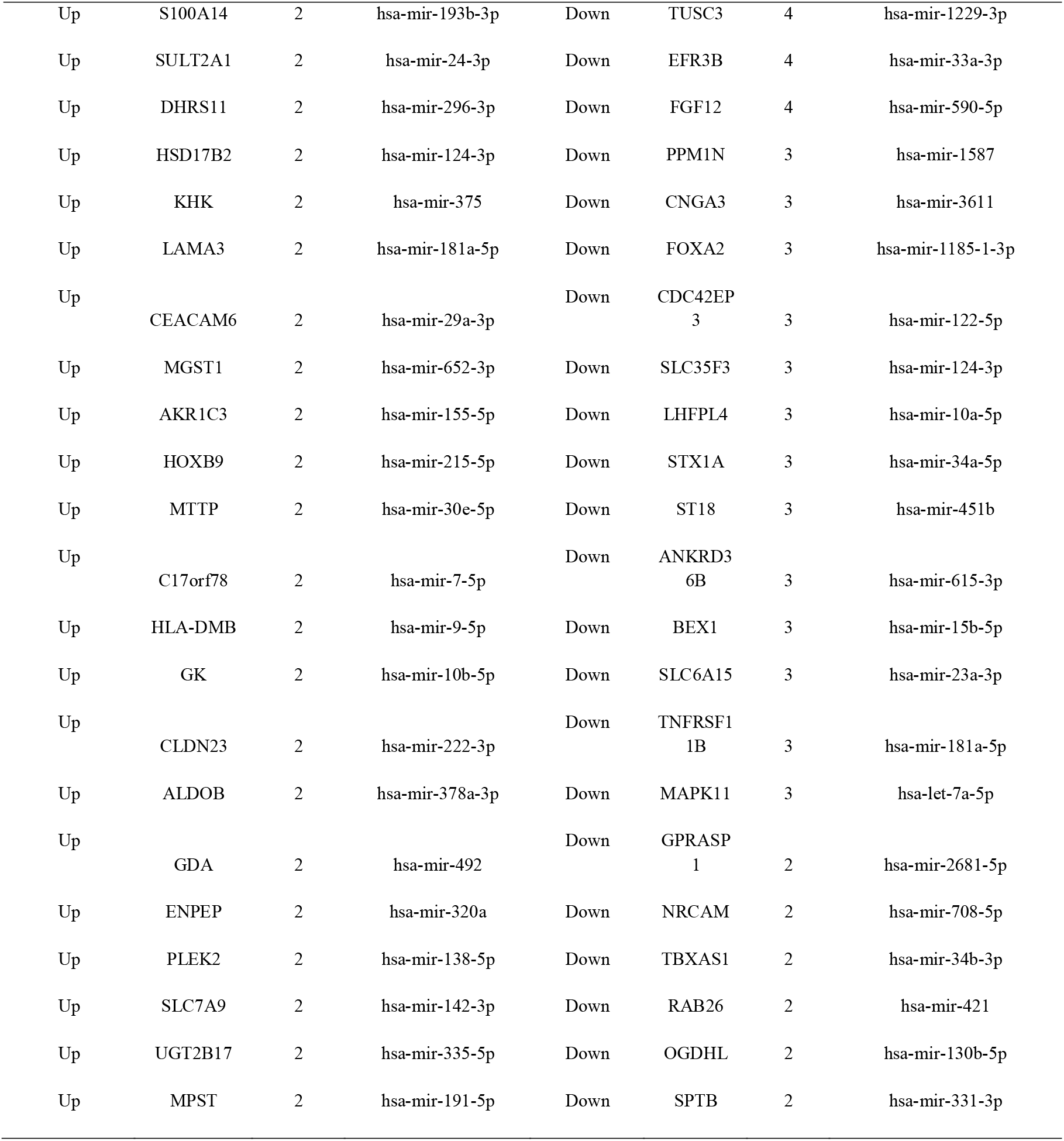
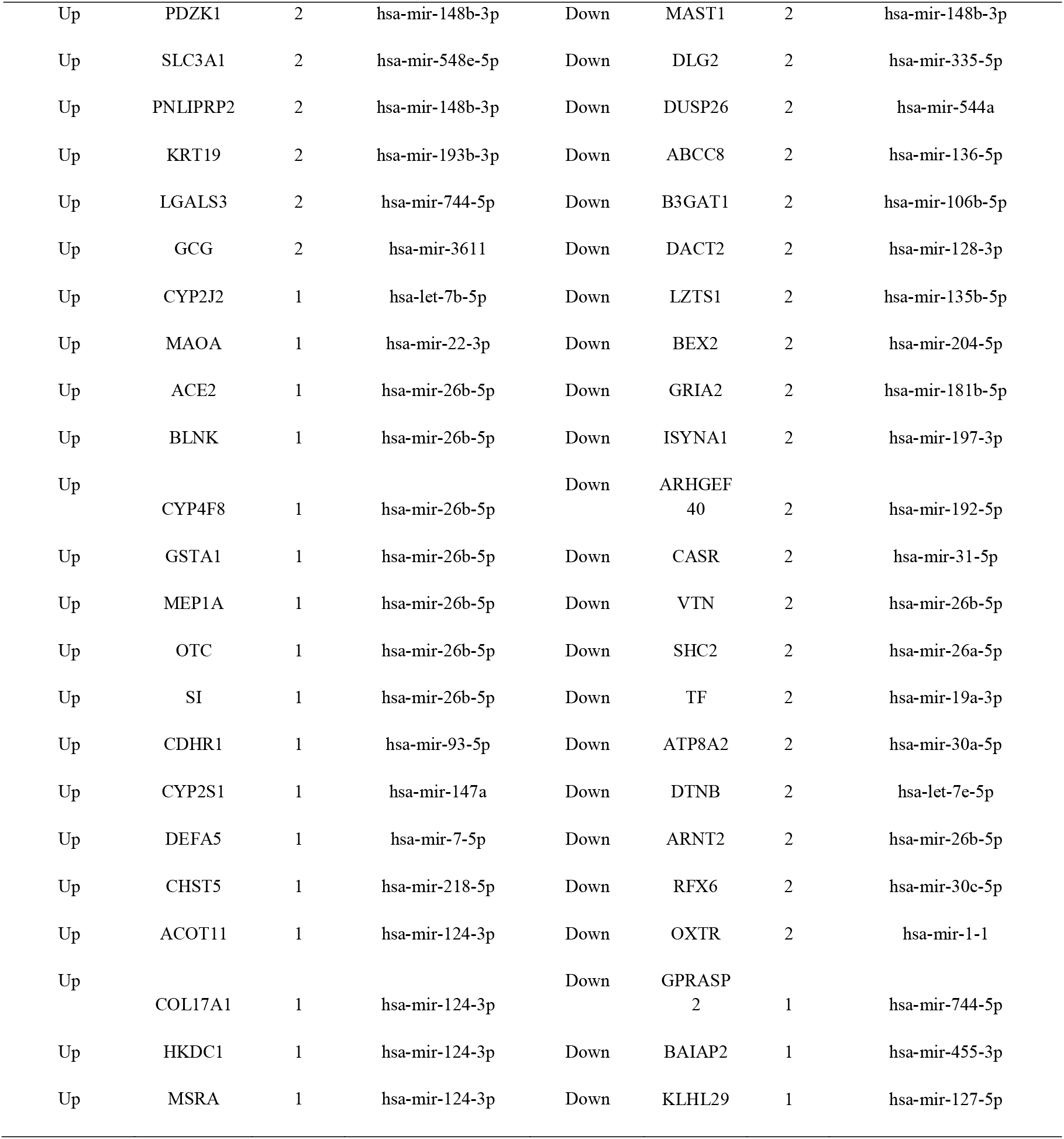
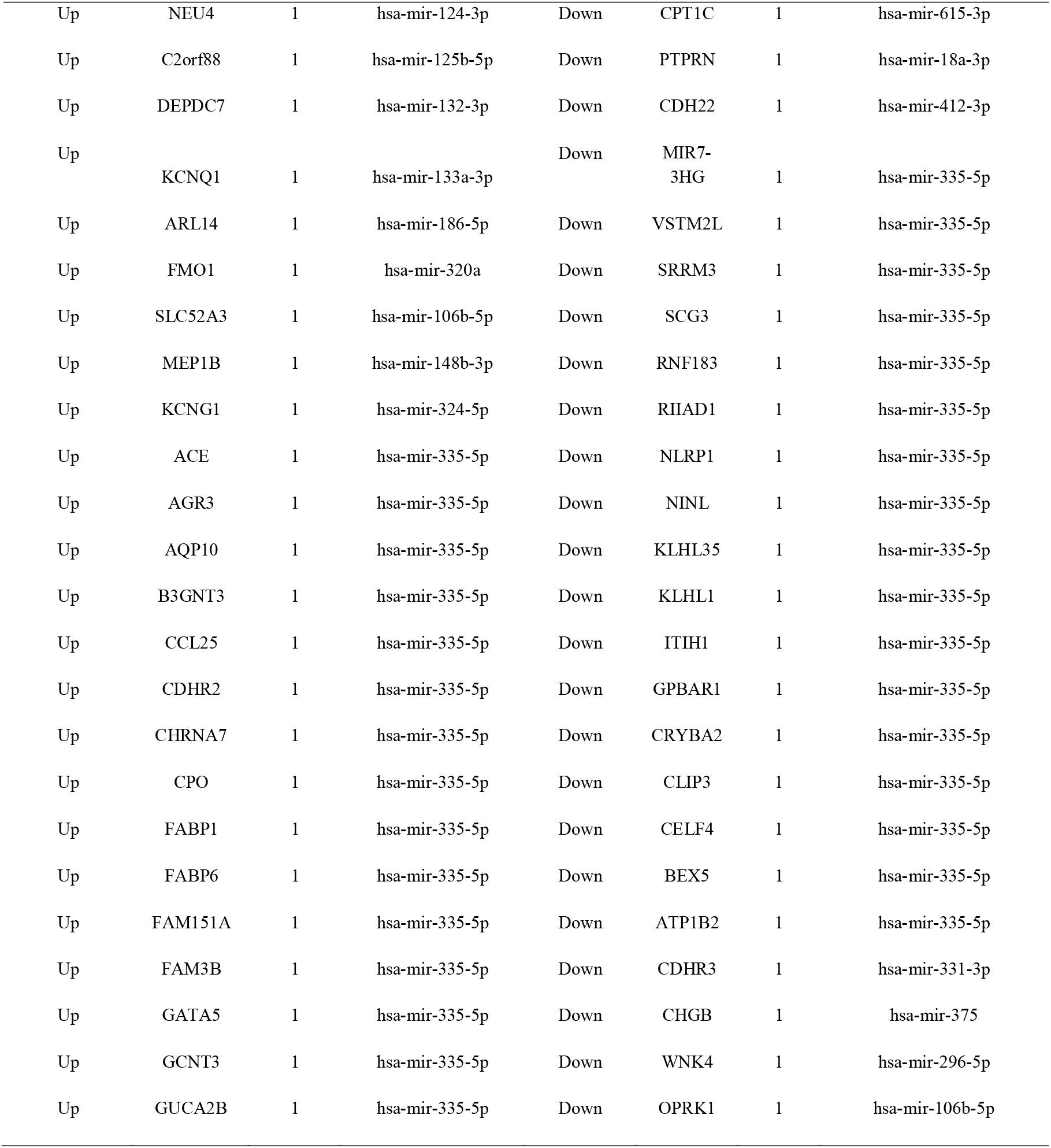
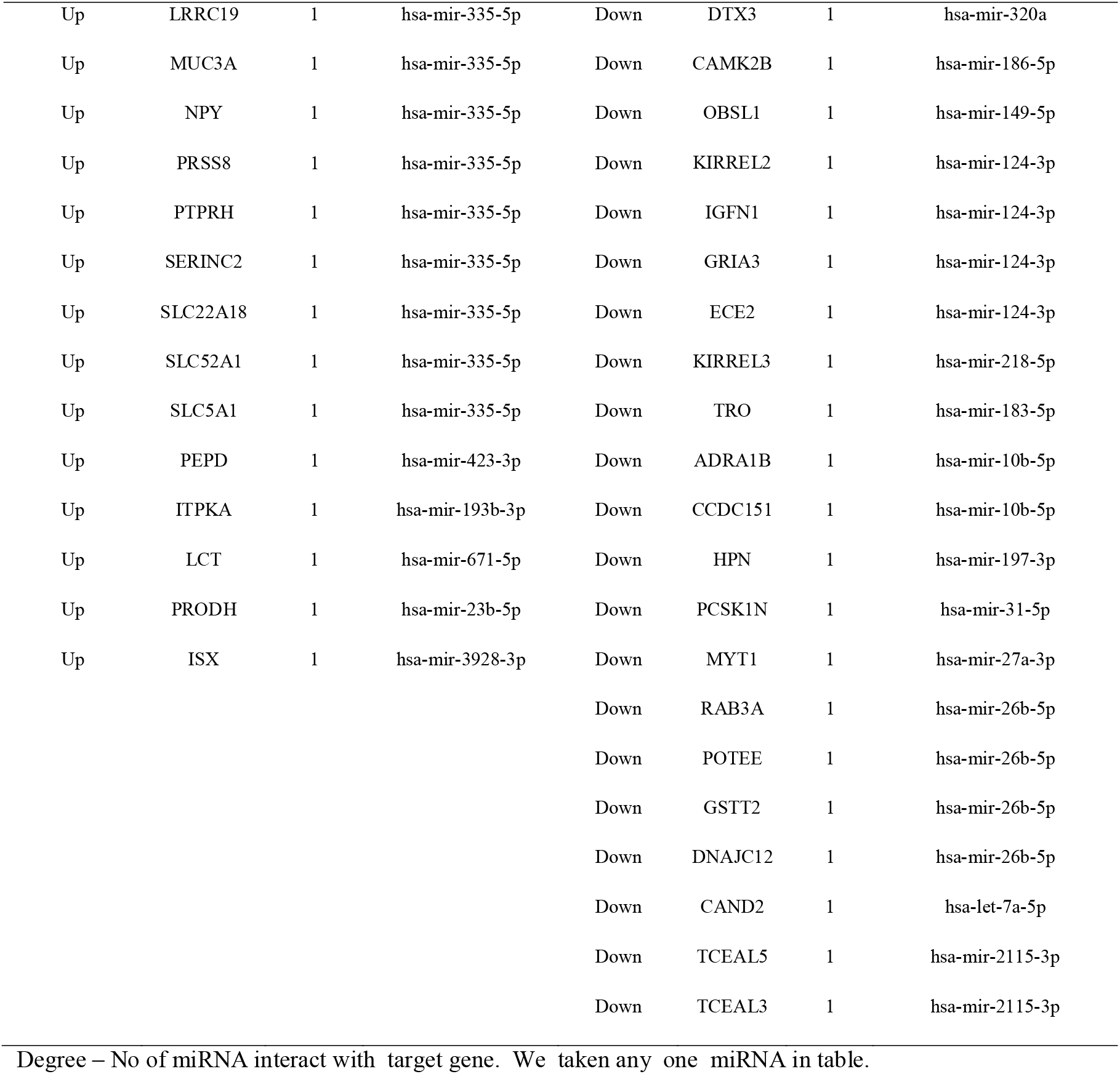
miRNA - target gene interaction table

### Construction of the target gene - TF network

TFs play a key role in the control of gene expression, so the TFs that controlled up and down regulated genes were also predicted (Fig. 7A and Fig. 7B). Top 5 up regulated targeted genes such as CYBRD1 interacts with 158 TFs, MUC13 interacts with 152 TFs, UGT1A6 interacts with 128 TFs, ATP1A1 interacts with 119 TFs and YIF1B interacts with 114 TFs are listed in Table 9. These target genes in this network were enriched in transmembrane transport of small molecules, O-linked glycosylation of mucins, chemical carcinogenesis, carbohydrate digestion and absorption. Meanwhile, Top 5 down regulated targeted genes such as TBXAS1 interacts with 186 TFs, HS6ST3 interacts with 165 TFs, RUNX1T1 interacts with 156 TFs, ATP1B2 interacts with 138 TFs and RFX6 interacts with 125 TFs are listed in Table 9. These target genes in this network were enriched in aspirin blocks signaling pathway involved in platelet activation, transcriptional misregulation in cancer, cardiac conduction and regulation of beta-cell development.

**Figure 7.**
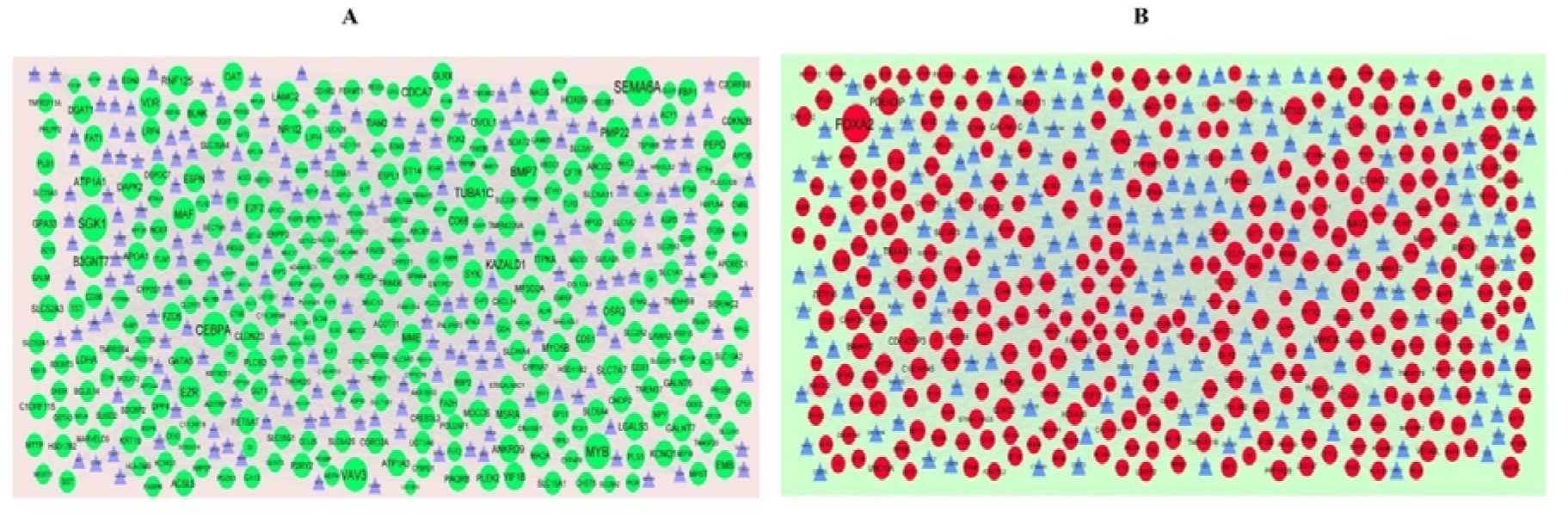
(A) The network of up regulated genes and their related TFs. (Blue triangle - TFs and green circles-target up regulated genes) (B) The network of down regulated genes and their related TFs. (Blue triangle - TFs and red circles - target down regulated genes)

**Table 9.**
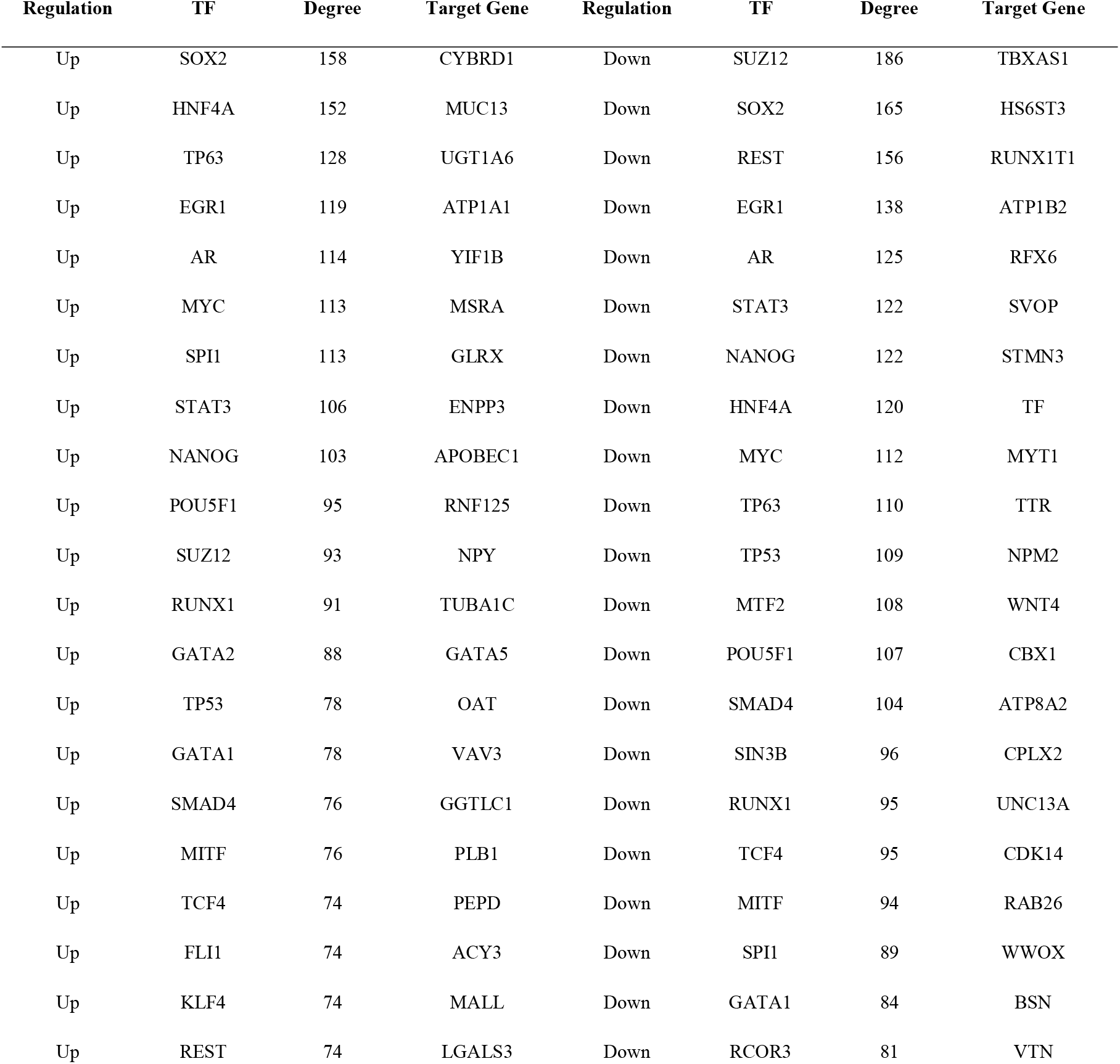
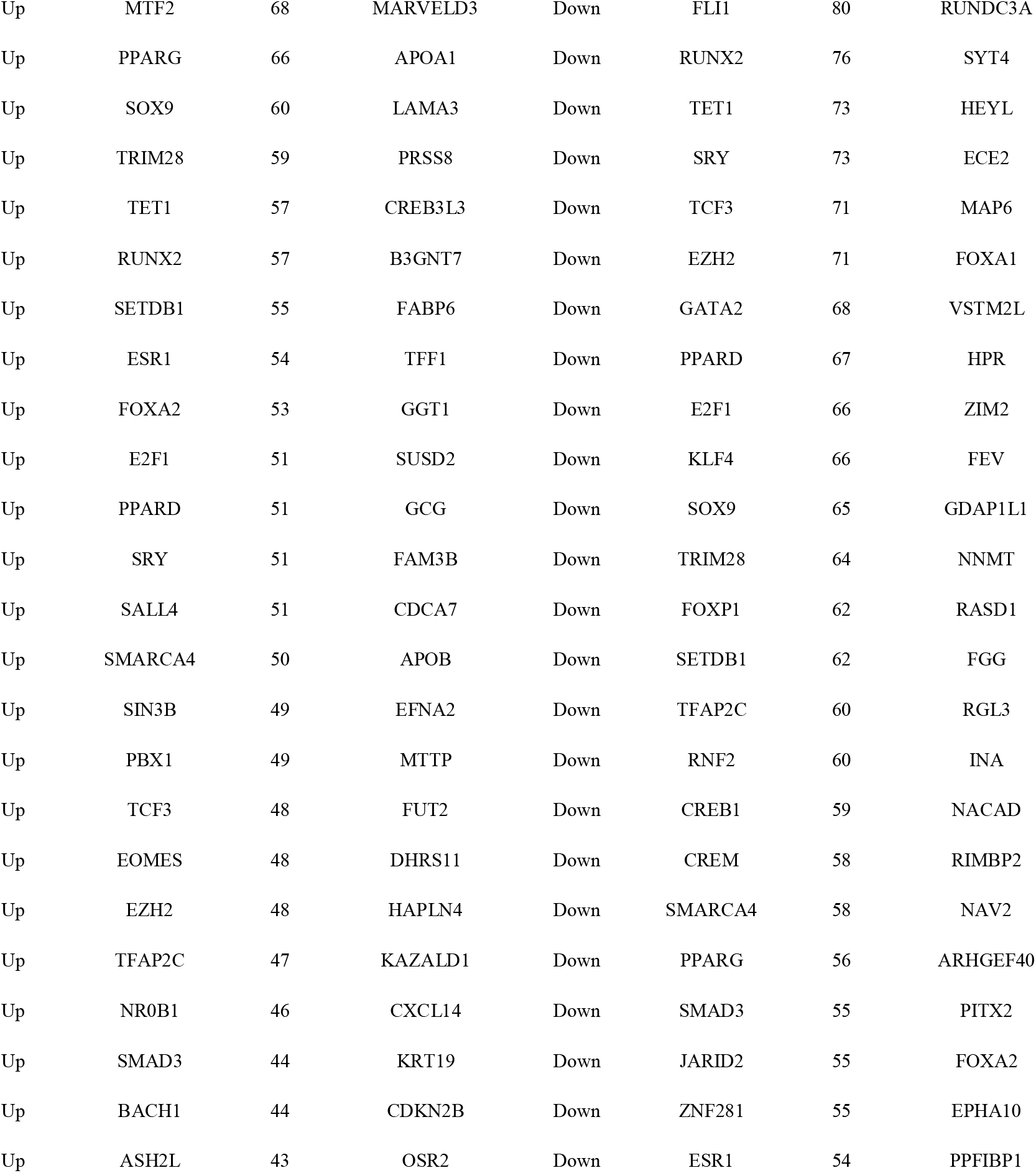
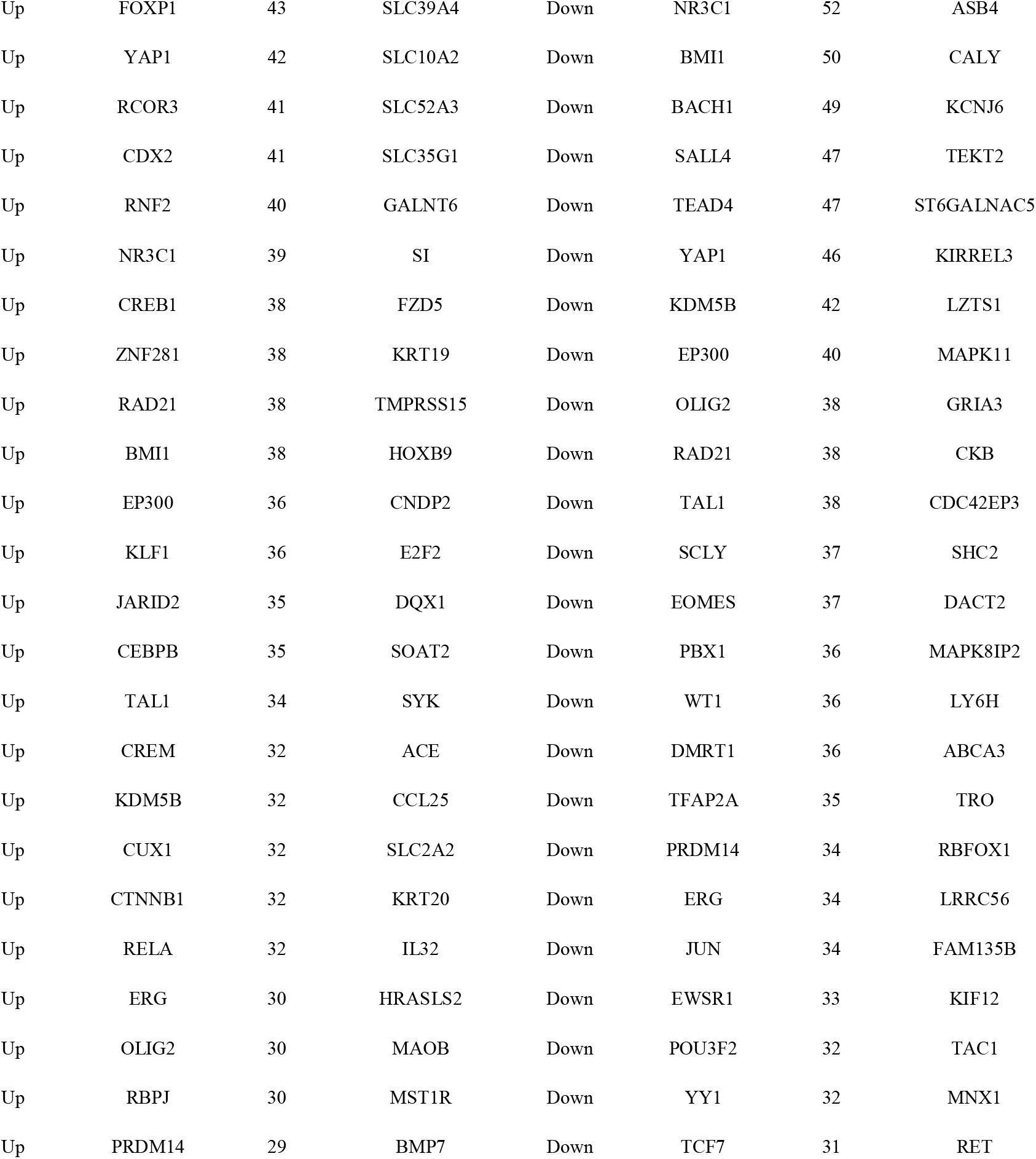
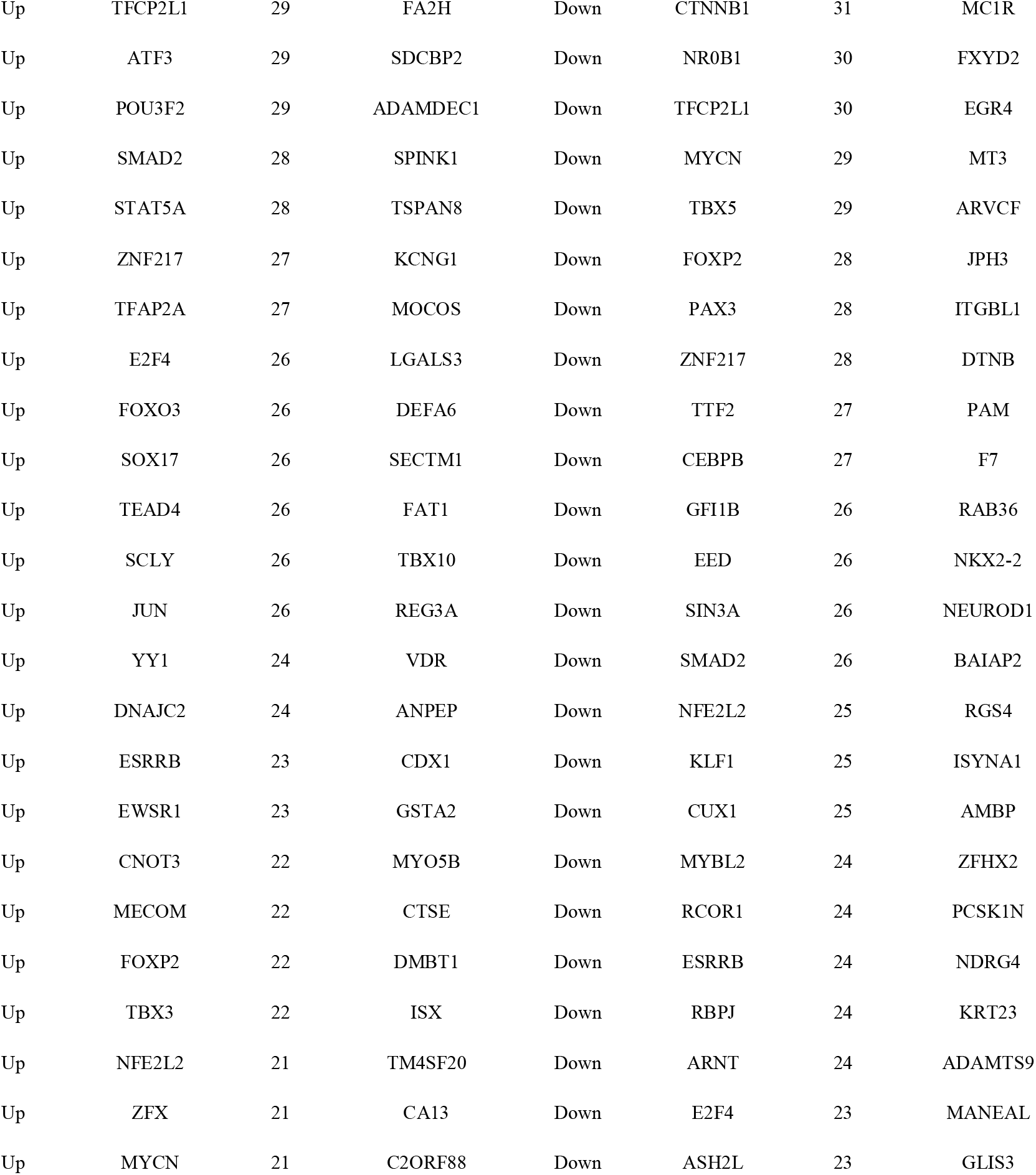
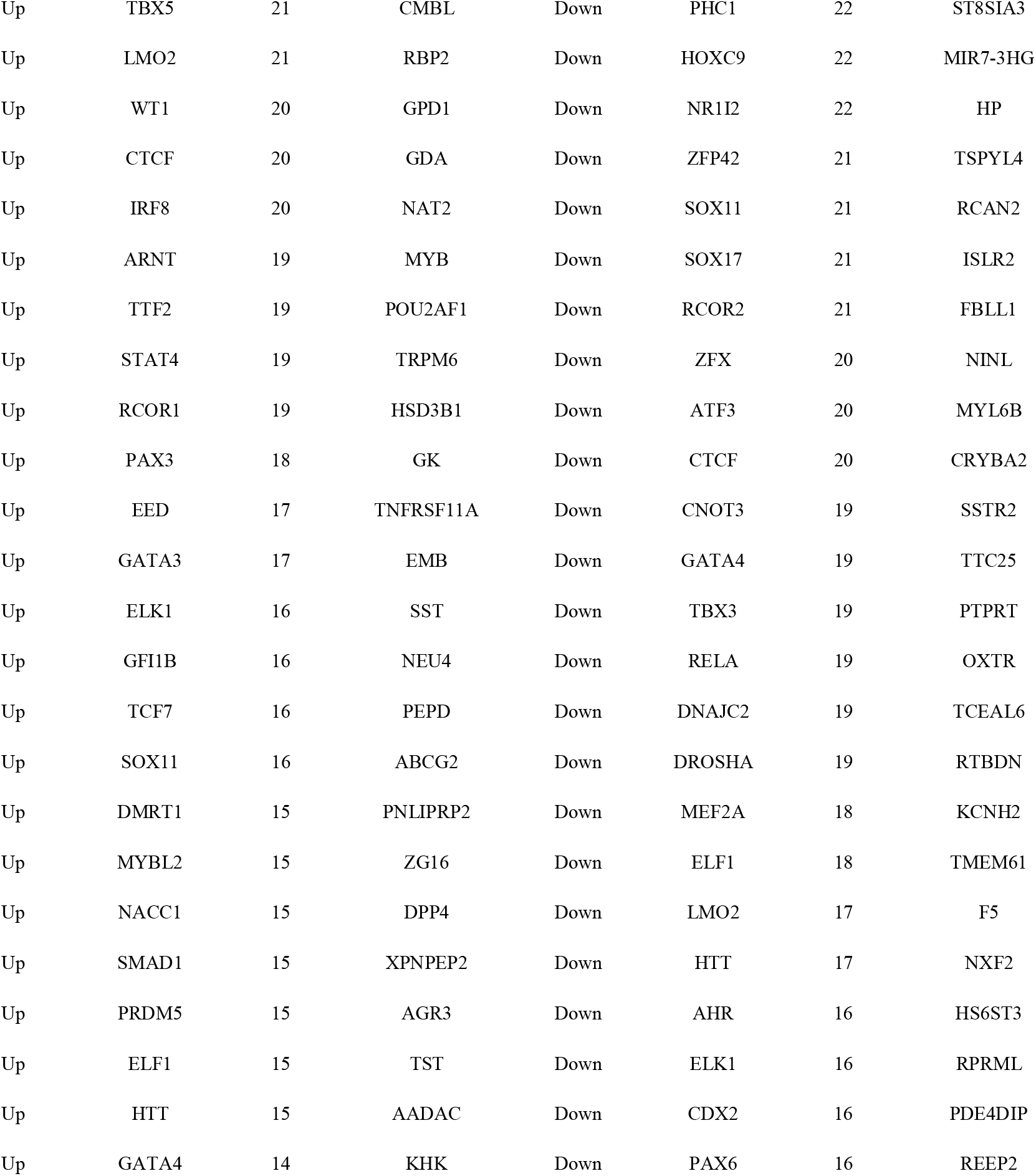
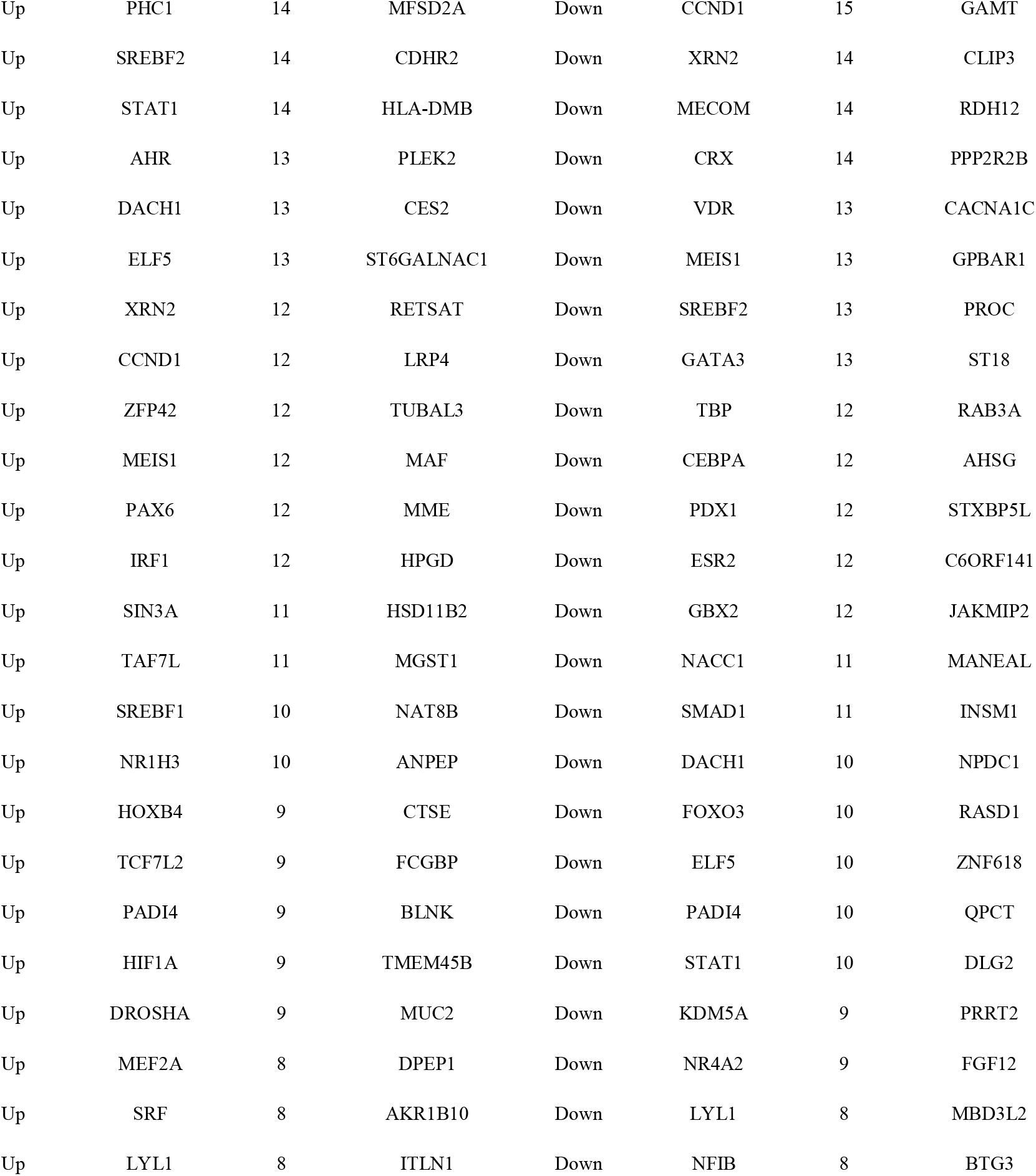
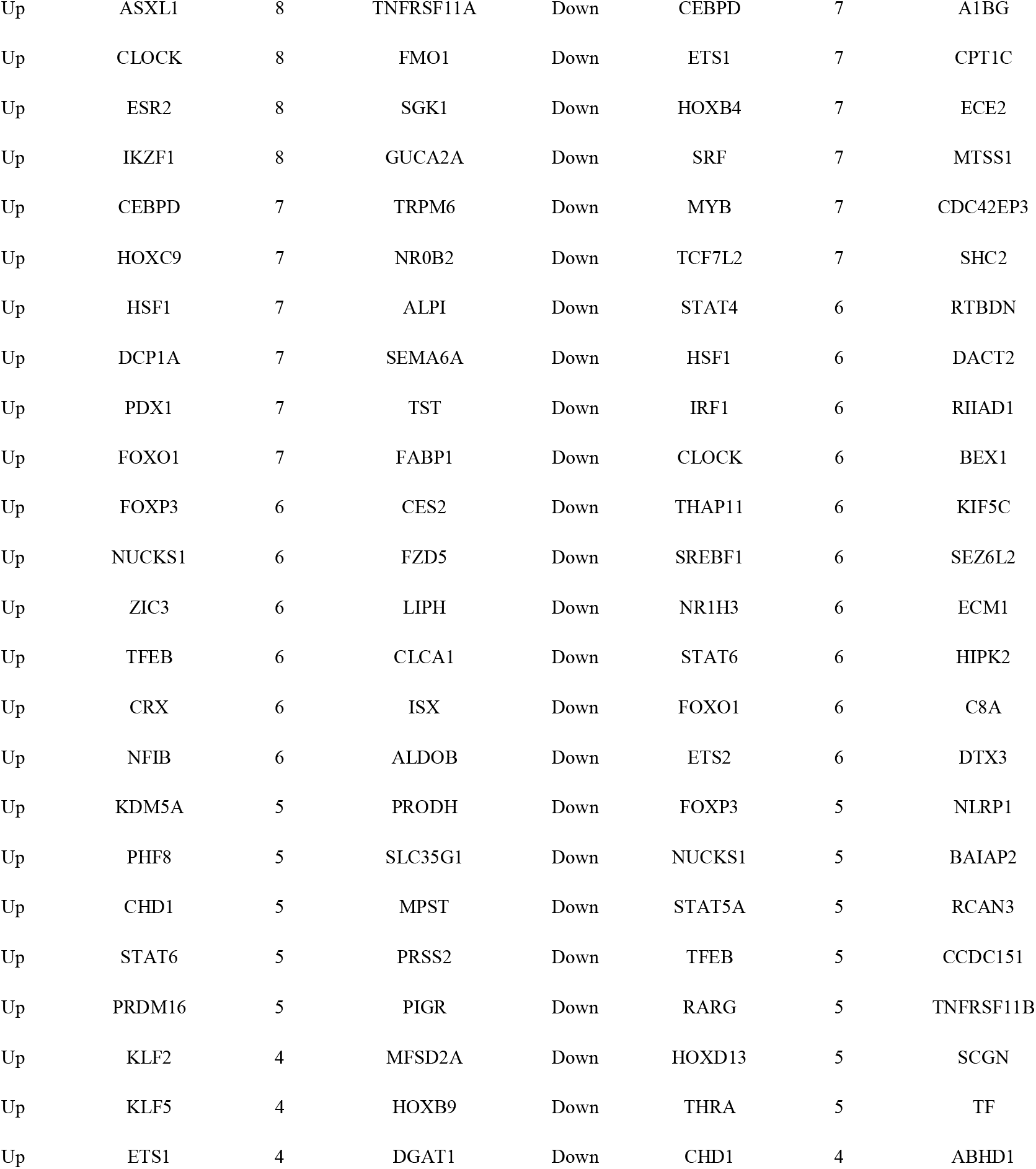
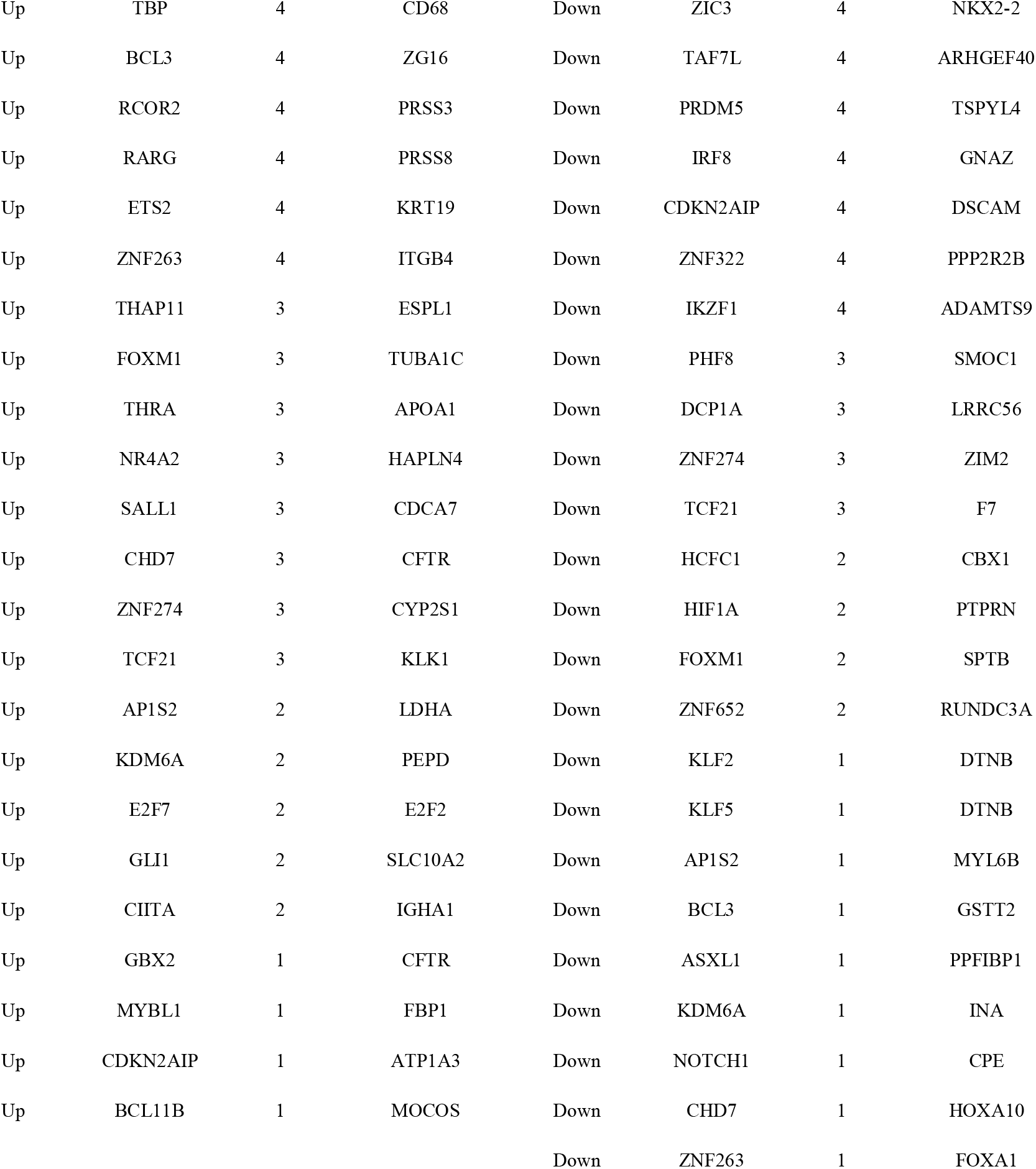
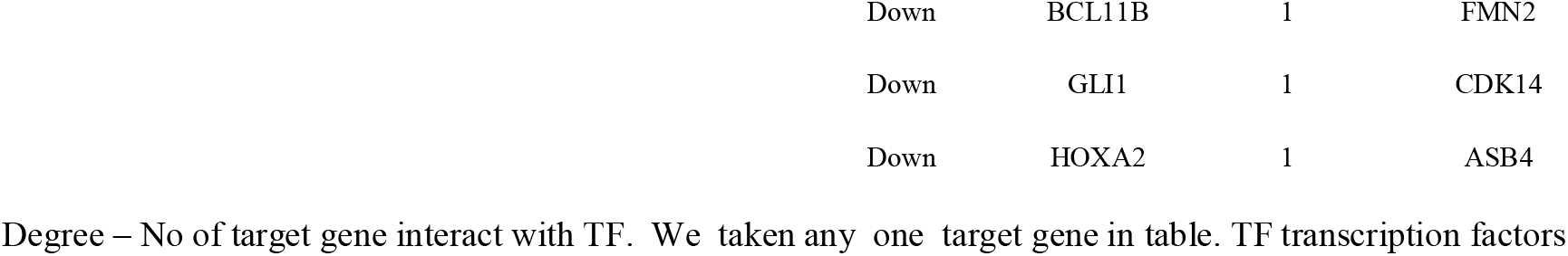
TF - target gene interaction table

**Table 10.**
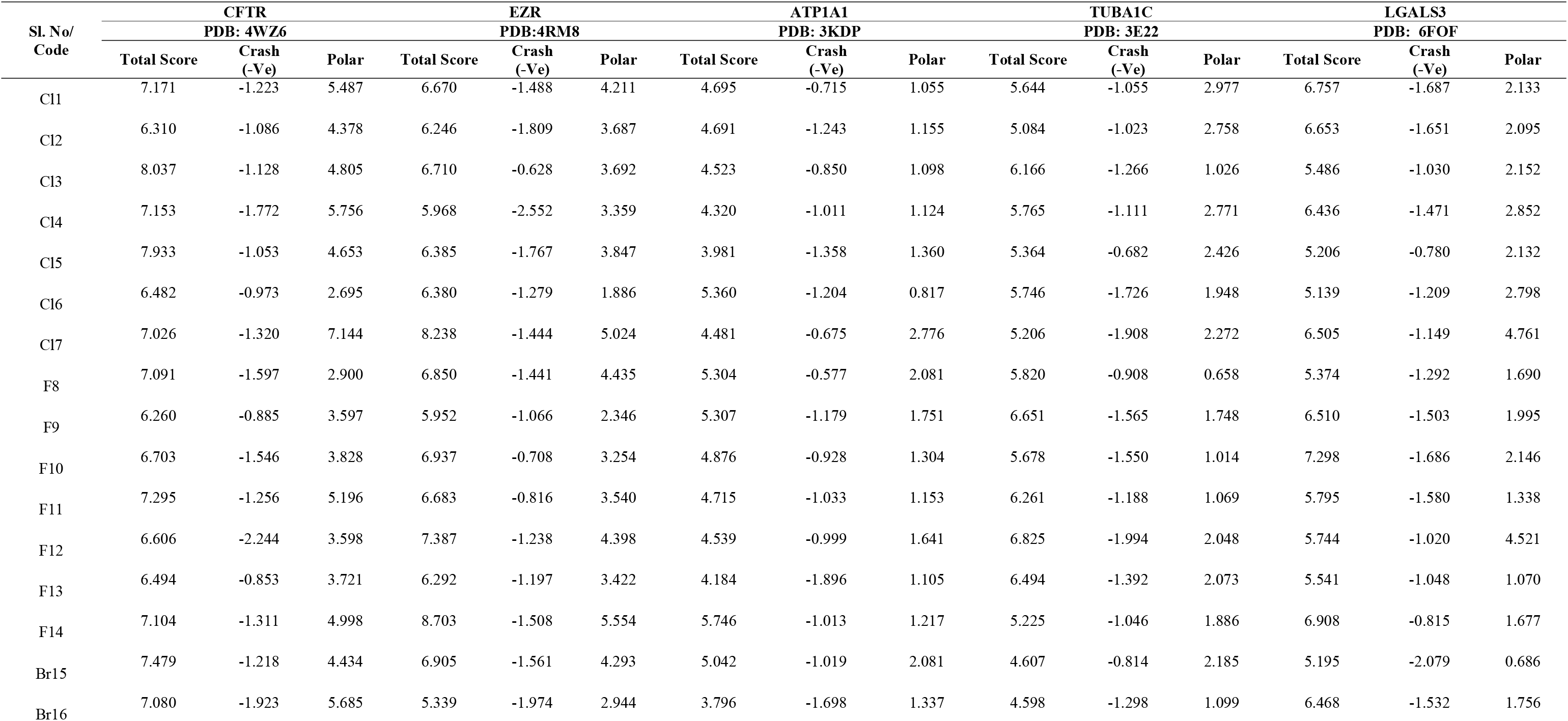
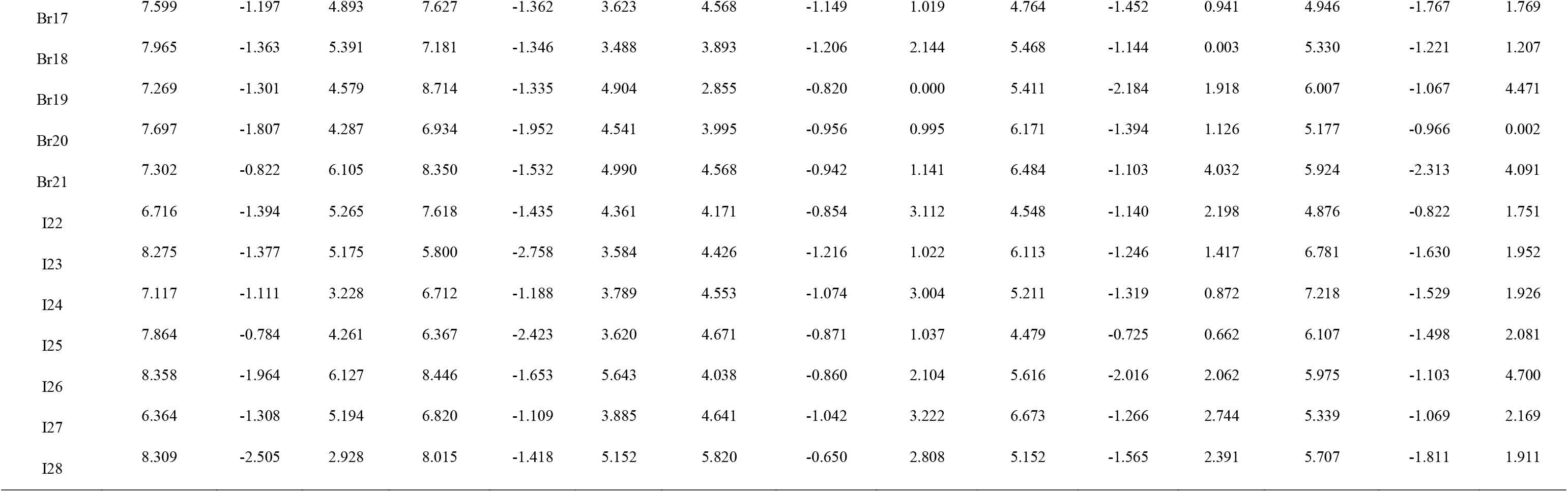
Docking results of Designed Molecules on Over expressed Proteins

### Validations of hub genes

The prognostic effects of the 10 hub genes (up and down regulated genes) in the PPI network were evaluated in PROGgeneV2. It showed that higher mRNA expressions of ATP1A1 (HR: 0.48 [0.35–0.67] (Fig. 8IA), P = 1.37e-05), LGALS3 (HR: 0.77 [0.62–0.77] (Fig. 8IB), P = 0.024321), SYK (HR: 0.65 [0.57–0.75], P = 0) (Fig. 8IC), VDR (HR: 0.77 [0.69–0.87], P = 1.45e-05) (Fig. 8ID), OBSL1 (HR: 0.56 [0.38–0.84], P = 0.00487(Fig. 8IE)), KRT40 (HR: 0.66 [0.58–0.76], P = 0) (Fig. 8IF), NINL (HR: 0.52 [0.36–0.74], P = 0.00034) (Fig. 8IG) and PPP2R2B (HR: 0.75 [0.64–0.89], P = 0.000777) (Fig. 8IH) were associated with better overall survival in NET patients), whereas higher expression of LDHA (HR: 2.52 [2.05–3.1], P = 0) (Fig. 8IIA) and WWOX (HR: 4.87 [3.48–6.82], P = 0) (Fig. 8II B) were related to poor overall survival in NET patients. ROC curve analysis was used to assess the ability of the expression levels of ATP1A1, LGALS3, SYK, VDR, LDHA, OBSL1, KRT40, NINL, PPP2R2B and WWOX to distinguish between metastatic NET and normal control. The results suggested that AUC values of ATP1A1, LGALS3, SYK, VDR, LDHA, OBSL1, KRT40, NINL, PPP2R2B and WWOX were 0.973, 0.991, 0.955, 0.976, 0.994, 0.042, 0.930, 0.921, 0.939 and 0.942 (Fig. 9I). These results had significant potential to distinguish between metastatic NET and normal control. However, the AUCs in the current investigations are relatively high and may be sufficient for each molecule on its own to predict recurrence. The expression levels of certain central genes identified, including ATP1A1, LGALS3, SYK, VDR, LDHA, OBSL1, KRT40, NINL, PPP2R2B and WWOX, were further examined by RTL PCR. In the metastatic NET, genes such as ATP1A1, LGALS3, SYK, VDR and LDHA were significantly up regulated as compared with normal control (Fig. 9IIA – 9II E), while in metastatic NET, genes such as OBSL1, KRT40, NINL, PPP2R2B and WWOX were significantly down regulated as compared with normal control (Fig. 9IIF – 9IIJ). The results indicated that, consistent with the micro array results.

**Figure 8.**
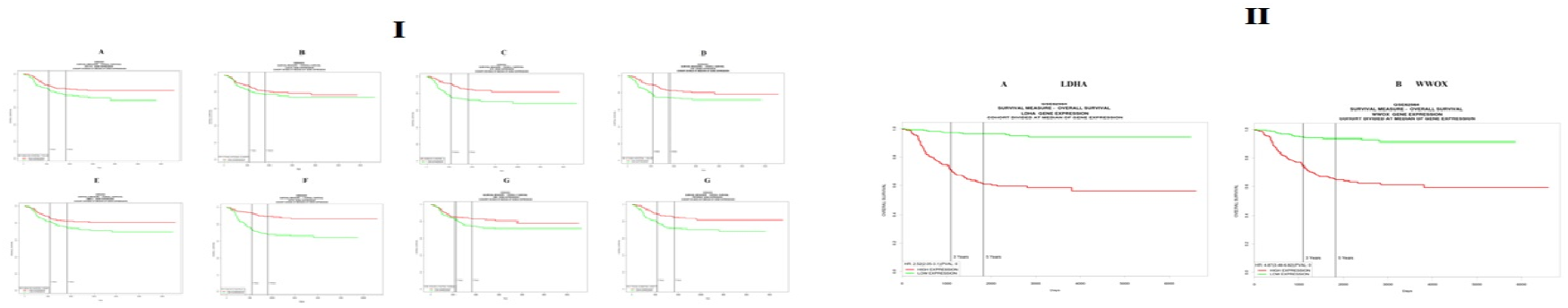
(I) Kaplan-Meier survival curves using GSE62564 data validate the prognostic value of genes have favorable overall survival in NETs. (Green – Low expression; Red-high expression) (A = ATP1A1; B = LGALS3; C = SYK; D = VDR; E = OBSL1; F = KRT40; G = NINL; H = PPP2R2B) (II) Kaplan-Meier survival curves using GSE62564 data validate the prognostic value of genes have worse overall survival in NETs. (Green – Low expression; Red-high expression) (A = LDHA; B = WWOX)

**Figure 9.**
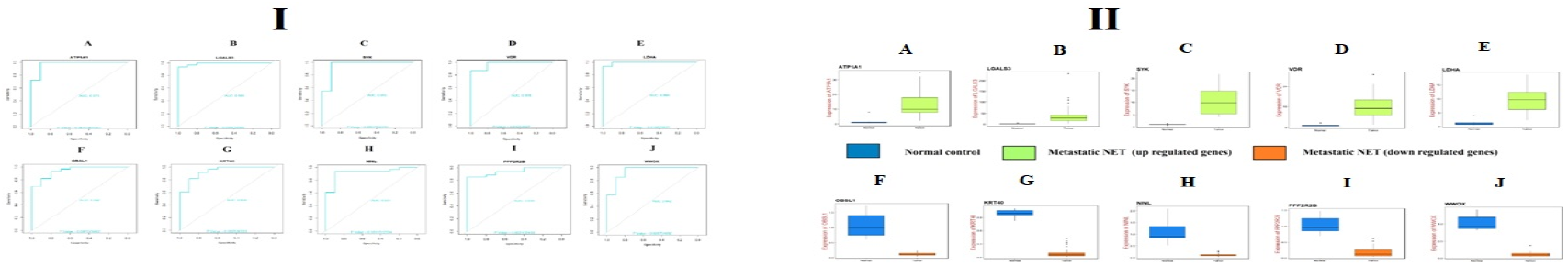
(I) ROC curve validated the sensitivity, specificity of hub genes as a predictive biomarker for NET prognosis. A) ATP1A1 B) LGALS3 C) SYK D) VDR E) LDHA F) OBSL1 G) KRT40 H) NINL I) PPP2R2B J) WWOX (II) Validation of hub genes (up and down regulated) by RT-PCR. A) ATP1A1 B) LGALS3 C) SYK D) VDR E) LDHA F) OBSL1 G) KRT40 H) NINL I) PPP2R2B J) WWOX

### Molecular docking studies

The docking simulation was performed in the current study to identify the active site conformation and major interactions responsible for complex stability with the binding sites receptor. Docking experiments on designed molecules containing purine heterocyclic ring were performed using Sybyl X 2.1 software. Molecules containing the heterocyclic ring of purine were constructed based on the nucleotide structure and are most widely used alone or in conjunction with other antineoplastic drug. Most neuroendocrine tumors occur in the lungs, appendix, small intestine, rectum and pancreas. When neuroendocrine cells develop changes (mutations) in their DNA, neuroendocrine tumours arise. There are instructions in the DNA within a cell that tell the cell what to do. The modifications suggest that the neuroendocrine cells replicate rapidly and form a tumour. The over expressed genes in neuroendocrine tumours and their X-RAY crystallographic structure of one protein from each over-expressed genes such as CFTR (Cystic fibrosis transmembrane conductance regulator), EZR (Ezrin), ATP1A1 (ATPase Na+/K+ transporting subunit alpha 1), TUBA1C (Tubulin alpha 1c) and LGALS3 (Galectin 3) and their co-crystallized PDB code of 4WZ6, 4RM8, 3KDP, 3E22 and 6FOF respectively were selected for docking. In order to identify the possible molecule, an analysis of the designed molecules was carried out. The key molecules that have been designed have obtained a C-score greater than 5 and above are said to be active. A total of 28 molecules were designed shown in Fig. 10 and performed docking studies, few molecules obtained excellent binding energy (C-score) greater than 8 and few molecules obtained optimum binding score 4-4.9, only one molecule with one protein obtained less binding score 2.0-3.0 respectively. The molecules obtained good binding score more than 5-8 and are I26, I28, I24, Cl4, Br16, I23, Br20, Br18, Br15, Br21, F9, Br19, Cl1, Cl2, I25, F14, F8, Br17, Cl7 with 4WZ6 the molecules Br19, F14, Br16, Br21, Cl7, I28, Br18, I22, F12, Br16, Cl1, Br20, Br15, F8, I27, I25, Cl4, F9, Cl1, Cl5, Cl6, I23, F13, Cl3 with 4RM8, the molecules F14, Cl6, F10, F8, Br15, I28, Cl7, F11, F12, F13, I27, Cl3 with 3KDP, the molecules F12, I27, F10, F13, Br21, F9, Br20, Cl4, I24, Cl2, Cl6, F11, Cl1, I26, Br16, Br19, Cl5, F14, I25, Cl7, I28, Cl3, F8 with 3E22 and the molecules F11, I25, F14, I24, Cl1, Cl3, F10, Cl7, Br17, Cl2, I23, Br19, I26, Br21, F9, F12, I28, F13, Cl4, F8, I27, Br16, Cl5, Br15, Br20, Cl6 with 6FOF respectively. The molecules obtained optimum binding score less than 5 (3.0-4.9) are Cl4, F9, I23, Br21, Br18, I25, Cl4, I24, Cl2, I22, I26 with 3KDP, the molecules Br18, Br15, Cl7, I22, I23 with 3E22 and the molecules Br18, I22 with 6FOF respectively. One molecule Br19 obtained very less binding score of 2with protein 3KDP respectively the values are depicted in Table 1. With all proteins the molecule Cl4, Cl6, F14, F8, Cl7, F11, F12, F13, I27, Cl3, F10 and I28 obtained good binding score.The hydrogen bonding interactions and other bonding interactions with amino acids with protein code 4RM8of molecule Br17 are depicted by 3D and 2D (Fig.11 and Fig. 12).

**Figure 10.**
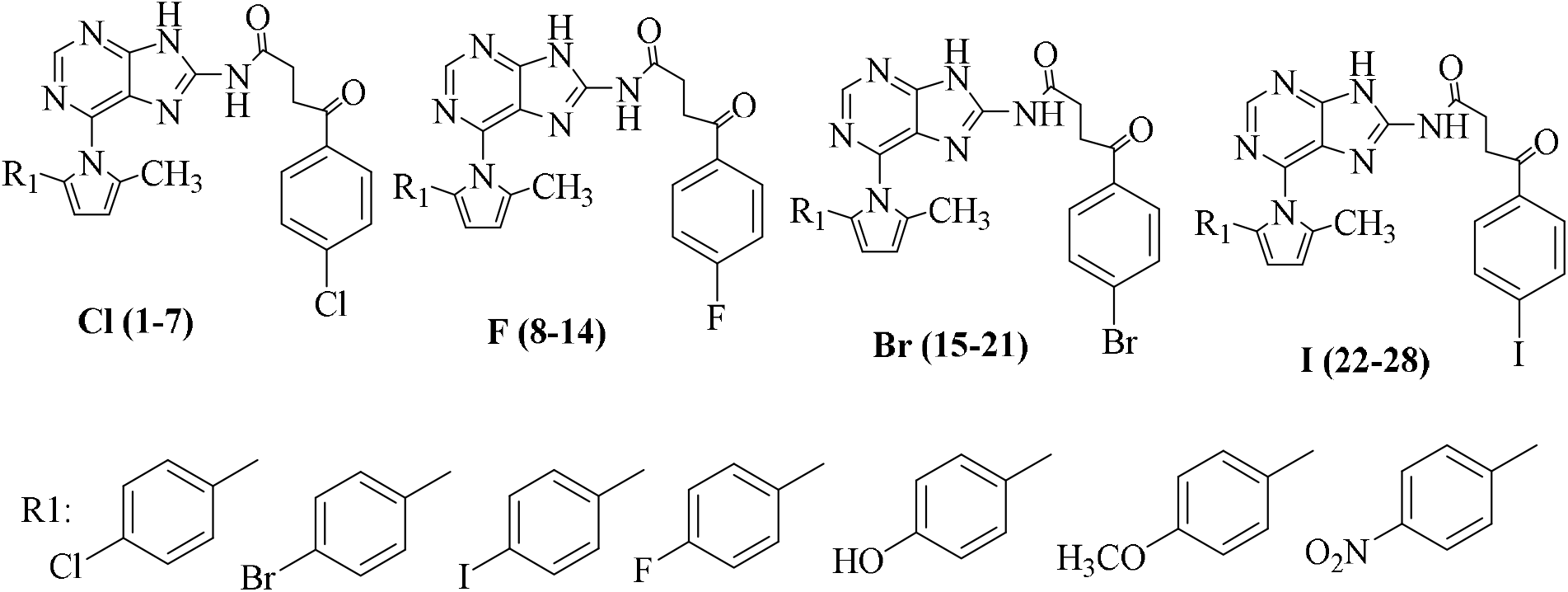
Structures of Designed Molecules

**Figure 11.**
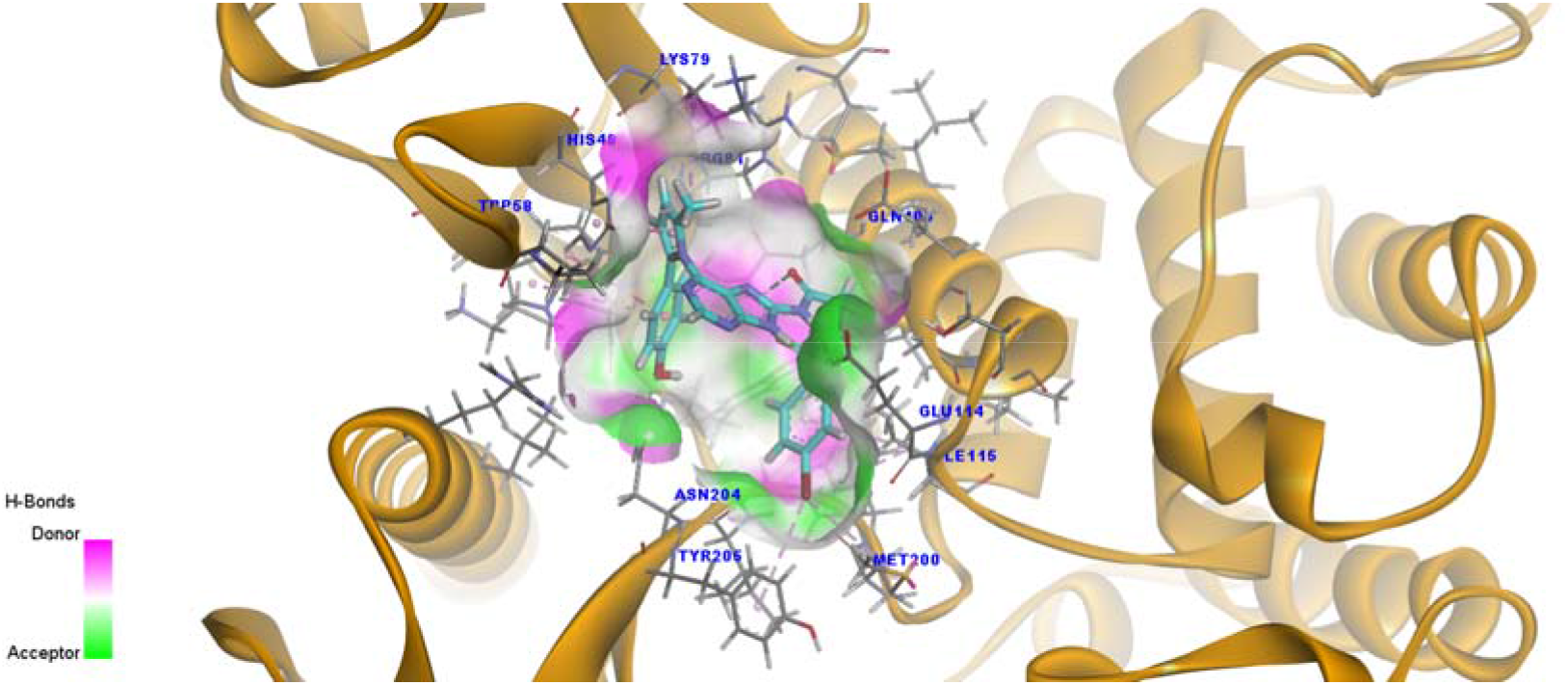
3D Binding of Molecule Br17with 4RM8

**Figure 12.**
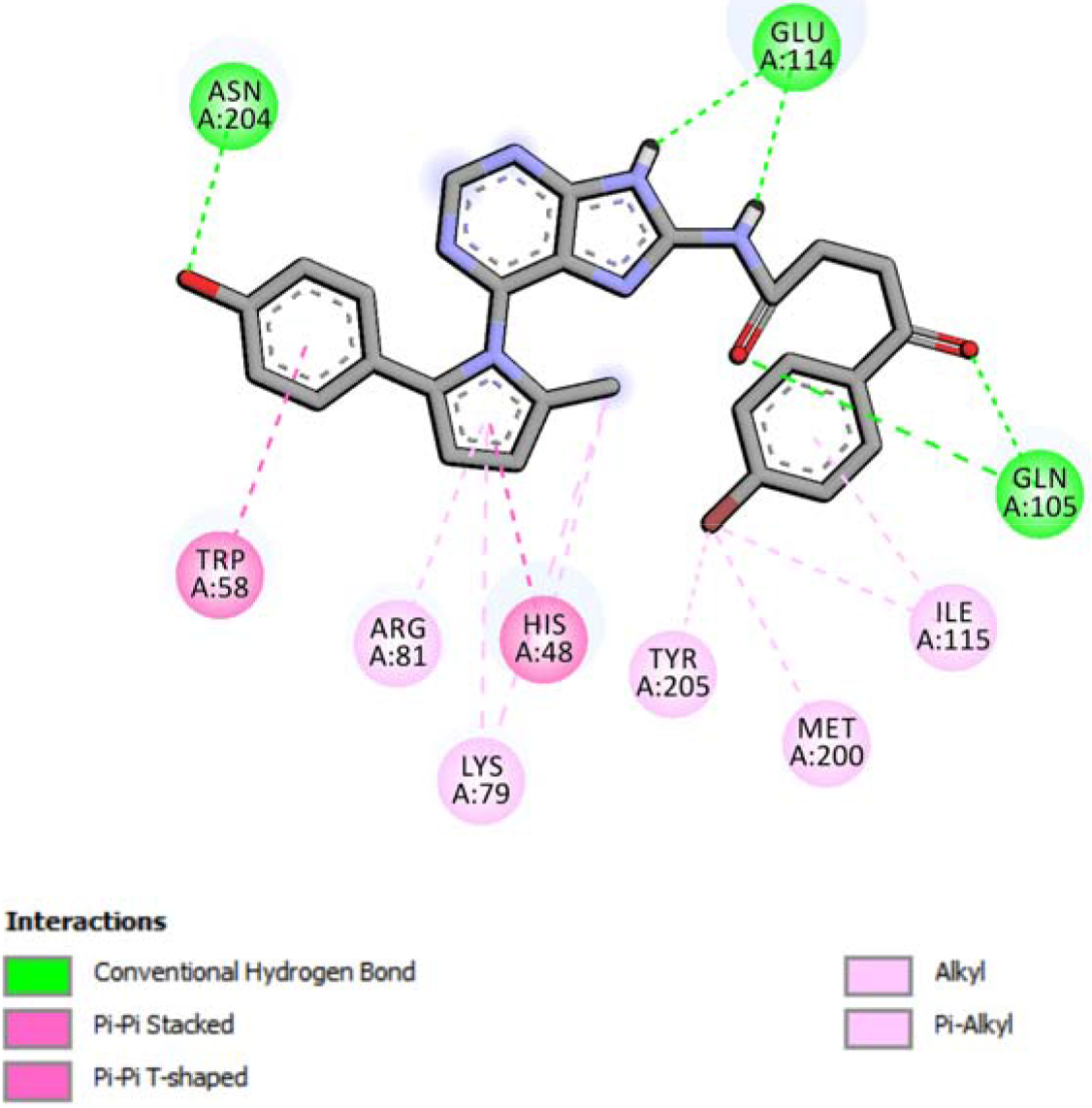
2D Binding of Molecule Br17with 4RM8

## Discussion

Various molecular methods have been used in the attempt to illuminate the heterogeneous nature of NET, resolve the mechanisms behind its progression, and introduce novel therapeutic and prognostic targets [56]. Microarray analysis or genome scale analysis has proven to be strong in numerous experimental settings and has the possible to feature the dynamic molecular diversity encountered during cancer development [57]. Since studies investigating gene expression changes during development of NET are lacking, this study sought to identify DEGs in NET that may be exploited as specific prognostic and therapeutic targets.

In the present study, a total of 459 up regulated and 453 down regulated genes were identified between patients with metastatic NET and on normal (controls). Genes such as TIAM2 [58] and SCIN (scinderin) [59] were associated with invasion of various types cancer cells, but these genes might be involves in invasion of NET cells. Polymorphic genes such as SLC6A4 [60] and SLC23A1 [61] were responsible for pathogenesis of various cancer types, but these polymorphic genes might be linked with development of NET. Increased expression of APOBEC1 was important for pathogenesis of colon cancer [62], but elevated expression of this gene may be identified with growth of NET. TTR (transthyretin) was responsible for progression of NET [63]. Methylation inactivation of tumor suppressor genes such as OGDHL (oxoglutarate dehydrogenase like) [64], CAMK2B [65] and RPRML (reprimo like) [66] were associated with progression of various cancer types, but loss of these genes might be liable for development of NET. Loss of tumor suppressor SRCIN1 was diagnosed with growth of gastric cancer [67], but loss of this gene might be linked with development of NET.

In pathway enrichment analysis for DEGs was performed. Genes such as ALDOB (aldolase, fructose-bisphosphate B) [68], KHK (ketohexokinase) [69], CHRNA7 [70], GSTA3 [71], APOA1 [72], CEBPA (CCAAT enhancer binding protein alpha) [73], PCK1 [74], ATP1A1 [75], CLCA1 [76], EMB (embigin) [77], SLC22A18 [78], FUT5 [79] and ITLN1 [80] were linked with progression of various cancer types, but these genes might be identified with growth of NET. Polymorphic genes such as GSTA1 [81], GSTA2 [82], MGST1 [83], NAT2 [84], SULT1A2 [85], SULT2A1 [86], UGT1A6 [87], UGT2B15 [88], UGT2B17 [89], ABCC2 [90], FUT2 [91] and APOB (apolipoprotein B) [92] were liable for progression of various cancer types, but these polymorphic genes might be diagnosed with growth of NET. Genes such as G6PC [93] and SGK1 [94] were associated with proliferation of various cancer cells types, but these gene might be linked with proliferation of NET cells. Increased expression of genes such as SLC2A2 [95], ABCG5 [96], SLC39A4 [97], SLC3A1 [98], SLC7A7 [99], LGALS3 [100], MUC13 [101], MUC17 [102], MUC2 [103], REG1A [104], REG3A [105], REG3G [106] and SEMA6A [107] were responsible for advancement of various cancer types, but high expression of these genes might be liable for growth of NET. Genes such as ABCB1 [108], ABCG2 [109] and SLC26A3 [110] were linked with drug resistance in various cancer types, but these genes might be associated with chemo resistance in NET. CFTR (cystic fibrosis transmembrane conductance regulator) was responsible for progression of NET [111]. Methylation inactivation of tumor suppressor genes such as MYO5B [112], FUT3 [113], CPS1 [114] and FBP1 [115] were identified with progression of various cancer types, but loss of these genes might be important for progression of NET. Genes such as SLC2A5 [116] and FUT6 [117] were associated with invasion of various cancer types, but these genes might be responsible for invasion of NET cells. Stimulation of PCK2 was associated with development of lung cancer [118], but activation of this gene might be linked with progression of NET. Polymorphic genes such as ATP1B2 [119] and F5 (coagulation factor V) [120] were responsible for progression of breast cancer, but these polymorphic genes might be linked with pathogenesis of NET. Genes such as FXYD2 [121], RAB3A [122], ALB (albumin) [123], A1BG [124], AHSG (alpha 2-HS glycoprotein) [125], FGG (fibrinogen gamma chain) [126], SCG3 [127], TF (transferrin) [128], CKB (creatine kinase B) [129] and F7 (coagulation factor VII) [130] were linked with pathogenesis of various cancer types, but high expression of these genes might liable for progression of NET. Genes such as FOXA1 [131] and FOXA2 [132] were linked with development of NET. Genes such as ECM1 [133] and KCNH2 [134] were involved in invasion of various cancer cells types, but these genes might be involved in invasion of NET cells.

In GO enrichment analysis for DEGs was performed. Polymeric genes such as ACE (angiotensin I converting enzyme) [133], CD36 [136], FABP2 [137], MFSD2A [138], SLC52A3 [139] and LCT (lactase) [140] were responsible for advancement of various cancer types, but these polymorphic genes might be involved in development of NET. Genes such as ACE2 [141], CA13 [142], SLC10A2 [143] and KCNQ1 [144] were liable for progression of various cancer types, but these genes might be linked with development of NET. Increase expression of genes such as ACSL5 [145], FABP1 [146], FABP6 [147], P2RY2 [148], PDZK1 [149], EZR (ezrin) [150] and SI (sucrase-isomaltase) [151] were diagnosed with growth of various cancer types, but high expression of these genes might be identified with progression of NET. Methylation inactivation of tumor suppressor genes such as SYK (spleen associated tyrosine kinase) [152] and TNFRSF11A [153] were associated with pathogenesis of various cancer types, but loss of these genes might be diagnosed with growth of NET. Genes such as CPLX2 [154], CHGA (chromogranin A) [155], CHGB (chromogranin B) [155] and CPE (carboxypeptidase E) [156] were responsible for progression of NET. Polymorphic gene CTNND2 was linked with development of prostate cancer [157], but this gene might be involves in growth of NET. Methylation inactivation of tumor suppressor genes such as JPH3 [158], LZTS1 [159] and TAC1 [160] were important for progression of various cancer types, but loss of these genes might be linked with development of NET. Alteration in SLC18A1 was diagnosed with growth of colorectal cancer [161], but modification in this gene might be liable for advancement of NET. Decrease expression of genes such as SLC8A2 [162], ABCA3 [163], PCSK1 [164], APC2 [165], CRMP1 [166] and RCAN3 [167] were identified with progression of various cancer types, but low expression of these genes might be liable for progression of NET. Genes such as PTPRN2 [168] and DCX (doublecortin) [169] were liable for invasion of various cancers cells types, but these genes might be linked with invasion of NET cells. Genes such as FGF13 [170], MTSS1 [171] and NRCAM (neuronal cell adhesion molecule) [172] were identified with pathogenesis of various cancer types, but these genes might be responsible for advancement of NET.

PPI network was constructed and analyzed. LDHA was associated with invasion of hepatic cancer cells [173], but this gene might be responsible for invasion of NET cells. Polymorphic gene HSD17B2 was liable for progression of breast cancer [174], but this polymorphic gene might be associated with development of NET. Methylation inactivation of tumor suppressor genes such as WWOX [175] and PPP2R2B [176] were identified with growth of various cancer types, but loss of these genes might be liable for pathogenesis of NET. CBX1 was liable of advancement of hepatocellular carcinoma [177], but this gene might be linked with development of NET. PHLDB2 was responsible for invasion of colon cancer cells [178], but this gene might be linked with invasion of NET cells. Decrease expression of HIPK2 was identified with growth of colorectal cancer [179], but low expression was this gene might be important for pathogenesis of NET. TUBA1C, C15orf48, ACOT11, PLA2G12B, DHDH (dihydrodiol dehydrogenase), KRT40 and PNMA2 might be the novel biomarkers for NET.

Modules were isolated from PPI network and analyzed. Increased expression of genes such as CNDP2 [180] and VAV3 [181] were important for pathogenesis of various cancer types, but high expression of these genes might be associated with development of NET. PEPD, BLNK, GPRASP2, GPRASP1 and CEP126 might be the novel biomarkers for NET.

Target gene - miRNA network was constructed and analyzed. MACC1 was linked with invasion of colon cancer cells [182], but this gene might be associated with invasion of NET cells. E2F2 was responsible for proliferation of colon cancer cells [183], but this gene might be important for proliferation of NET cells. Methylation inactivation of tumor suppressor genes such as HEYL [184] and SMOC1 [185] were diagnosed with growth of various cancer types, but loss of these genes might be important for advancement of NET. SULT1B1, SVOP, KCNJ6 and SORCS2 might be the novel biomarkers for NET.

Target gene - TF network was constructed and analyzed.. Target genes such as TBXAS1 [186], HS6ST3 [187] and RFX6 [188] were liable for progression of various cancer types of cancer, but these genes might be linked with progression of NET. RUNX1T1 was associated with pathogenesis of pancreatic endocrine tumors [189], but this may be identified with growth of NET. YIF1B might be the novel biomarker for NET.

## Conclusion

In the current investigation, bioinformatics analyses were used to screen for DEGs between metastatic NET and normal control. Among the 10 hub genes, ATP1A1, LGALS3, LDHA, SYK, VDR, OBSL1, KRT40, WWOX, NINL and PPP2R2B were indicated to be correlated with one another, and those genes were indicated to be able to help predict the prognosis of NET. The results of the current investigation may prove valuable from the perspectives of basic research and clinical treatment for NET. However, even though the current results might contribute to the understanding of the molecular pathogenesis of NET, further investigation are needed to verify genes linked with NET.

## Acknowledgement

We thank Erik Kristiansson, Chalmers University of Technology, Department of Mathematical Sciences, Chalmers Tvärgata 3, Göteborg, Sweden, very much, the author who deposited their microarray dataset, GSE65286, into the public GEO database.

## Author Contributions

Praveenkumar Devarbhavi - Investigation and resources

Basavaraj Vastrad - Writing original draft, and review and editing

Anandkumar Tengli **-** Writing original draft and investigation

Chanabasayya Vastrad - Software and investigation

Iranna Kotturshetti - Supervision and resources

## Availability of data and materials

The datasets supporting the conclusions of this article are available in the GEO (http://www.ncbi.nlm.nih.gov/geo) repository. [(GSE65286) (https://www.ncbi.nlm.nih.gov/geo/query/acc.cgi?acc=GSE65286)]

## Consent for publication

Not applicable.

## Ethical approval

This article does not contain any studies with human participants or animals performed by any of the authors.

## Informed consent

No informed consent because this study does not contain human or animals participants.

